# *Paramecium* Polycomb Repressive Complex 2 physically interacts with the small RNA binding PIWI protein to repress transposable elements

**DOI:** 10.1101/2021.08.12.456067

**Authors:** Caridad Miró Pina, Takayuki Kawaguchi, Olivia Charmant, Audrey Michaud, Isadora Cohen, Adeline Humbert, Yan Jaszczyszyn, Laurence Del Maestro, Daniel Holoch, Slimane Ait-Si-Ali, Olivier Arnaiz, Raphaël Margueron, Sandra Duharcourt

## Abstract

Polycomb Repressive Complex 2 (PRC2) maintains transcriptionally silent genes in a repressed state via deposition of histone H3 K27 trimethyl (me3) marks. PRC2 has also been implicated in silencing transposable elements (TEs) yet how PRC2 is targeted to TEs remains unclear. To address this question, we performed tandem affinity purification combined with mass spectrometry and identified proteins that physically interact with the *Paramecium* Enhancer-of-zeste Ezl1 enzyme, which catalyzes H3K9me3 and H3K27me3 deposition at TEs. We show that the *Paramecium* PRC2 core complex comprises four subunits, each required *in vivo* for catalytic activity. We also identify PRC2 cofactors, including the RNA interference (RNAi) effector Ptiwi09, which are necessary to target H3K9me3 and H3K27me3 to TEs. We find that the physical interaction between PRC2 and the RNAi pathway is mediated by a RING finger protein and that small RNA recruitment of PRC2 to TEs is analogous to the small RNA recruitment of H3K9 methylation SU(VAR)3-9 enzymes.

## INTRODUCTION

Polycomb Repressive Complex 2 (PRC2) is essential for the development of multicellular organisms. It maintains transcriptionally silent genes in a repressed state by trimethylating histone H3 on lysine 27 (H3K27me3) (Cao et al., 2002; Czermin et al., 2002; Kuzmichev et al., 2002; Müller et al., 2002). PRC2 is composed of a catalytic subunit, the SET domain-containing Enhancer-of-zeste enzyme, and two other core subunits, EED and SUZ12. A fourth core subunit, RBAP46/48, is also necessary for full H3K27 methyltransferase activity (Cao and Zhang, 2004; Jiao and Liu, 2015; Montgomery et al., 2005; Pasini et al., 2004). The recruitment of PRC2 and the regulation of its enzymatic activity are tightly controlled by PRC2 cofactors and modulated by the chromatin context (for reviews: (Chittock et al., 2017; Holoch and Margueron, 2017; Schuettengruber et al., 2017)). In addition to its role in the transcriptional repression of protein-coding genes, PRC2 can also target and silence transposable elements (TEs) (Déléris et al., 2021). While TEs are generally repressed by DNA and/or H3K9 methylation (Bourque et al., 2018), H3K27me3 is also associated with TEs in distantly related eukaryotes, including ciliates (Frapporti et al., 2019; Xu et al., 2021), diatoms (Veluchamy et al., 2015), bryophytes (Montgomery et al., 2020) and fungi (Carlier et al., 2021). In ciliates, enhancer-of-zeste enzymes are involved in the repression of TEs (Frapporti et al., 2019; Xu et al., 2021). In plants and animals, H3K27me3 is also found on TEs in mutants and cell types lacking DNA methylation (Déléris et al., 2021). Yet it is currently unknown how PRC2 is recruited to TEs.

In the ciliate *Paramecium tetraurelia*, enrichment of H3K27me3 and H3K9me3 is found at TEs, with a co-occurrence of these two marks on the same TE copies (Frapporti et al., 2019). Deposition of H3K27me3 and H3K9me3 at TEs is catalyzed by the *Paramecium* Enhancer-of-zeste Ezl1 enzyme during development of the somatic MAC (Frapporti et al., 2019). Ciliates have evolved to silence TEs by purging then from the somatic genome during the sexual cycle. Ciliates contain distinct germline and somatic genomes in two types of nuclei within the same cytoplasm. Each *P. tetraurelia* has two transcriptionally inactive germline micronuclei (MIC) and a transcriptionally active somatic macronucleus (MAC). At each sexual generation, the MICs undergo meiosis and, during autogamy, two haploid nuclei fuse to form the zygotic nucleus, which divides twice. The four resulting diploid nuclei develop into two new MICs and two new MACs. The old maternal MAC is fragmented, the fragments being diluted and lost after a few cell divisions. Development of the new functional MAC requires elimination of a large proportion of the germline genome, leading to a highly streamlined somatic genome. In *Paramecium,* the eliminated sequences include TEs and numerous single-copy elements (internal eliminated sequences, IESs) derived from TEs (Arnaiz et al., 2012; Guérin et al., 2017; Sellis et al., 2021). No conserved sequence motif that might serve as a specific recognition signal has been identified among eliminated sequences.

Understanding how such diverse sequences are recognized and excised remains challenging. The process involves small (s)RNA-directed heterochromatin formation (for review (Betermier and Duharcourt, 2014)). 25 nt scanRNAs (scnRNAs), produced from the germline genome during meiosis by Dicer-like proteins, are loaded onto two PIWI proteins (Ptiwi01 and Ptiwi09). Germline-specific scnRNAs are believed to guide the deposition of H3K9me3 and H3K27me3 onto sequences to be eliminated in the developing MAC. Indeed, the scnRNA pathway is required for the deposition of H3K9me3 and H3K27me3 in the developing MAC and for DNA elimination (Lhuillier-Akakpo et al., 2014; Sandoval et al., 2014). However, how deposition of H3K9me3 and H3K27me3 is mediated by PIWI proteins is unknown.

DNA cleavages are introduced at the boundaries of sequences to be eliminated through the action of the PiggyMac (Pgm) endonuclease (Bischerour et al., 2018). In Pgm-depleted cells, in which DNA elimination is inhibited, H3K9me3 and H3K27me3 marks are enriched at TEs (Frapporti et al., 2019). This suggests that scnRNA-guided deposition of H3K27me3 and H3K9me3 marks might promote the tethering of the elimination machinery, leading to the removal of the marked chromatin. Loss of H3K27me3 and H3K9me3 upon Ezl1 depletion leads to retention of TEs in the new developing MAC, TE transcriptional upregulation, and lethality of the sexual progeny (Frapporti et al., 2019; Lhuillier-Akakpo et al., 2014). A second class of developmental-specific small RNAs, the 26-31 nt iesRNAs, which map exclusively to IESs and are bound by two other PIWI proteins (Ptiwi10 and Ptiwi11), appear to facilitate efficient excision (Allen et al., 2017; Sandoval et al., 2014).

Here, we identify the proteins that physically interact with Ezl1 by pulldown and mass spectrometry (MS). We assessed the role of the identified proteins using systematic RNAi-mediated gene knockdown (KD). We found that three Ezl1 interactors, likely functional homologs of the EED, SUZ12 and RBAP46/48 subunits, phenocopied Ezl1 and assembled with Ezl1 into a stable PRC2-Ezl1 core complex in a heterologous expression system. We also identified a number of cofactors that interact with PRC2, including the RNA interference (RNAi) effector and scnRNA-binding protein Ptiwi09, an uncharacterized Ring Finger protein, Rf4, and a protein of unknown function with no conserved domain, Eap1. Rf4 and Eap1 are essential for sexual progeny production, DNA elimination and, in contrast to Ptiwi09, are dispensable for scnRNA biogenesis, suggesting that Rf4 and Eap1 act downstream of scnRNAs. Genome-wide analyses reveal that all cofactors are required for correct targeting of H3K27me3 and H3K9me3 to TEs and their transcriptional repression. We further show that the PRC2-Ezl1 core complex interacts with Ptiwi09 via the Rf4 protein, providing an RNAi-directed mechanism for the specific recruitment of PRC2 to TEs.

## RESULTS

### Identification of Ezl1-interacting partners

To determine how Ezl1 is targeted to specific chromatin sites and how its activity is regulated, we identified its protein partners. Nuclear extracts were prepared from *Paramecium* cells expressing a functional 3xFLAG-HA-tagged Ezl1 during development, when H3K9me3 and H3K27me3 are present in the new developing MAC and DNA elimination occurs (Figure 1A) (Frapporti et al., 2019). Control nuclear extracts were prepared from cells transformed with a plasmid containing the 3xFLAG-HA tag alone under the *EZL1* regulatory sequences (see Materials and Methods). Immunoprecipitation of the FLAG tag and HA tag from the two nuclear extracts, followed by mass spectrometry (n=4, Supplementary Data), recovered Ezl1, as expected, and Caf1 (Ignarski et al., 2014) (Figure 1B and 1C), an ortholog of the PRC2 subunit RBAP46/48/Nurf55 (Figure 1D). Caf1 is known to be required for H3K9me3 and H3K27me3 deposition and DNA elimination (Ignarski et al., 2014). We identified a likely homolog of the human EED, Eed (Figure 1B and 1C), which contains 7 WD40 repeats, and the residues required for the aromatic cage encircling the methyl-lysine (Figure 1D and Supplementary Figure 1), and a likely homolog of mammalian SUZ12, Suz12.like, which has a highly divergent amino-acid sequence and lacks the N-terminal part of SUZ12 , but has a zinc finger and an acidic patch both present in the C-terminal region of the SUZ12 (Figure 1D and Supplementary Figure 2).

**Figure 1.**
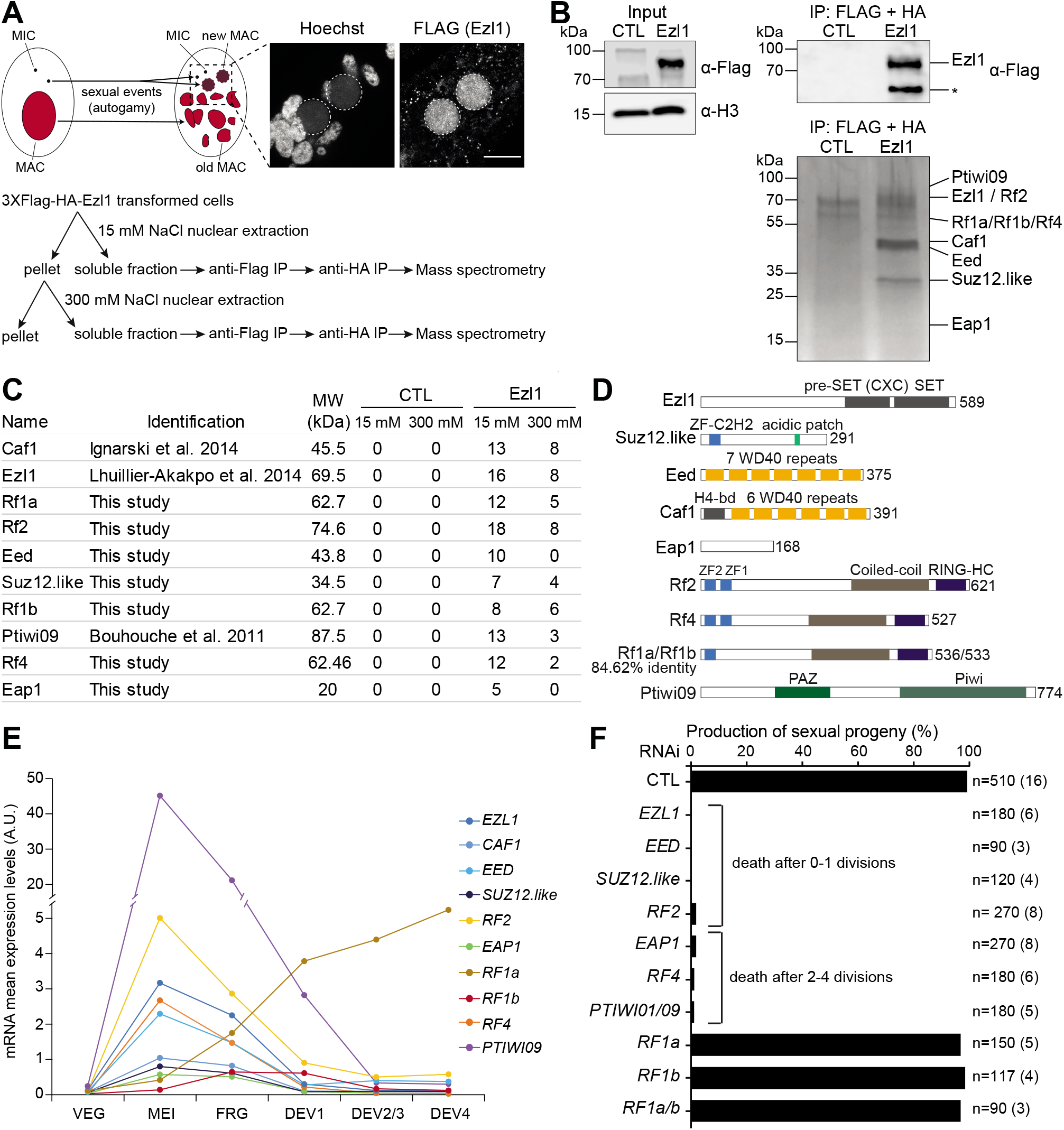
Identification of Ezl1-interacting partners. A. Schematic representation of Ezl1 immunoprecipitation experiments. At each sexual cycle, new MAC and new MIC develop from the zygotic nucleus, while the maternal MAC is fragmented. The confocal image depicts the FLAG fluorescence signal in a *Paramecium* cell expressing a *3xFLAG-HA-EZL1* transgene when nuclear extracts were prepared (T=^~^10 hours after the onset of sexual events). Overlay of Z-projections of magnified views of Hoechst staining and of FLAG-specific antibodies are presented. Dashed white circles indicate the new developing MACs where the Ezl1 fusion protein accumulates at this developmental stage. The other Hoechst-stained nuclei are fragments from the old vegetative MAC. Scale bar is 10 μm. B. Western blot analysis of nuclear extracts from *Paramecium* cells expressing *3XFLAG-HA* (CTL) or *3XFLAG-HA-EZL1* (Ezl1) before (Input) and after double affinity purification (IP). FLAG antibodies were used for 3xFLAG-HA-Ezl1 detection and H3 antibodies for normalization. Silver-stained gel after double affinity purification from the same nuclear extracts. Tandem affinity purified 3xFLAG-HA tagged Ezl1 is known to exhibit catalytic activity towards lysines 9 and 27 of histone H3 (Frapporti et al., 2019). C. Label-free mass spectrometry analysis of 3xFLAG-HA-Ezl1 affinity purification. The table lists the number of unique peptides in one replicate of the 15 mM NaCl and 300 mM NaCl IP experiments for the 10 proteins with the highest fold change in 3xFLAG-HA-Ezl1 IP over the control IP (in all replicates). The number of peptides and the protein coverage in all experiments are provided in Supplementary Data. D. Predicted domains of the identified Ezl1-interacting proteins. E. mRNA expression levels of the genes encoding for Ezl1-interacting proteins at different developmental stages during autogamy in arbitrary units (in thousands) (Arnaiz et al., 2017). F. Production of sexual progeny following RNAi-mediated gene silencing. Two non-overlapping gene segments used for *RF1a* and *RF1b* RNAi (see Materials and Methods) gave the same results. Cells were starved to induce autogamy and, following 2–3 days of starvation, post-autogamous cells were transferred individually to standard growth medium to assess the ability of sexual progeny to resume vegetative growth. The total number of cells analyzed for each RNAi and the number of independent experiments (in parenthesis) are indicated. Cells usually divided two to four times before dying upon *RF4* and *EAP1* KD, as for *PTIWI01/09* KD, and unlike *EZL1*, *EED*, *SUZ12.like*, and *RF2* KD, where cells usually did not divide or only did so once before dying.

Among the Ezl1-associated proteins four, named Rf1a, Rf1b, Rf2 and Rf4, contain a RING-HC finger domain (Figure 1B and 1C). Rf1a and Rf1b are close paralogs (84.62% amino acid identity) (Aury et al., 2006). The four Rf proteins display a similar domain organization, with a C-terminal RING finger domain, a coiled-coil domain, and one or two zinc finger domains at the N-terminus (Figure 1D and Supplementary Figure 3). We identified an uncharacterized small (^~^20 kDa) acidic protein with no conserved domain, referred to as Ezl1-Associated Protein 1 (Eap1) (Figure 1B, 1C and 1D). The scnRNA-binding protein Ptiwi09 (Furrer et al., 2017) was also among the MS hits (Figure 1B and 1C). Although Ptiwi09 and Ptiwi01 are paralogs (98.97% amino acid identity) (Bouhouche et al., 2011) and many MS peptides matching Ptiwi09 also matched Ptiwi01, the only paralog-specific peptides were from Ptiwi09, identifying this protein as the interacting partner (Supplementary Data). Thus, Ezl1-interacting partners include proteins which are likely functional homologs of PRC2 subunit based on domain conservation (Caf1, Eed, Suz12.like) or are putative cofactors having no clear homology with Polycomb proteins, including the RING finger proteins (Rf1a and Rf1b, Rf2 and Rf4), Eap1, and the RNAi effector Ptiwi09.

### Ezl1 partner roles in H3K9me3/H3K27me3 accumulation

RNA-seq expression data (Arnaiz et al., 2017) shows that the *PTIWI09*, *CAF1*, *EED*, *SUZ12.like*, *EAP1*, *RF2*, and *RF4* genes display expression profiles similar to that of *EZL1*, while expression of the *RF1a* and *RF1b* genes are upregulated later during MAC development (Figure 1E). To investigate Ezl1-interacting protein function, we performed KD experiments using RNAi during the sexual cycle of autogamy. Given Ezl1, Caf1, and Ptiwi01/09 are essential for the production of viable sexual progeny (Bouhouche et al., 2011; Ignarski et al., 2014; Lhuillier-Akakpo et al., 2014), we asked if the other Ezl1-interacting proteins were also essential. KD of *EZL1* or *PTIWI01/09* led to lethality of the sexual progeny, while KD of a control, non-essential gene did not impair survival of the post-autogamous progeny (Figure 1F and Supplementary Table 1). KD of *EED* and *SUZ12.like*, *RF2*, *RF4*, and *EAP1* also led to lethality of the sexual progeny (Figure 1F) whereas KD of *RF1A* and *RF1B* did not, either alone or in combination (see Materials and Methods), suggesting they are not essential (Figure 1F). Thus, the Ezl1-interacting proteins that display an mRNA expression profile similar to that of *EZL1* are essential during the sexual cycle, their depletion leading to lethality of the sexual progeny either immediately (Ezl1, Eed, Suz12.like, and Rf2) or after a few divisions (Ptiwi01/09, Eap1, Rf4) (Figure 1F).

Given PRC2 core complex subunits are required for enhancer-of-zeste catalytic activity (Holoch and Margueron, 2017), we examined whether depletion of each Ezl1 partner had an effect on accumulation of H3K9 and H3K27 trimethylation during autogamy, using immunofluorescence and confocal microscopy (Frapporti et al., 2019; Lhuillier-Akakpo et al., 2014). H3K27me3 was detected in the maternal MAC, while H3K9me3 and H3K27me3 accumulated in the new developing MACs in control conditions (Figure 2A), as previously described (Frapporti et al., 2019; Lhuillier-Akakpo et al., 2014). In contrast, we could not detect accumulation of H3K27me3 in the maternal MAC nor of H3K9me3 and H3K27me3 in the new developing MACs in *EED*, *SUZ12.like* or *RF2* KDs (Figure 2A and 2B and Supplementary Figure 4), as previously reported for *EZL1* KD (Frapporti et al., 2019; Lhuillier-Akakpo et al., 2014) and *CAF1* KD (Ignarski et al., 2014). Quantification of H3K9me3 and H3K27me3 fluorescence in the new developing MACs confirmed that H3K9me3 and H3K27me3 accumulated in the new MAC in control cells, while both were significantly diminished in the absence of Ezl1, Eed, Suz12.like or Rf2 (Figure 2B and Supplementary Figure 4) (see Materials and Methods), indicating that Ezl1, Eed, Suz12.like and Rf2 are required for the deposition of H3K9me3 and H3K27me3 in the developing new MAC.

**Figure 2.**
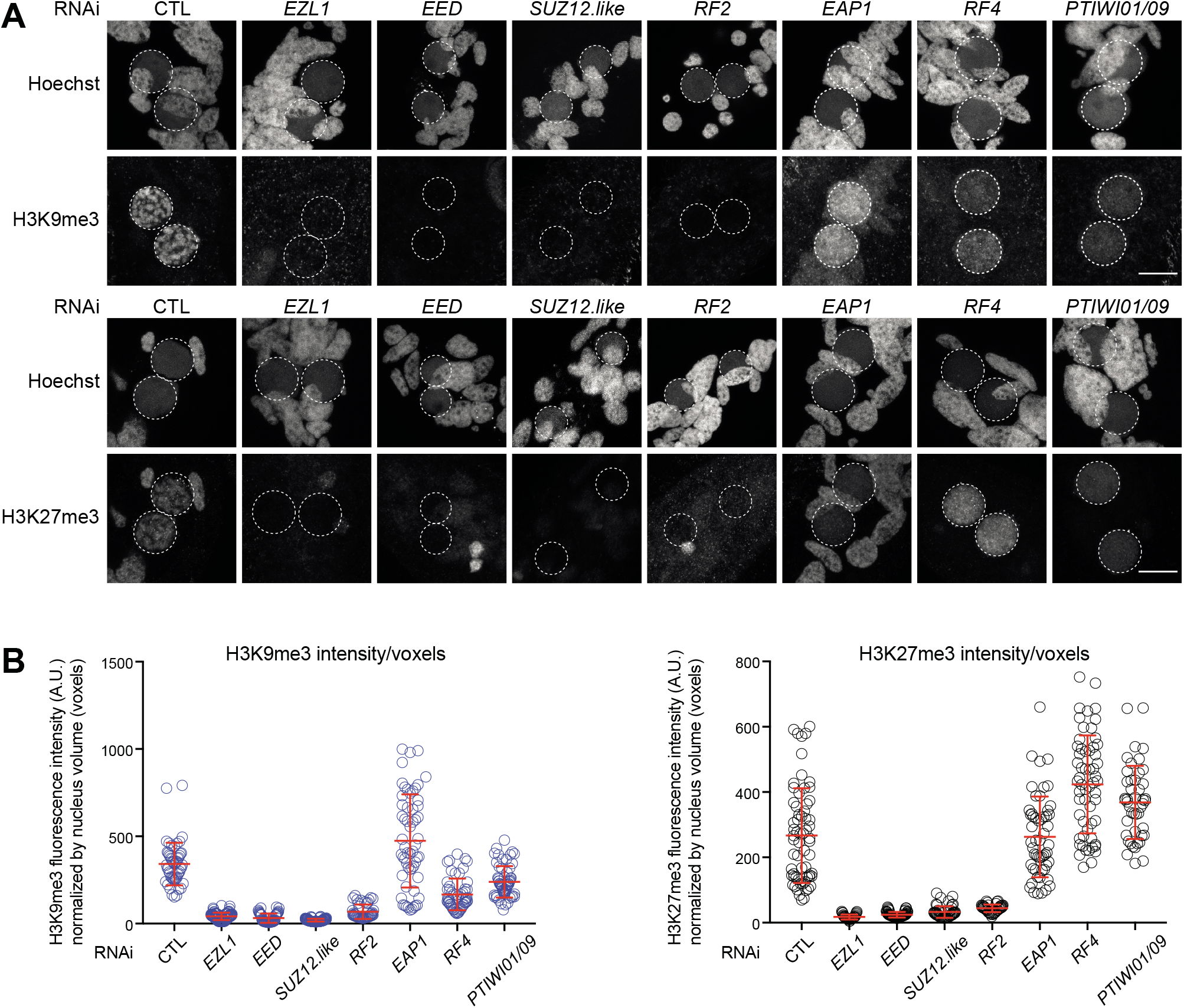
Depletion of Ezl1-interacting partners alters H3K9me3 and H3K27me3 deposition. A. Immunostaining with H3K9me3 (top) or H3K27me3 (bottom) antibodies during autogamy (T=23-T35 after the onset of sexual events). Representative images are displayed. Overlay of Z-projections of magnified views of Hoechst staining and of H3K27me3- or H3K9me3-specific antibodies are presented. Dashed white circles indicate the new developing MACs. The other Hoechst-stained nuclei are fragments from the old maternal vegetative MAC. In control cells, H3K27me3 is detected in the fragments from the old vegetative MAC, while both H3K9me3 and H3K27me3 are detected in the new developing MACs. Scale bar is 10 μm. B. Quantification of H3K9me3 and H3K27me3 fluorescence signal in the developing MAC during autogamy (same populations). Bar plots represent the total H3K9me3 or H3K27me3 fluorescence intensity in the developing new MAC divided by the number of voxels in the different RNAi conditions. Mann-Whitney statistical tests: significant difference with CTL (p< 0,0001) in all RNAi conditions, except for *EAP1*: not significant (H3K27me3), [P-value=0,0065] (H3K9me3).

In contrast, H3K9me3 and H3K27me3 could still be detected in the new MAC upon depletion of the other Ezl1-interacting proteins (Eap1, Rf4 or Ptiwi01/09) (Figure 2B and Supplementary Figure 4). Quantification of H3K9me3 and H3K27me3 fluorescence in the new developing MACs indicated that (i) H3K9me3 accumulation is significantly diminished upon Rf4 or Ptiwi01/09 depletion (Figure 2B) while it is delayed upon Eap1 depletion (Figure 2B and Supplementary Figure 4), and (ii) H3K27me3 accumulation is delayed upon Eap1 or Ptiwi01/09 depletion, and significantly increased upon Rf4 depletion. H3K27me3 did not accumulate in the maternal MAC upon Rf4 and Ptiwi01/09 depletion (Figure 2B and Supplementary Figure 4).

H3K9me3 and H3K27me3 signals in new developing MACs display a diffuse pattern that gradually forms nuclear foci before the signals coalesce into one single nuclear focus and disappear in control RNAi, as previously reported (Figure 2A and Supplementary Figure 4) (Lhuillier-Akakpo et al., 2014). While depletion of Eap1, Rf4, and Ptiwi01/09 does not prevent H3K9me3 and H3K27me3 deposition in developing MACs (Figure 2A), the H3K9me3 and H3K27me3 signals remain diffuse as development proceeds and no foci are detected (Figure 2A). Thus, Eap1, Rf4, and Ptiwi01/09 are not required for H3K9me3 and H3K27me3 deposition but are required for foci formation, as reported for downstream effectors of the DNA elimination pathway (Lhuillier-Akakpo et al., 2014; Vanssay et al., 2020).

Altogether, our data show that *EED*, *SUZ12.like*, and *RF2* KD abolished H3K9me3 and H3K27me3 accumulation, as previously observed for *EZL1* and *CAF1* KD. In contrast, *EAP1*, *RF4*, and *PTIWI01/09* KD did not prevent the accumulation H3K9me3 and H3K27me3; instead, their accumulation in the developing MACs was either decreased, delayed or even increased, and their localization was altered with no formation of nuclear foci.

### Ezl1, Caf1, Eed and Suz12.like form a complex

Ezl1, Caf1, Eed, and Suz12.like display domain similarities with known PRC2 subunits (Figure 1D) and are all essential for the deposition of H3K27me3 and H3K9me3 (Figure 2), suggesting they form the *Paramecium* core PRC2 complex. To test this hypothesis, we assessed whether these four proteins interact with each other, by performing co-immunoprecipitation (co-IP) experiments in Sf9 insect cells co-infected with baculoviruses expressing Flag-Ezl1, Caf1, Eed, and Suz12.like. The cell lysate was subjected to Flag-IP, and the retained proteins were eluted with Flag-peptide and analysed by Coomassie-stained SDS-PAGE (Figure 3A). Caf1, Eed, and Suz12.like were all pulled down together with Ezl1. To determine whether Ezl1 interacts separately with each subunit or whether these proteins form a stable complex altogether, we purified the co-IP on an ion exchanger, loaded the resulting material on a sizing column (Figure 3B), and analysed the elution profile by western blot, using the His-tags to identify each protein. All subunits have a comparable pattern of elution, with a peak around 1 MDa, consistent with the existence of a stable complex. We performed histone methyl transferase assays with these four subunits using oligonucleosomes as substrate, and the mammalian PRC2 as a positive control (Supplementary Figure 5). However, under these conditions, the recombinant complex was not active.

**Figure 3.**
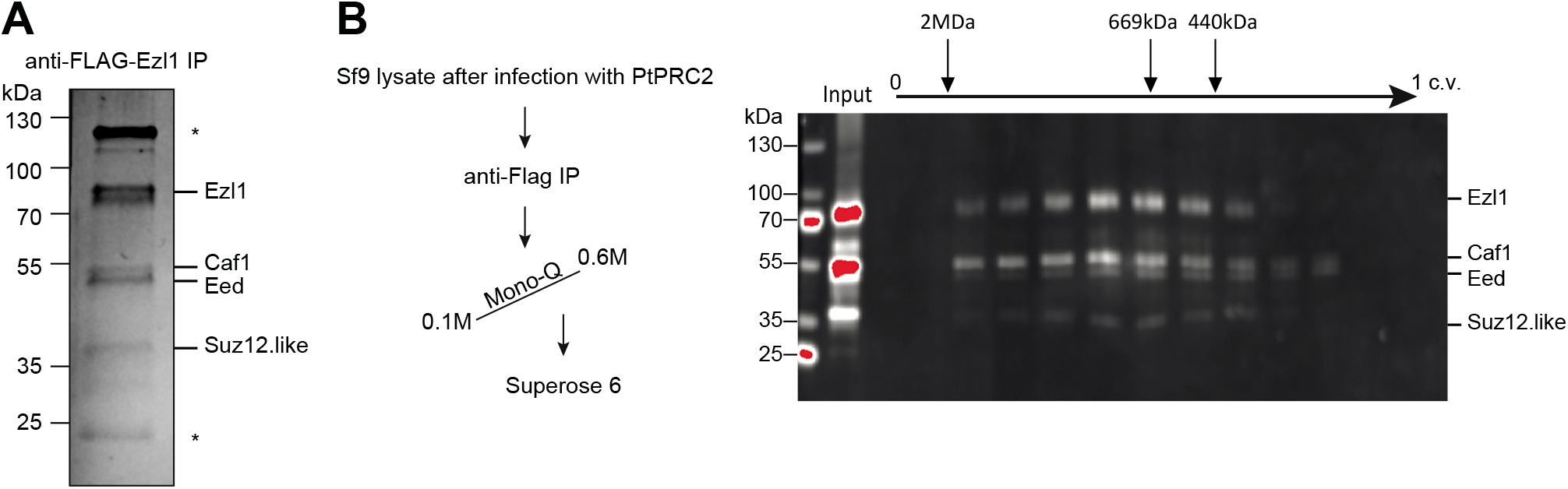
Ezl1, Caf1, Eed and Suz12.like form a complex. A. Coomassie staining of anti-FLAG immunoprecipitation from Sf9 cells co-infected with baculoviruses expressing Flag-Ezl1, Caf1, Eed and Suz12.like (stars = contaminant proteins). B. Left: Purification scheme. Right: Sizing column elution analysis by anti-HIS western blot (all the proteins are HIS-tagged; c.v. = column volume).

Altogether, we show that the *Paramecium* PRC2 core complex consists of Ezl1, Caf1, Eed, and Suz12.like but, in contrast to the metazoan complex, it is not enzymatically active on nucleosomes.

### DNA elimination and TE repression require the core complex

Depletion of Ezl1 or Caf1 impairs DNA elimination (Frapporti et al., 2019; Ignarski et al., 2014; Lhuillier-Akakpo et al., 2014). To analyze the genome-wide effects of depleting *EED* or *SUZ12.like* from the PRC2 core complex, we performed high-throughput sequencing of DNA extracted from a nuclear preparation enriched for new MACs from autogamous cells. We evaluated the effects of *EED* or *SUZ12.like* KD on IES retention by comparison with sequencing data from non-silenced autogamous cells of the same strain (Lhuillier-Akakpo et al., 2014). 70% of IESs are statistically significantly retained in the developing MAC after *EED* or *SUZ12.like* KD (corresponding to 34,607 and 32,964 IESs, respectively, [P-value <0.05]) (Figure 4A), similar to phenotypes observed in *EZL1* and *CAF1* KD (Figure 4B). We analyzed the effects of each depletion on the elimination of MIC-limited sequences other than IESs by comparing the sequencing read coverage of different genomic compartments. The germline genome that is collinear with MAC chromosomes (MAC-destined) is similarly covered in all datasets (Supplementary Figure 6A). The germline-limited (MIC-limited) portion of the genome is not covered by control MAC reads and is well-covered by control MIC reads, as well as by MAC reads from *EZL1*, *CAF1*, *EED*, or *SUZ12.like* KD (Figure 4C), indicating that germline-specific sequences are retained. TEs, which are mostly found in the MIC-limited sequences, are well covered, indicating that TE copies, classified as TIR (n=261), LINE (n=770), Solo-ORF (n=136), and SINE (n= 13) elements (Guérin et al., 2017), are retained upon *EZL1*, *CAF1*, *EED* or *SUZ12.like* KD (Figure 4D).

**Figure 4.**
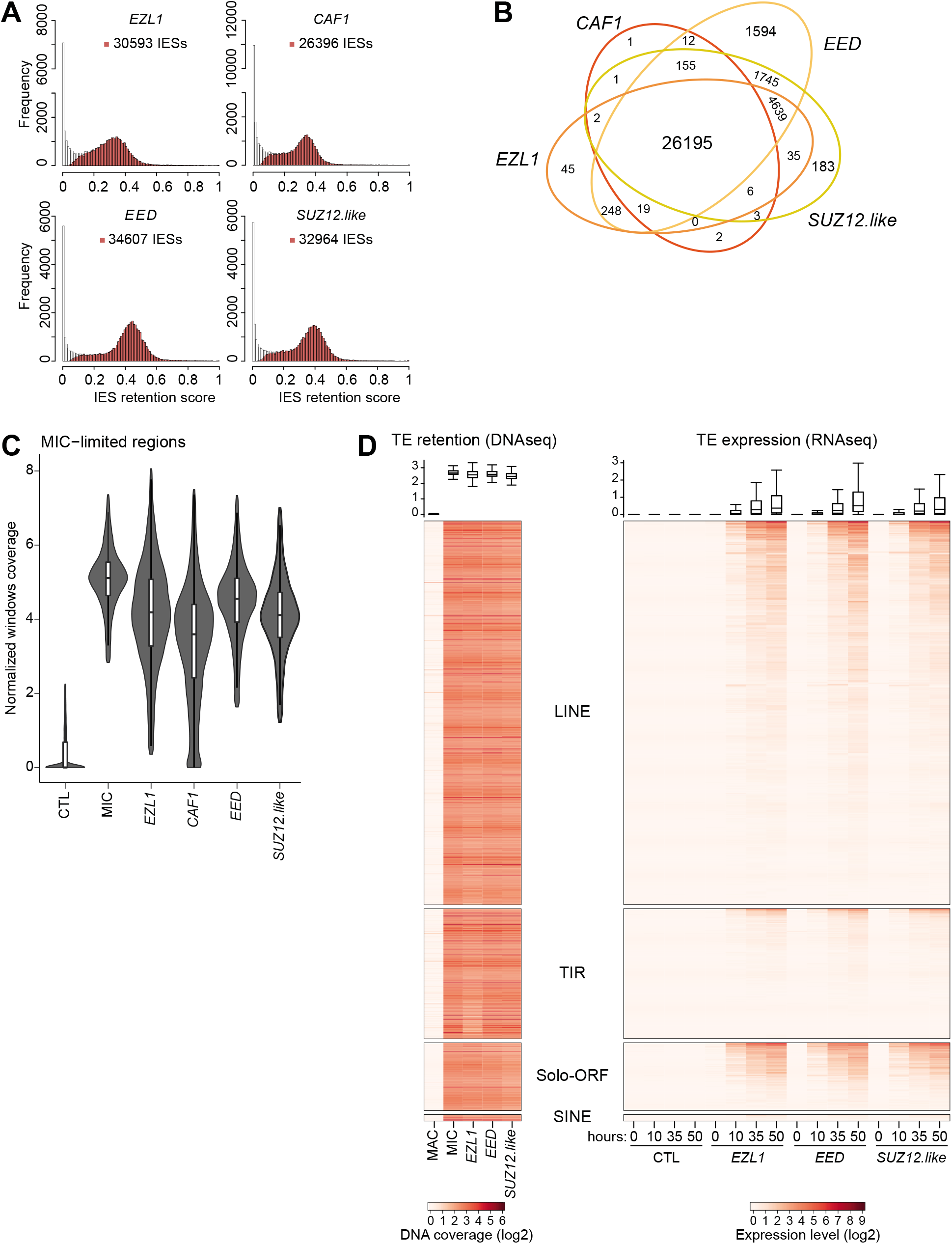
DNA elimination and TE repression require the core complex. A. Histograms of IES retention scores (RS) for *EZL1*, *CAF1*, *EED*, or *SUZ12.like* KD datasets. The distribution of IES RS for significantly retained IESs is shown in red. RS = (IES+) / (IES+ + IES−), where IES+ are the reads that contain an IES end sequence and IES-are the reads that contain the macronuclear IES junction. B. Venn diagram of significantly retained IESs upon the different KDs. C. Violin plot superimposed with a boxplot of normalized coverage of MIC-limited regions using RPKM of 1 kb windows of the MIC assembly for control (CTL), MIC and upon the different KDs. D. Left panel: Heatmap of TE normalized coverage in DNA extracted in MAC, MIC and upon *EZL1*, *EED* or *SUZ12.like* KD. Right panel: Heatmap of RNA expression levels at different time points during development (T=0; T=10; T=35 and T=50) upon the different KDs. In both heatmaps, each row represents a TE copy, and the TE copies are ordered by RNA coverage within each family (LINE, TIR, Solo-ORF, SINE). The boxplots above the heatmaps show the coverage (RPKM log2) for all TE copies (left: DNA coverage, right: RNA coverage).

To determine whether the depletion of each PRC2 core subunit also leads to transcriptional upregulation of TEs, as previously reported upon *EZL1* KD (Frapporti et al., 2019), we performed paired-end RNA-seq at different developmental stages (T=0; T=10; T=35; T=50 hours after the onset of sexual events) (Supplementary Table 2 and Supplementary Figure 7). Comparison of steady-state levels of poly-A RNAs between control, *EZL1*, *EED*, or *SUZ12.like* RNAi was done by mapping RNA-seq reads and counting the fragments onto TE copies (n=1,180) (Materials and Methods), revealing that TE copies are increasingly expressed during MAC development upon *EZL1*, *EED*, *SUZ12.like* RNAi (Figure 4D). The effect is most pronounced late in development (T=50 hours) with ^~^25 % of annotated TE copies, which belong to all four major TE families, being expressed (> 1 RPKM). In contrast, no transcriptional upregulation of TEs was observed upon *PGM* KD (Frapporti et al., 2019), suggesting that retention of TEs in the MAC genome is not sufficient to trigger their transcriptional upregulation. Instead, upregulation of TE transcription can be considered a specific consequence of Ezl1, Eed, and Suz12.like depletion. Thus, depletion of Eed, Suz12.like and Caf1 phenocopy Ezl1 depletion, consistent with these four proteins forming the PRC2-Ezl1 core complex. Their depletion caused the same defects on the elimination of MIC-specific sequences, including the retention of TEs and IESs. Furthermore, like Ezl1, Eed and Suz12.like are required for TE transcriptional silencing.

### Ptiwi09, Eap1 and Rf4 interact with the PRC2-Ezl1 complex

The IP-MS data revealed that Ezl1 interacts with scnRNA-binding Ptiwi09, Eap1, and Rf4 proteins (Figure 1), which are non-essential for Ezl1 catalytic activity (Figure 2). To confirm these interactions, we performed reciprocal immunoprecipitations using functional tagged fusion proteins.

To affinity purify Ptiwi09, *Paramecium* cells expressing a *3XFLAG-HA-PTIWI09* fusion protein, known to bind scnRNAs *in vivo* (Furrer et al., 2017), were used to prepare nuclear extracts during the period when the fusion protein accumulates in the new developing MACs (Figure 5A). Anti-FLAG immunopurification followed by MS (n=2) identified Ptiwi09 peptides (bait), and Ezl1, Suz12.like, Rf2, Rf4, and Eap1 peptides in the Ptiwi09 IP but not in controls (Figure 5A), confirming the interaction between PRC2-Ezl1 and Ptiwi09. The Caf1 and Eed PRC2 core subunits were found in both Ptiwi09 IPs (Supplementary Data). No Rf1a and Rf1b peptides were identified (Figure 5A and Supplementary Data). The top hits (log2 fold change > 1.5, coverage > 8%) included the Ptiwi09 paralogs, Ptiwi01 and Ptiwi03 (98.97% and 58.4% amino acid identity, respectively), (Figure 5A) and the Piwi proteins, Ptiwi10 and Ptiwi11, which share 98.63% amino acid identity and bind iesRNAs, involved in IES excision (Furrer et al., 2017; Sandoval et al., 2014) (Figure 5A). We also identified the putative RNA helicase Ptmb.220 and its paralog (93.21% amino acid identity), which we refer to as Ema1.1 and Ema1.2 (Figure 5D), both being required for DNA elimination and survival of the sexual progeny (Nowak et al., 2011). Its *Tetrahymena* counterpart (Ema1p) is required for scnRNA turnover (Aronica et al., 2008). The uncharacterized protein PTET.51.1.P0270129, which is constitutively expressed during vegetative growth and during the sexual cycle, was also found among the Ptiwi09 interacting partners, and among Eap1 and Rf4 interacting partners (see below) (log2 fold change > 3, coverage > 15% and log2 fold change > 1.5, coverage > 8%, respectively) (Figure 5A). Thus, proteins interacting with Ptiwi09 include known RNAi effectors and the PRC2 core complex, Rf2, Eap1, and Rf4 (Figure 5).

**Figure 5.**
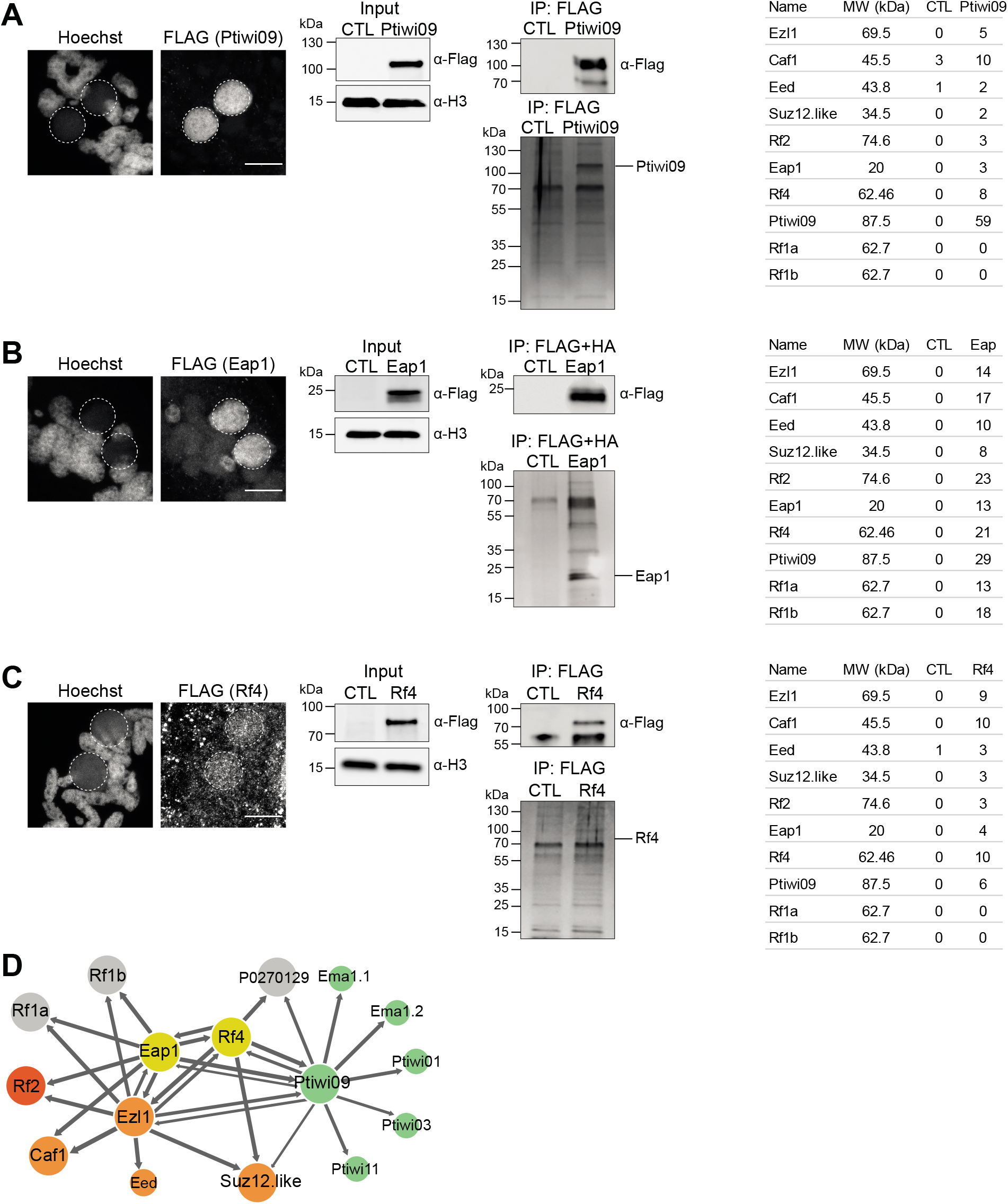
Ptiwi09, Eap1 and Rf4 interact with the PRC2-Ezl1 complex. Immunoprecipitation of Ptiwi09 (A), Eap1 (B) and Rf4 (C) from *Paramecium* nuclear extracts. In each panel, Left: anti-FLAG immunostaining of cells transformed with *3xFLAG-HA* fusion transgenes (for *PTIWI09*, *EAP1* and *RF4*, respectively) at the developmental time when nuclear extracts were performed (T=^~^10 hours after the onset of sexual events). Overlay of Z-projections of magnified views of Hoechst staining and FLAG antibody are presented. Dashed white circles indicate the new developing MACs. The other Hoechst-stained nuclei are fragments from the old vegetative MAC. Scale bar is 10 μm. Middle: Top: Western blot analysis of nuclear extracts from *Paramecium* non-injected cells (CTL) or cells expressing each fusion protein (Ptiwi09, Eap1 and Rf4) and pulled-down proteins from the same nuclear extracts. FLAG antibodies were used for Ptiwi09, Eap1 and Rf4 detection and H3 antibodies for normalization of nuclear extracts. Bottom: Silver-stained gel of pulled-down proteins. Right: Table listing the number of unique peptides identified by label-free mass spectrometry analysis in one replicate of the 10 proteins with a highest fold change in 3xFLAG-HA-Ezl1 IP over the control IP (in all replicates). D. Protein interaction network of Ezl1, Eap1, Rf4 and Ptiwi09-associated proteins. For each IP-MS bait, arrows point to the proteins that are enriched over the control (Ezl1 IPs: proteins with a log2 fold change ≥ 8; Eap1 IPs: proteins with a log2 fold change ≥ 10.5; Rf4 IPs: proteins with a log2 fold change ≥ 8; Ptiwi09 IPs: proteins with a log2 fold change ≥ 7.5 and coverage ≥ 8). For each IP, the width of the arrow depends on the log2 fold change of the protein normalized to the log2 fold change of the bait. The putative *Paramecium* PRC2 core complex is shown in orange, Rf2 in red, Eap1 and Rf4 in yellow and Ptiwi09 and other proteins of the scnRNA pathway in green.

We used a similar strategy to confirm the interaction between Ezl1 and Eap1. Tandem affinity purification of the functional Eap1-3xFLAG-HA fusion protein (Supplementary Figures 8A and 8B) from nuclear extracts prepared when the protein accumulates in the new developing MACs (Figure 5B), followed by MS analysis, identified the top 10 hits found in Ezl1 immunopurification in both Eap1 IP replicates (Figure 5B and Supplementary Data), revealing that Eap1 interacts with the PRC2-Ezl1 core complex, Rf1a and Rf1b, Rf2, Rf4 and Ptiwi09.

Similarly, nuclear extracts prepared when the functional 3xFLAG-HA-Rf4 fusion protein (Supplementary Figures 9A and 9B) accumulates in the new developing MACs (Figure 5C) were used to perform anti-FLAG immunoprecipitations followed by MS analysis (n=2). The PRC2 core subunits were identified, as were Ptiwi09, Eap1, Rf4 and Rf2 (in one replicate), but not Rf1a and Rf1b (Figure 5C and Supplementary Data).

Altogether, these reciprocal immunoprecipitations confirm the interactions between Ptiwi09, Eap1 and Rf4 and the PRC2-Ezl1 core complex.

### The core complex, Eap1 and Rf4 are dispensable for scnRNA biogenesis

Given Ptiwi09, Eap1 and Rf4 depletion yielded similar phenotypes (Figures 1 and 2), we investigated whether Eap1 and Rf4, like Ptiwi01/09, are involved in the scnRNA pathway. To analyze scnRNA levels, we sequenced total small RNA populations in the 15-30 nt range from Eap1-, Rf4-, Suz12.like- and Ezl1-depleted cells and compared them to control and Ptiwi01/09-depleted cells (Furrer et al., 2017) (Supplementary Table 2). sRNA reads were mapped to reference genomes and read counts were normalized to the total number of reads (See Materials and Methods). scnRNAs are produced from the MIC at very early stages of autogamy (T=0 hours after the onset of sexual events). At this time point, 25-nt scnRNAs form the majority of sRNAs in the control sample (Vanssay et al., 2020), as also seen in *EZL1*, *SUZ12.like*, *EAP1* and *RF4* KD cells (Figure 6A), and in contrast to the loss of scnRNAs upon depletion of the scnRNA biogenesis factors Ptwi01/09 (Figure 6A). Thus, Eap1 and Rf4, as well as the PRC2-Ezl1 core complex, are not involved in scnRNA biogenesis, and act downstream of (or in parallel to) scnRNA biogenesis.

**Figure 6.**
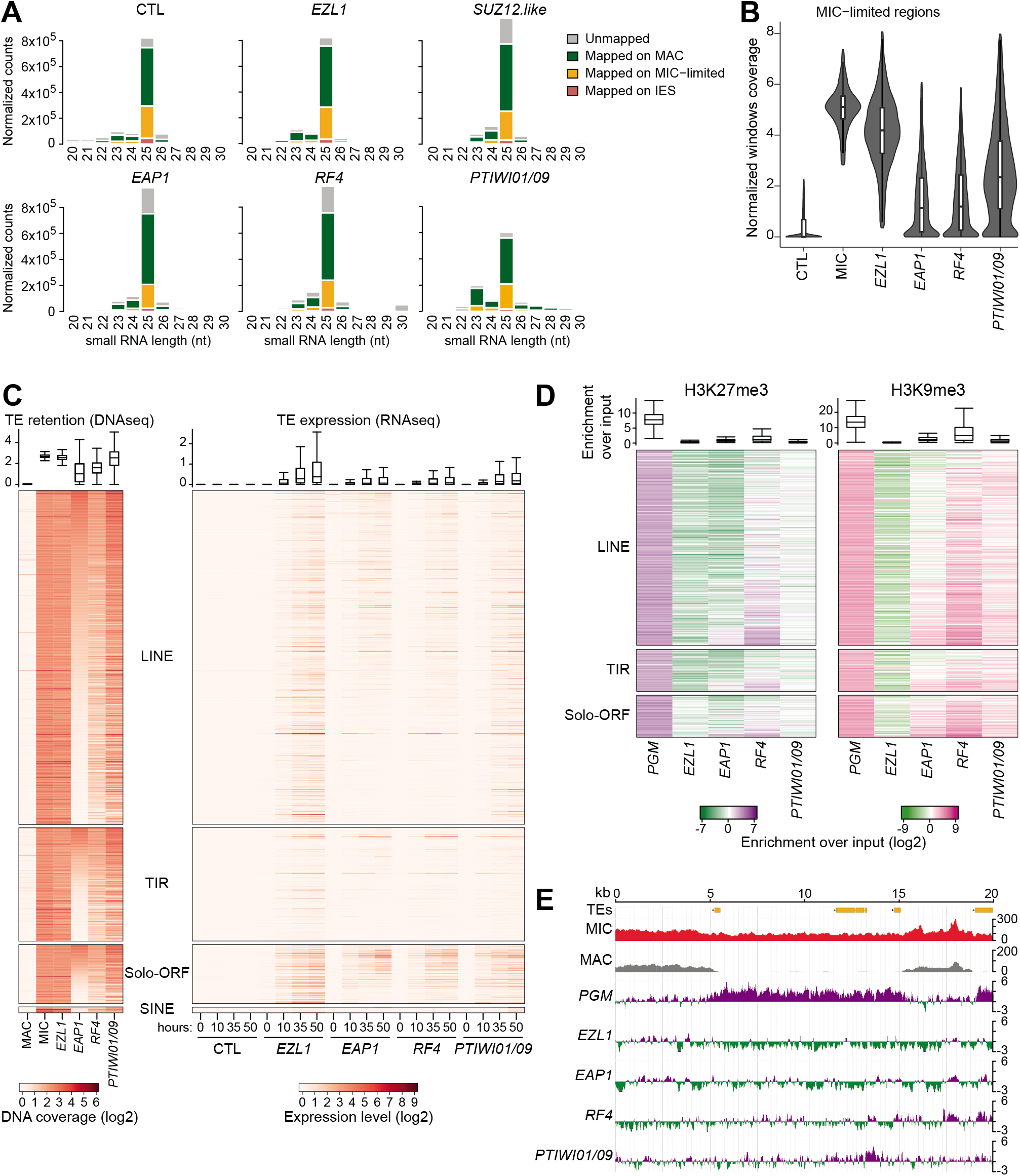
Eap1 and Rf4, like Ptiwi09, are required for DNA elimination, TE silencing, and H3K27me3/H3K9me3 deposition at TEs. A. Analysis of small RNA populations at T=0 hour after the onset of sexual events in different RNAi conditions. Small RNA libraries were sequenced and mapped to the reference genomes. Bar plots show the normalized reads at each sRNA size that are unmapped or match the MAC genome, IESs and MIC-limited genome reference sequences. B. Violin plot superimposed with a boxplot of normalized coverage of MIC-limited regions using RPKM of 1 kb windows of the MIC assembly in control (CTL), MIC and different KD conditions. C. Left panel: Heatmap of TE normalized coverage for MAC, MIC and *EZL1*, *EAP1*, *RF4*, or *PTIWI01/09* KD. Right panel: Heatmap of RNA expression levels at different time points during development upon RNAi-mediated KD. In both heatmaps TE copies are ordered by *EAP1* DNA coverage in each family. The boxplots above the heatmaps show the coverage (RPKM log2) for all TE copies (left: DNA coverage, right: RNA coverage). D. Heatmaps of H3K27me3 and H3K9me3 enrichment over input (log2) for TE copies as determined by ChIP-seq (see Supplementary Figure 11 for the two biological replicates). The TE copies are grouped by TE families (LINE, TIR, Solo-ORF, SINE) and then ordered by *EAP1* DNA coverage. The boxplots show the fold enrichment over the input for the considered TE copies (See Materials and Methods). E. Representative genomic region depicting H3K27me3 at TEs (NODE_1273_length_39901_cov_28.471216 between 20 kb and 40 kb). Lanes show TE annotation, MIC and MAC genome coverages and H3K27me3 normalized enrichment over input (log2) in the different KD conditions.

### Eap1 and Rf4 are required for DNA elimination and TE silencing

Given the PRC2-Ezl1 complex controls DNA elimination, we investigated the impact of Eap1 and Rf4 PRC2 cofactors on DNA elimination. Sequencing of genomic DNA extracted from nuclei preparations enriched for new MACs upon depletion of each protein indicated that only 3.8% and 6.1% IESs [P-value <0.05] are retained upon *EAP1* and *RF4* KD, respectively, similar to Ptiwi01/09 KD (5.7% IESs) (Furrer et al., 2017) (Supplementary Figure 6B). A small subset of IESs depend on the scnRNA pathway, and all require Ezl1 for their excision (Lhuillier-Akakpo et al., 2014). We found that IESs retained in *EAP1*, *RF4* and *PTIWI01/09* KD showed a mild overlap, and almost all these IESs are included in the *EZL1*-dependent subset (Supplementary Figure 6B).

To analyze the effects on the elimination of MIC-limited sequences other than IESs, we calculated the sequencing read coverage of MIC-limited regions in the different KDs and found that these regions are covered upon *EAP1*, *RF4* or *PTIWI01/09* KD, even if it is to a lesser extent than upon *EZL1* KD (Figure 6B and Supplementary Figure 6A). This difference could be due to a lower effect on all MIC-specific regions or to the retention of a subset of MIC-specific sequences. To distinguish these possibilities, we focused on the fraction of MIC-limited regions that comprise annotated TEs (Guérin et al., 2017), and observed that TEs, from all four of the major distinct TE families, are retained in these conditions (Figure 6C), and coverage is once again less than in *EZL1* KD, with the lowest coverage in *EAP1* KD (Figure 6C). Thus, DNA elimination is inhibited upon Eap1, Rf4, and Ptiwi01/09 KD and, as a consequence, TEs are retained in the new MAC.

Given steady-state TE transcript levels are increased upon depletion of the PRC2-Ezl1 core subunits (Figure 4D), we performed transcriptome profiling by RNA-seq at the same developmental stages (T=0; T=10; T=35; T=50 hours after the onset of sexual events) upon *EAP1*, *RF4* or *PTIWI01/09* KD (Supplementary Table 2). As for *EZL1* KD, ^~^15% of annotated TE copies become expressed (>1 RPKM) during MAC development (T=50 hours) upon *PTWI01/09* KD (Figure 6C). TE de-silencing in Eap1- and Rf4 -depleted cells is weaker (with ^~^ 10% of all TE copies expressed >1 RPKM at T50) than in Ptiwi01/09-depleted cells. The patterns of TE derepression are very similar between *EAP1* and *RF4* KD (Figure 6C).

Altogether, we conclude that DNA elimination is inhibited upon *EAP1*, *RF4*, and *PTIWI01/09* KD and, as a consequence, TEs are retained in the new MAC, and their transcript levels increased.

### Ptiwi09, Eap1 and Rf4 are required for H3K9me3 and H3K27me3 deposition at TEs

Depletion of the core PRC2-Ezl1 subunits abolished the deposition of H3K9me3 and H3K27me3, while these two histone marks still accumulated in the new MAC upon depletion of the cofactors Eap1, RF4 and Ptiwi09 (Figure 2). We previously reported that TEs are enriched for H3K9me3 and H3K27me3 in wild-type conditions (at T=10 hours after the onset of sexual events) or upon depletion of the Pgm endonuclease (at T=50 hours after the onset of sexual events) (Frapporti et al., 2019). To determine whether H3K9me3 and H3K27me3 are correctly targeted to TEs, we performed chromatin immunoprecipitation (ChIP) experiments using H3K9me3 and H3K27me3 antibodies (T=50) upon *EAP1*, *RF4* and *PTIWI01/09* KD, and upon *PGM* KD as a positive control. Given that genes, but not TEs, are enriched for H3K4me3 (Frapporti et al., 2019), H3K4me3 ChIP was used as a control. ChIP-qPCR analysis showed that genes (*ACTIN* and *GAPDH*) are enriched in H3K4me3 in all conditions, TEs showed enriched occupancy for both H3K27me3 and H3K9me3 in *PGM* KD conditions while this enrichment was lost upon *EZL1* KD (Supplementary Figure 10). H3K9me3 and H3K27me3 enrichment were reduced for all the analyzed TEs upon *PTIWI01/09* KD, although to a lower extent than in *EZL1* KD conditions. Upon *EAP1* or *RF4* KD, enrichment for H3K9me3 and H3K27me3 was overall lower but it varied among TEs (Supplementary Figure 10).

We extended our analysis by performing ChIP-sequencing. The heterogeneity of DNA coverage among TE copies, especially in *EAP1* KD, prompted us to select the TE copies that are covered (> 10 RPKM) in the inputs for the subsequent analyses. Consistent with previous work (Frapporti et al., 2019), the selected TE copies (251 LINEs, 54 TIRs, 54 Solo-ORFs and 3 SINEs) are enriched in H3K27me3 and H3K9me3 marks with respect to the input in *PGM* KD conditions, while there is no enrichment for H3K4me3 (Figure 6D and Supplementary Figure 11). The patterns are alike for H3K27me3 and H3K9me3 marks, confirming that both histone marks are present on the same TEs. In contrast, *EZL1* KD led to H3K9me3 and H3K27me3 enrichment diminution (Figure 6D and Supplementary Figure 11), as previously reported (Frapporti et al., 2019). Similarly, H3K9me3 and H3K27me3 enrichment are decreased upon *PTIWI01/09* or *EAP1* KD (Figure 6D and Supplementary Figure 11). The decrease in enrichment is weaker and more heterogeneous among TE copies in Rf4-depleted cells than in Ptiwi01/09- or Eap1-depleted cells (Figure 6D). Snapshots of the genome browser illustrate that MIC-limited regions comprising annotated TEs are specifically enriched for H3K27me3 (Figure 6E) and for H3K9me3 (Supplementary Figure 11) upon *PGM* KD and that this enrichment is greatly diminished upon *PTIWI01/09*, *EAP1* and *RF4* KD. Thus, Ptiwi01/09, Eap1 and Rf4 appear to be required for proper H3K9me3 and H3K27me3 targeting.

### Rf4 connects the PRC2-Ezl1 complex to the RNAi effector Ptiwi09

The phenotypes caused by the depletion of Eap1 and RF4 resemble those observed upon Ptiwi01/09 depletion. We therefore reasoned that Eap1 or Rf4 might connect the RNAi effector Ptiwi09 to the *Paramecium* PRC2 complex, allowing its recruitment to MIC-specific regions. To assess this hypothesis, we compared the proteins interacting with Ezl1 in control conditions with those found in Eap1- or Rf4- depleted cells. The localization of the 3xFLAG-HA-Ezl1 fusion protein in either Eap1 or Rf4-depleted cells was confirmed by immunofluorescence (Figure 7A) and by Western blot analysis (Figure 7B). 3xFLAG-HA-Ezl1 IPs from nuclear extracts in the different RNAi conditions were analyzed by MS (Supplementary Data). For each identified protein, we calculated the fold change of total peptides upon *EAP1* or *RF4* RNAi over the control RNAi and adjusted Ezl1 fold change to 1 (Figure 7C). Eap1 fold change is lower in Eap1-depleted cells, while no major decrease in the fold change could be observed for the other Ezl1-interacting partners (Figure 7C) although Rf4 enrichment is higher in *EAP1* RNAi (Figure 7C). Thus, Eap1 depletion does not appear to affect the interaction of the Ezl1 protein with any of its partners. In contrast, in Rf4-depleted cells (Figure 7C), not only the Rf4 fold change is lower but the fold change of Ptiwi09 and of another protein (P02702129) of unknown function, which interacts with both Rf4 and Ptiwi09 (Figure 5D), are decreased. No other major decrease could be observed for the other Ezl1-interacting partners. Therefore, we conclude that Rf4 depletion prevents Ezl1 interaction with Ptiwi09, suggesting that Rf4 bridges the PRC2-Ezl1 complex to Ptiwi09.

**Figure 7.**
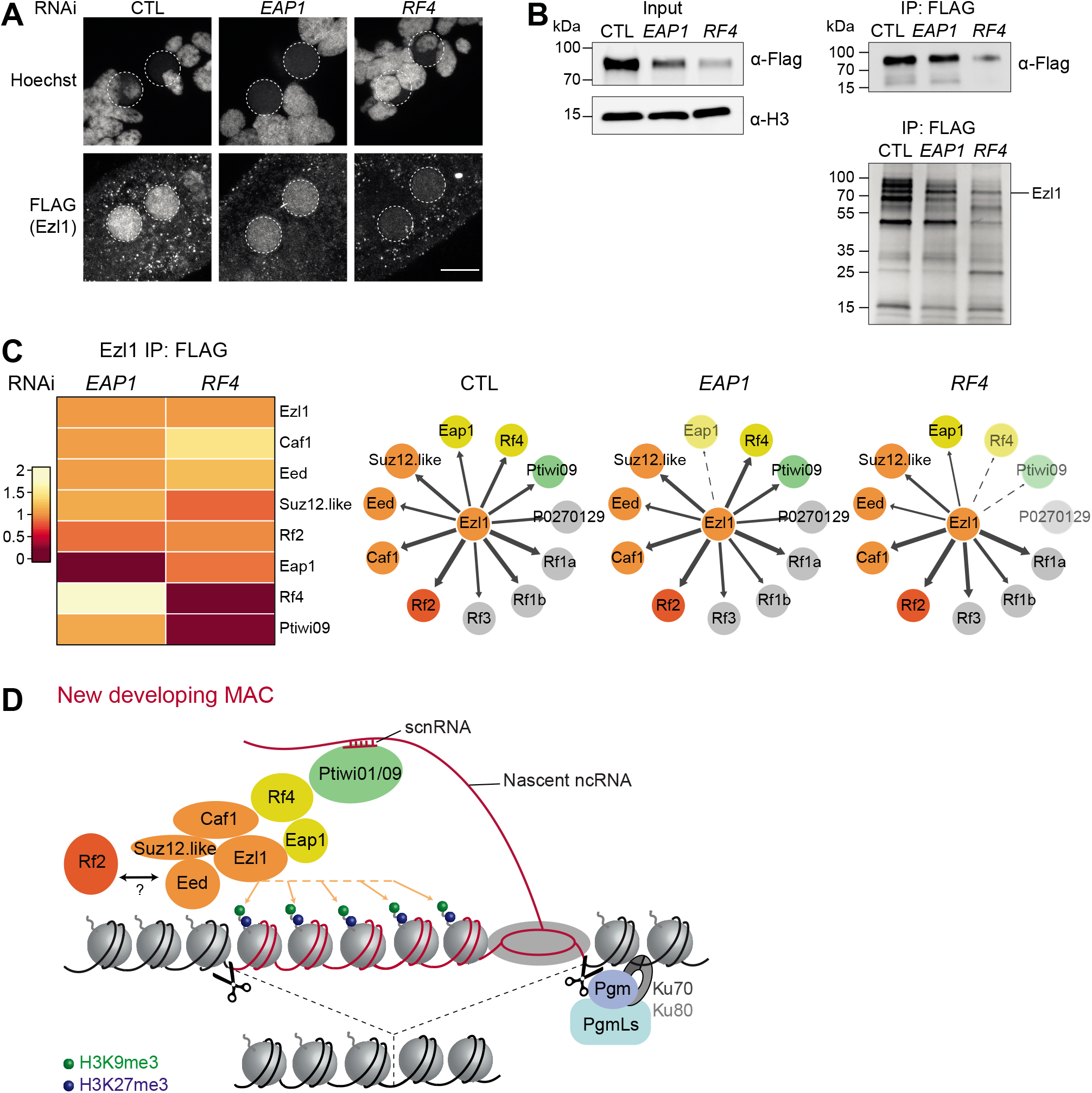
Rf4 connects the PRC2-Ezl1 complex to the RNAi effector Ptiwi09. A. Anti-FLAG immunostaining of cells transformed with a *3xFLAG-HA-EZL1* transgene and silenced for the non-essential gene *ICL7* (CTL), *EAP1* or *RF4* at the developmental time in which nuclear extracts for immunoprecipitation experiments shown in panel B were performed. The Ezl1 fusion proteins accumulated in the new developing MACs, although the fluorescence intensity was reduced, especially in Rf4-depleted cells. Overlay of Z-projections of magnified views of Hoechst staining and FLAG-specific antibody are presented. Dashed white circles indicate the new developing MACs. The other Hoechst-stained nuclei are fragments from the old vegetative MAC. Scale bar is 10 μm. B. Top: Western blot analysis of nuclear extracts (Input) and anti-FLAG pull-down upon *ICL7* (CTL), *EAP1* or *RF4* RNAi. FLAG antibodies were used for 3xFLAG-HA-Ezl1 detection and H3 antibodies for normalization. Bottom: Silver-stained gel of the anti-FLAG immunoprecipitates. C. Left: Heatmap of total peptides fold change in *EAP1* or *RF4* knockdown conditions over control knockdown conditions normalized to Ezl1 fold change. Right: Protein interaction diagrams of Ezl1-associated proteins in *ICL7* (CTL), *EAP1* and *RF4* RNAi conditions. The proteins with a log2 fold change ≥ 3 in the 3xFLAG-HA-Ezl1 control RNAi conditions over the non-transformed control IP are presented. For each IP-MS bait, arrows point to the proteins that are enriched over the control IP (non-transformed cells). The width of the arrows is proportional to the log2 fold change of the protein normalized for Ezl1 log2 fold change in each IP. The dashed arrows correspond to proteins which log2 fold change is at least 2 times lower than in control knockdown conditions. The *Paramecium* PRC2 core complex is shown in orange, Rf2 in red, Eap1 and Rf4 in yellow and Ptiwi09 in green and other proteins in grey. The number of peptides, fold changes and coverage of all proteins in all experiments are provided in Supplementary Data. D. Model for the RNAi-mediated recruitment of PRC2-Ezl1 to TEs. Base pairing interactions between scnRNAs, which are bound to Ptiwi01/09, and nascent ncRNAs produced in the new developing MACs are thought to mediate the specific recognition of germline-specific sequences, including TEs (in red). The RING finger protein Rf4 appears to connect the RNAi effector protein Ptiwi09 to PRC2, which catalyzes the deposition of H3K9me3 and H3K27me3 on MIC-specific sequences, allowing their elimination by the Pgm excision complex.

## DISCUSSION

PRC2-mediated repression of protein-coding genes is well established but its activity at TEs remains poorly characterized. Here, we identified the proteins that interact with Ezl1, the catalytic subunit of *Paramecium* PRC2, which deposits both H3K27me3 and H3K9me3 at TEs. We show that the PRC2 core complex is composed of core subunits, essential for *in vivo* for catalytic activity, with protein domain similarities to known PRC2 subunits. Several cofactors interact with the core complex, including RING-HC-domain-containing proteins (Rf1a, Rf1b, Rf2 and Rf4), a small acidic protein with no conserved domain (Eap1) and the scnRNA-binding protein Ptiwi09. We further show that the core complex is connected to the RNAi pathway via the RING-HC cofactor Rf4, and both the RNAi pathway and Rf4 are essential for correct deposition of H3K9me3 and H3K27me3 at TEs. We thus identify the protein that links the heterochromatin machinery to the nuclear RNAi pathway in *Paramecium* and show that the mechanism of recruitment of PRC2-Ezl1 to TEs resembles that of SU(VAR)3-9 H3K9 methyltransferases.

### The *Paramecium* PRC2-Ezl1 complex

In addition to the enhancer-of-zeste catalytic subunit, mammalian and *Drosophila* PRC2 complexes comprise two other core subunits: EED and SUZ12 (Cao and Zhang, 2004; Jiao and Liu, 2015; Montgomery et al., 2005; Pasini et al., 2004). Being highly divergent, *Paramecium* Eed and Suz12.like had thus far eluded identification on the basis of sequence homology but were identified, among the Ezl1-interacting partners, based on structural similarity to corresponding proteins in other organisms (Figure 1). In *C. elegans* and in the yeast *Cryptococcus neoformans*, no SUZ12 protein has been identified as a PRC2 subunit but the MES-3 and Bnd1 proteins, which appear to be required for catalytic activity, may be functional SUZ12 homologs (Bender et al., 2004; Dumesic et al., 2015). The high sequence divergence of SUZ12 proteins could arise because they act as a binding platform for PRC2 core subunits and accessory proteins, which are not evolutionarily conserved. Caf1, initially characterized for its role in DNA elimination (Ignarski et al., 2014), is a RBAP46/48 homolog and, except in *C. elegans* and *Tetrahymena* (Tursun, 2017; Xu et al., 2021), is found in most PRC2 complexes, likely because of its histone H3-H4 binding function.

The *Paramecium CAF1*, *EED*, *SUZ12.like* and *EZL1* genes display very similar expression profiles (Figure 1) and their depletion leads to identical phenotypes (Figures 1, 2 and 4) (Ignarski et al., 2014; Lhuillier-Akakpo et al., 2014), suggesting that the proteins form a multiprotein complex *in vivo.* Indeed, recombinant Ezl1, Caf1, Eed and Suz12.like proteins co-IP in a heterologous insect system and co-elute on a sizing column, consistent with the formation of a stable complex (Figure 3). However, unlike its mammalian counterpart, the putative recombinant PRC2-Ezl1 core complex did not exhibit *in vitro* methyltransferase activity on nucleosomes (Supplementary Figure 5). The main difference between active endogenous Ezl1 IP (Frapporti et al., 2019) and the inactive recombinant core PRC2-Ezl1 complex is the presence of cofactors, suggesting they are likely required for a fully active complex. Indeed, PRC2 cofactors interact with the core complex in metazoan, modulating its catalytic activity and/or recruitment to target loci (Holoch and Margueron, 2017). In addition to the four *Paramecium* PRC2-Ezl1 core complex subunits, we identified six other Ezl1-interacting proteins, which likely include PRC2 cofactors.

The *Paramecium* PRC2-Ezl1 appears to have a noncanonical composition compared to previously studied PRC2 complexes. Eap1 is a small protein (^~^20 kDa) which is highly acidic and does not harbor domains of known function. Orthologs of Eap1 are not identifiable outside *Paramecium* species. The *EAP1* gene is specifically expressed during sexual events, and its mRNA expression profile is very similar to that of *EZL1* (Figure 1). *EAP1* is an essential gene required for H3K9me3 and H3K27me3 deposition at TEs, acting downstream of scnRNA biogenesis (Figures 2 and 6). How Eap1 triggers histone mark deposition is unclear. The delay we observe in H3K9me3 and H3K27me3 accumulation in the developing MAC upon Eap1 depletion suggests that it stimulates methylation activity (Figure 2 and Supplementary Figure 4). Our ChIP data suggest that Eap1 directly or indirectly recruits the PRC2-Ezl1 complex to target loci (Figure 6), acting to establish or maintain heterochromatin marks.

Using either Ezl1 or Ptiwi09 as a bait (Figures 1 and 5), we show that the scnRNA-binding protein Ptiwi09 physically interacts with the PRC2-Ezl1 core complex. A connection between Ptiwi01/09 and Ezl1 was previously indicated by the requirement of the RNAi pathway for the deposition of H3K9me3/H3K27me3 marks in the developing MAC (Chalker et al., 2013; Coyne et al., 2012; Lhuillier-Akakpo et al., 2014; Xu et al., 2021; Zhao et al., 2019), but no direct physical interaction had yet been reported. The composition of PRC2 appears to differ among evolutionarily distantly related ciliates (Xu et al., 2021). In *Tetrahymena*, the scnRNA-binding protein Twi1 is not found among Ezl1-interacting partners, yet a weak interaction was detected by western blot, under cross-linking conditions, between Ezl1 and the putative RNA helicase Ema1, which is thought to mediate scnRNA-nascent RNA pairing (Aronica et al., 2008; Xu et al., 2021).

Among the set of RING-HC finger domain containing proteins identified as Ezl1-interacting proteins (Rf1a, Rf1b, Rf2, and Rf4), depletion of the Rf2 protein, like the *Tetrahymena* RNF1 and RNF2 proteins, phenocopied Ezl1 depletion (Figures 1, 2 and 4) (Xu et al., 2021). In contrast, depletion of Rf4, which shares the same domain organization and expression profile as Rf2, delayed H3K9me3 accumulation in the developing MAC but did not delay nor prevent H3K27me3 accumulation (Figure 2 and Supplementary Figure 4). The *RF4* KD phenotype suggests that H3K27me3 and H3K9me3 are sequentially deposited in the new developing MACs, and that the Rf4 protein may specifically modulate the H3K9 methylation activity of the PRC2-Ezl1 complex.

RING-HC finger domains are found in RING1/2 and the Polycomb Group Ring Finger (PCGF) protein family, core components of the Polycomb Repressive Complex 1 (Gahan et al., 2020). However, whether Rf2 and Rf4 proteins are homologs of PRC1 components, as proposed for the *Tetrahymena* RNF1 and RNF2 (Xu et al., 2021), is unclear. While the RING1/2 and PCGF of PRC1 share a RING finger domain coupled to a C-terminal RAWUL domain, the Rf2 and Rf4 proteins have a distinct domain organization, with a RING finger domain located at the C-terminus and no identified RAWUL domain (Supplementary Figure 3). The lack of this PRC1 diagnostic signature makes it difficult to ascertain the phylogenetic relationship between the RING-HC proteins identified as Ezl1-interacting partners and PRC1 components. Our results suggest that the substrate specificity of PRC2-Ezl1 can be modulated *in vivo* by the cofactor Rf4, which specifically affects H3K9me3 activity (Figure 2).

### RING-HC Rf4 bridges the RNAi effector to the PRC2-Ezl1 complex

scnRNAs, initially produced from the germline MIC genome during meiosis by a specific RNAi pathway, trigger the elimination of homologous sequences in the developing MAC (Bouhouche et al., 2011; Furrer et al., 2017; Lepère et al., 2009, 2008; Lhuillier-Akakpo et al., 2014; Sandoval et al., 2014). Previous work had shown that scnRNAs are required for the accumulation of H3K9me3 and H3K27me3 in the developing MAC in *Paramecium* (Lhuillier-Akakpo et al., 2014) and in *Tetrahymena* (Liu et al., 2007, 2004). We now demonstrate that depletion of the Ptiwi01/09 proteins, which results in destabilization and reduced accumulation of scnRNAs (Furrer et al., 2017), prevent the deposition of H3K9me3 and H3K27me3 on TEs in the developing MAC (Figure 6). We identify the scnRNA-binding protein Ptiwi09 as a cofactor of the PRC2-Ezl1 complex (Figure 1 and 7). The Ptiwi09 paralogs, Ptiwi01 and Ptiwi03, were also identified in the anti-FLAG Ezl1 pull down but with fewer peptides (Supplementary Data), likely because of the lower expression levels of these genes (Arnaiz et al., 2017). Depletion of the PRC2-Ezl1 cofactor Rf4 specifically disrupts the interaction between Ptiwi09 and Ezl1, while Ezl1 still interacts with the other subunits of the core complex (Figure 7). Accumulation of H3K9me3 and H3K27me3 in the developing MAC upon Rf4 depletion suggests that Rf4 is not required for PRC2-Ezl1 enzymatic activity (Figure 2). Instead, the diminution of H3K9me3 and H3K27me3 marks associated with TEs upon its depletion indicates that Rf4 is essential for correct targeting of the PRC2-Ezl1 complex, downstream from scnRNA production (Figure 6). Our work provides a mechanism for the recruitment of PRC2-Ezl1 to TEs (Figure 7). The physical association with the scnRNA/RNAi machinery would tether the PRC2-Ezl1 complex to specific genomic regions in order to initiate silent chromatin formation. In this mechanism, scnRNAs target the Piwi proteins to complementary nascent transcripts to guide heterochromatin formation, therefore providing specificity, despite the lack of conserved motif on eliminated sequences.

The same principle applies in fungi and in metazoans, where small RNAs guide Piwi proteins to complementary nascent transcripts on chromatin and repress the transcriptional activity of TEs and other repeats (Martienssen and Moazed, 2015). In *Drosophila*, the PIWI-interacting RNA (piRNA) pathway orchestrates the deposition of H3K9 methylation by the SetDB1/Eggless methyltransferase and transcriptional repression at piRNA target loci, by poorly understood mechanisms. A link between Piwi and the SetDB1/Eggless H3K9 methyltransferase was recently reported. The SUMO E3 ligase Su(var)2-10 physically interacts with Piwi and its auxiliary factors, Asterix and Panoramix, and its auto-SUMOylation appears to induce the recruitment of SetDB1/Eggless at TEs (Ninova et al., 2020). In *S. pombe*, the RITS complex, which contains the small RNA binding protein Argonaute Ago1, physically interacts with the methyltransferase complex (CLRC), which catalyzes H3K9 methylation at pericentromeric repeats (Holoch and Moazed, 2015). This interaction is mediated by the Stc1 protein, which has been shown to physically interact with both the RITS and the CLRC complexes and is required for H3K9me3 deposition (Bayne et al., 2010). Stc1 is a small protein, only found in fungi and algae, with a LIM domain, characterized by the presence of tandem zinc-finger domains that are required for the interaction with Ago1 (He et al., 2013). Like Stc1, Rf4 links the RNAi pathway and the heterochromatin machinery. The repression of TEs in eukaryotes involves their transcriptional silencing via targeted chromatin modifications and analogous small RNA guided strategies are employed to recruit methyltransferases to TEs. Instead of H3K9 methyltransferases, ciliates such as *Paramecium* have apparently rewired the RNAi pathway toward a Polycomb enhancer-of-zeste enzyme with dual substrate specificity.

## ACKNOWLEDGMENTS

We wish to thank the members of the Duharcourt lab for fruitful discussions. We thank G. Riddihough (http://lifescienceeditors.com) for comments on the manuscript. This work was supported by the Centre National de la Recherche Scientifique, the Agence Nationale pour la Recherche (ANR) [project “LaMarque” ANR-18-CE12-0005 to SD] and [project “POLYCHROME” ANR-19-CE12-0015 to SD, RM, SA and OA]; the LABEX Who Am I? to SD (ANR-11-LABX-0071; ANR-11-IDEX-0005-02); the Fondation de la Recherche Médicale “Equipe FRM DEQ20160334868” to SD. CMP was recipient of PhD fellowships from Université de Paris and Fondation ARC, and a LABEX Who Am I? transition postdoc fellowship, OC was recipient of a PhD fellowship from Université de Paris and IC was recipient of a postdoctoral fellowship from the LABEX Who Am I? We acknowledge the ImagoSeine facility, member of the FranceBioImaging infrastructure supported by the ANR-10-INSB-04. The sequencing benefited from the facilities and expertise of the high-throughput sequencing platform of I2BC.

## AUTHOR CONTRIBUTIONS

CMP conducted most experiments with the help of OC, IC, AH for cloning and silencing experiments; TK, AM, DH, RM performed experiments with recombinant proteins; YJ made the sequencing libraries; LD, SA prepared the samples for mass spectrometry; OA designed and performed the bioinformatic analyses of NGS data; CMP, TK, DH, RM, SD designed the experiments and CMP, SD wrote the paper with input from all co-authors. SD supervised the project.

## MATERIALS AND METHODS

### *Paramecium* strains, cultivation, and autogamy

All experiments were carried out with the entirely homozygous strain 51 of *P. tetraurelia*. Cells were grown in wheat grass powder (WGP) infusion medium bacterized the day before use with *Klebsiella pneumoniae*, unless otherwise stated, and supplemented with 0.8 mg/mL β-sitosterol. Cultivation and autogamy were carried out at 27 °C as described (Beisson et al., 2010a, 2010b).

### Gene silencing experiments

Plasmids used for T7Pol-driven dsRNA production in silencing experiments were obtained by cloning PCR products from each gene using plasmid L4440 and *Escherichia coli* strain HT115 DE3, as previously described (Galvani and Sperling, 2002). Sequences used for silencing of *ND7*, *ICL7a*, *PGM*, *EZL1*, *EED*, *SUZ12.like*, *RF2*, *RF4*, *EAP1*, *PTIWI01*, *PTIWI09*, *RF1a* and *RF1b* were segments 870-1266 of PTET.51.1.G0050374 (*ND7*); 1-580 of PTET.51.1.G0700039 (*ICL7a*); 873-1439 of PTET.51.1.G0490162 (*PGM*); 989-1501 of PTET.51.1.G1740049 (*EZL1*); 145-444 of PTET.51.1.G0240079 (*EED*); 590-946 of PTET.51.1.G0190277 (*SUZ12.like*); 1514-1916 of PTET.51.1.G1190062 (*RF2*); 10-446 (*RF4#1*) or 623-1038 (*RF4#2*) of PTET.51.1.G0570234; 298-486 of PTET.51.1.G1310069 (*EAP1*); 41-441 of PTET.51.1.G0710112 (*PTIWI01*); 50-439 of PTET.51.1.G0660118 (*PTIWI09*); 18-447 (*RF1a#1*) or 1090-1538 (*RF1a#2*) of PTET.51.1.G1170141 (*RF1a*), 78-426 (*RF1b#1*) or 856-1365 (*RF1b#2*) of PTET.51.1.G0840161 (*RF1b*) and 162-461 (*RF3#1*) or 553-1063 (*RF3#2*) of PTET.51.1.G1280028 (*RF3*). For co-silencing of *RF1a* and *RF1b*, RNAi fragments *RF1a#1* and *RF1b#1* or *RF1a#2* and *RF1b#2* were cloned between the T7 promoters. Preparation of silencing medium and RNAi during autogamy were performed as described in (Baudry et al., 2009). Lethality of post-autogamous cells after RNAi was assessed by transferring 30–60 individual post-autogamous cells to standard growth medium. Cells with a functional new MAC were identified as normally growing survivors unable to undergo a novel round of autogamy if starved after ^~^8 divisions. Cells usually divided two to four times before dying upon *RF4* and *EAP1* KD, as for *PTIWI01/09* KD, and unlike *EZL1*, *EED*, *SUZ12.like*, and *RF2* KD, where cells usually did not divide or only did so once before dying (Figure 1F).

### Transformation with 3xFLAG-HA fusion transgenes

For the construction of in-frame *EAP1-3xFLAG-HA*, *3xFLAG-HA-RF4* and *3xFLAG-HA-PTIWI09* fusion plasmids, a 3xFLAG (DYKDDDDK-DYKDDDDK-DYKDDDDK)—HA (YPYDVPDYA) tag that was codon-optimized for the *P. tetraurelia* genetic code was added by RF cloning to the 5’ (*RF4*, *PTIWI09*) or 3’ (*EAP1*) ends of each gene. As a result, the 3xFLAG-HA is fused to the N-terminus or C-terminus of each gene and the fusion proteins are expressed under the control of their transcription signals (promoter and 3’UTR). The *EAP1* transcription signals contains 193-bp upstream and 508-bp downstream of its open reading frame. The *RF4* transcription signals contains 202-bp upstream and 199-bp downstream of its open reading frame. The *PTIWI09* transcription signals contain 514-bp upstream and 235-bp downstream of its open reading frame. The *EAP1-3xFLAG-HA* and *3xFLAG-HA-RF4* fusion transgenes were RNAi-resistant (the *3xFLAG-HA-RF4* construct is resistant to the RNAi fragment #2). The DNA fragments 571-1089 for *RF4* and 298-504 for *EAP1* coding sequences were replaced by RF cloning with synthetic DNA sequences (Eurofins Genomics) designed to maximize nucleotide sequence divergence with the endogenous genomic loci without modifying the amino acid sequences of the encoded proteins. The *3xFLAG-HA-EZL1* plasmid was described in (Frapporti et al., 2019) and the *3xFLAG-HA* plasmid contains the tag alone under the *EZL1* regulatory sequences (844-bp upstream and 308-bp downstream the *EZL1* open reading frame). Plasmids carrying *3xFLAG-HA*, *3xFLAG-HA-EZL1*, *EAP1-3xFLAG-HA*, *3xFLAG-HA-RF4* or *3xFLAG-HA-PTIWI09* transgenes were linearized by XmnI (*3xFLAG-HA* and *3xFLAG-HA-EZL1*), BglII (*3xFLAG-HA-RF4* and *EAP1-3xFLAG-HA*) or BseYI (*3xFLAG-HA-PTIWI09*) and microinjected into the MAC of vegetative 51 cells in the macronucleus. No lethality was observed in the post-autogamous progeny of injected cells, indicating that the none of the fusion constructs interfered with the normal progression of autogamy.

### Nuclear extraction, immunoprecipitation and Western blot

Nuclear extracts were performed as previously described (Frapporti et al., 2019). Autogamous cells (^~^T=10 hours) were lysed with a Potter-Elvehjem homogenizer in lysis buffer (10 mM Tris pH 6.8, 10 mM MgCl2, 0.2% Nonidet P-40, 1 mM PMSF (Sigma-Aldrich), 4 mM benzamidine (Sigma-Aldrich), 1x Complete EDTA-free Protease Inhibitor Cocktail tablets (Roche)). Following the addition of 2.5 volumes of washing solution (0.25M sucrose, 10 mM MgCl2, 10 mM Tris pH 7.4, 1 mM PMSF (Sigma-Aldrich), 4 mM benzamidine (Sigma-Aldrich), 1x Complete EDTA-free Protease Inhibitor Cocktail tablets (Roche)), the nuclei-containing pellet was collected by centrifugation and incubated in 1 volume of nuclear extraction buffer 2 × (100 mM Hepes pH 7.8, 100 mM KCl, 30 mM or 300 mM NaCl, 0.2 mM EDTA, 20% Glycerol, 2 mM DTT, 0.02% Nonidet P-40, 2 mM PMSF, 2x Complete EDTA-free Protease Inhibitor Cocktail tablets (Roche)) at 4 °C for 1 hour. The salt-extractable fraction was recovered following centrifugation at 10,000 g at 4°C for 3 minutes. For 3xFLAG-HA-Ezl1 tandem affinity immunoprecipitation, after 15mM NaCl nuclear extraction, the pellet was incubated for 1 hour in 1 volume of 600 mM NaCl nuclear extraction buffer 2 × and centrifuged at 10,000 g at 4°C for 3 minutes. For anti-FLAG immunoprecipitation, nuclear extracts were incubated overnight at 4°C with anti-FLAG M2 magnetic beads (Sigma #M8823) or anti-FLAG M2 agarose gel (Sigma #A2220). Beads were washed five times with TEGN buffer (20 mM Tris pH 8, 0.1 mM EDTA, 10% Glycerol, 150 mM NaCl, 0.01% Nonidet P-40) and eluted with 3xFLAG peptide (Sigma #F4799). For tandem immunoprecipitation the eluate was further affinity-purified on anti-HA antibody-conjugated agarose (Sigma# A2095) overnight at 4°C and eluted with the HA peptide (Sigma #I2149) overnight at 4°C. For Western blot, electrophoresis and blotting were carried out according to standard procedures. FLAG (1:1000, ThermoFisher Scientific #MA1-91878) and H3 (1:10,000) (Frapporti et al., 2019) primary antibodies were used. Secondary horseradish peroxidase-conjugated anti-mouse or anti-rabbit rabbit IgG antibody (Promega #W4021 and #W4011) were used at 1:2,500 dilution followed by detection by ECL (SuperSignal West Pico Chemiluminescent Substrate, Thermo Scientific).

### Mass spectrometry

The purified immunocomplexes were boiled during 5 min at 96°C in NuPAGE 4X loading buffer (Life Technologies) and NuPAGE™ Sample Reducing Agent (Life Technologies). Samples were then loaded on an SDS-polyacrylamide 4-12 % gel and run at 150 V constant for 5 min to allow the proteins enter the gel. The gel was washed with water; fixed for 20 – 30 min with fixative solution (10 % acetic acid, 50 % methanol, 40 % ultra-clean water) then washed 5 times in ultra-clean water. The bands containing the proteins were cut, stored in 1 mL of ultra-clean water and sent to mass spectrometry (MS) facility. MS identification of proteins was carried out in the Taplin Mass Spectrometry Facility (Harvard Medical School, Boston, MA, USA. https://taplin.med.harvard.edu/home).

Briefly, excised gel bands were cut into approximately 1 mm^3^ pieces. Gel pieces were then subjected to a modified in-gel trypsin digestion procedure (Shevchenko et al., 1996). Gel pieces were washed and dehydrated with acetonitrile for 10 min. After removal of acetonitrile, gel pieces were then completely dried in a speed-vac, then rehydrated in 50 mM ammonium bicarbonate solution containing 12.5 ng/μl modified sequencing-grade trypsin (Promega, Madison, WI) at 4°C. After 45 min, the excess of trypsin was removed and replaced with 50 mM ammonium bicarbonate solution to cover the gel pieces overnight at 37°C. Peptides were extracted by removing the ammonium bicarbonate solution, followed by one wash with a solution containing 50% acetonitrile and 1% formic acid. The samples were then dried in a speed-vac (^~^1 hr) and stored at 4°C. For the MS analysis the samples were reconstituted in 5-10 μl of HPLC solvent A (2.5% acetonitrile, 0.1% formic acid). A nano-scale reverse-phase HPLC capillary column was created by packing 2.6 μm C18 spherical silica beads into a fused silica capillary (100 μm inner diameter x ^~^30 cm length) with a flame-drawn tip (Peng and Gygi, 2001). After equilibrating the column, each sample was loaded via a Famos auto sampler (LC Packings, San Francisco CA) onto the column. A gradient was formed, and peptides were eluted with increasing concentrations of solvent B (97.5% acetonitrile, 0.1% formic acid). After elution, peptides were subjected to electrospray ionization and then entered an LTQ Orbitrap Velos Pro ion-trap mass spectrometer (Thermo Fisher Scientific, Waltham, MA). Peptides were detected, isolated and fragmented to produce a tandem mass spectrum of specific fragment ions for each peptide. Peptide sequences, and hence protein identity, were determined by matching the protein database (*Paramecium tetraurelia* strain d4-2 MAC genome (v1.0)) with the acquired fragmentation pattern by the software program Sequest (Thermo Fisher Scientific, Waltham, MA) (Eng et al., 1994). All databases include a reversed version of all the sequences and the data were filtered to between a one and two percent peptide false discovery rate.

For the proteins Ptiwi01, Ptiwi03 and Ptiwi09 and Rf1a and Rf1b that are paralogs with high amino acid identity, common peptides were assigned to either one, proportionally to the unique peptides identified for each one.

### Indirect immunofluorescence and quantification

Cells were fixed as previously described (Frapporti et al., 2019). Fixed cells were incubated overnight at room temperature with primary antibodies as follows: rabbit anti-H3K9me3 (1:200), rabbit anti-H3K27me3 (1:1000) and mouse anti-FLAG (1:200; Thermo Fisher Scientific #MA1-91878). Cells were labeled with Alexa Fluor 568-conjugated goat anti-rabbit IgG (Invitrogen #A-11036), Alexa Fluor 488-conjugated goat anti-rabbit IgG (Invitrogen #A-11034) or Alexa Fluor 568-conjugated goat anti-mouse IgG (Invitrogen #A-11031) at 1:500 for 1 h, stained with 1 μg/mL Hoechst for 5–10 min and finally mounted in Citifluor AF2 glycerol solution (Citifluor Ltd, London). Images were acquired using a Zeiss LSM 780 or 710 laser-scanning confocal microscope and a Plan-Apochromat 63 × /1.40 oil DIC M27 objective. Z-series were performed with Z-steps of 0.35 μm. Quantification was performed as previously described (Vanssay et al., 2020) using ImageJ. For each cell, the volume of the nucleus (in voxels) was estimated as follows: using the Hoechst channel, the top and bottom Z stacks of the developing MAC were defined to estimate nucleus height in pixels. The equatorial Z stack of the developing MAC was defined, and the corresponding developing MAC surface was measured in pixels. The estimated volume of the developing MAC was then calculated as the product of the obtained nucleus height by the median surface. For each Z stack of the developing MAC, the H3K9me3 or H3K27me3 fluorescence intensity was measured and corrected using the ImageJ “subtract background” tool. The sum of the corrected H3K9me3 or H3K27me3 fluorescence intensities for all the Z stacks, which corresponds to the total H3K9me3 or H3K27me3 fluorescence intensity, was divided by the estimated volume to obtain the H3K9me3 or H3K27me3 fluorescence intensity per voxel in each nucleus.

### Recombinant protein Production and HMT assays

All subunits of ptPRC2 were cloned in pFastBac through regular cloning strategy. Sf9 cells were grown in Insect XPRESS (Lonza) supplemented with 2%FBS, Penn/Strep and fungizone. Infected cells were lysed in BC300 (25 mM Tris pH 7.9, 1 mM EDTA, 10% glycerol, 300 mM KCl, 0.1 mM PMSF) and sonicated and supernatants were incubated on M2-beads (A2220, Sigma) overnight at 4°C. Beads were washed 3 times with BC300 and eluted in BC300 + 0.2 mg/ml Flag-peptide. For Mono-Q (PC 1.6/5), supernatant was dialyzed against BC100, loaded in the same buffer and eluted by a gradient from BC100 to BC600 (20 c.v.). Elutions were concentrated on Amicon Ultra-15 (Millipore) before loading on the sizing column (Superose 6 increase 3.2/300) equilibrated in BC300. All purification steps were monitored either by Coomassie-stained SDS-PAGE or anti-HIS western blot (Genscript #A00186). HMT assays were essentially performed as described in (Frapporti et al., 2019). Nucleosomes were reconstituted by conventional salt dialysis. We used nucleosomes reconstituted with histones of *Xenopus* sequence, having previously shown that endogenous PtPRC2 is active on this substrate (Frapporti et al., 2019).

### DNA extraction and sequencing

DNA for deep-sequencing was isolated from post-autogamous cells (day 2 of autogamy) as previously described (Arnaiz et al., 2012). Briefly, cells were lysed with a Potter-Elvehjem homogenizer in lysis buffer (0.25 M sucrose, 10 mM MgCl2, 10 mM Tris pH 6.8, 0.2% Nonidet P-40). The nuclei-containing pellet was washed with washing buffer (0.25 M sucrose, 10 mM MgCl2, 10 mM Tris pH 7.4), loaded on top of a 3-mL sucrose layer (2.1 M sucrose, 10 mM MgCl2, 10 mM Tris pH 7.4) and centrifuged in a swinging rotor for 1 hr at 210,000 g. The nuclear pellet was collected and diluted in 200 μl of washing buffer prior to addition of three volumes of proteinase K buffer (0.5 M EDTA pH 9, 1% N-lauryl sarcosine sodium, 1% SDS, 1 mg/mL proteinase K). Following overnight incubation at 55°C, genomic DNA was purified and treated with RNAse. Genomic DNA libraries were prepared with the Westburg NGS DNA Library prep kit, according to the manufacturer recommendations. The quality of the final libraries was assessed with an Agilent Bioanalyzer, using an Agilent High Sensitivity DNA Kit. Libraries were pooled in equimolar proportions and sequenced using paired-end 2×75 pb runs, on an Illumina NextSeq500 instrument, using NextSeq 500 High 150 cycles kit.

### RNA extraction and sequencing

RNA samples were extracted from 200–400-mL cultures at 2,000–4,000 cells/mL as previously described (Frapporti et al., 2019) at different time-points during autogamy (T=0; T=10; T=35; T=50 hours after the onset of sexual events). Total RNA quality was assessed with an Agilent Bioanalyzer 2100, using RNA 6000 pico kit (Agilent Technologies). Directional polyA RNA-Seq libraries were constructed using the TruSeq Stranded mRNA library prep kit (Illumina), following the manufacturer’s instructions. The quality of the final libraries was assessed with an Agilent Bioanalyzer, using an Agilent High Sensitivity DNA Kit. Libraries were pooled in equimolar proportions and sequenced using paired-end 2×75 pb runs, on an Illumina NextSeq500 instrument, using NextSeq 500 High Output 150 cycles kit. Small RNAs of 15-30 nt were purified from total RNA on a 15% TBU/Urea gel. Small RNA libraries were constructed using the NEBNext Small RNA kit according to the manufacturer recommendations. The quality of the final libraries was assessed with an Agilent Bioanalyzer, using an Agilent High Sensitivity DNA Kit. Libraries were pooled in equimolar proportions and sequenced using a single read 75 pb run on an Illumina NextSeq500 instrument, using NextSeq 500 High Output 75 cycles kit.

### Chromatin immunoprecipitation and sequencing

In all, 5-8 × 10^6^ autogamous cells (T=50 hours) were collected by centrifugation and cross-linked with 1% methanol-free formaldehyde (Thermo Fisher Scientific) PHEM 1 × (10 mM EGTA, 25 mM HEPES, 2 mM MgCl_2_, 60 mM PIPES pH 6.9) for 10 min at room temperature. After quenching with 0.125 M glycine, cells were washed and pelleted. Cells were then lysed with a Potter-Elvehjem homogenizer in lysis buffer (0.25 M sucrose, 10 mM Tris pH 6.8, 10 mM MgCl_2_, 0.2% Nonidet P-40, 1 mM PMSF (Sigma-Aldrich), 4 mM benzamidine (Sigma-Aldrich); 1x Complete EDTA-free Protease Inhibitor Cocktail tablets (Roche)). Following the addition of 2.5 volumes of washing solution (0.25 M sucrose, 10 mM MgCl_2_, 10 mM Tris pH 7.4, 1 mM PMSF (Sigma-Aldrich), 4 mM benzamidine (Sigma-Aldrich), 1x Complete EDTA-free Protease Inhibitor Cocktail tablets (Roche)), the nuclei-containing pellet was collected by centrifugation and resuspended at 2 × 10^6^ cells/mL in RIPA buffer without sodium dodecyl sulfate (SDS) (50 mM Tris pH 8, 150 mM NaCl, 10 mM EDTA, 1% Nonidet P-40, 0.5% sodium deoxycholate, 1 mM PMSF (Sigma-Aldrich), 4 mM benzamidine (Sigma-Aldrich), 1x Complete EDTA-free Protease Inhibitor Cocktail tablets (Roche)), flash-freezed in liquid nitrogen as 200 μL aliquots and stored at −80 °C. After addition of SDS to a final concentration of 1%, chromatin was sonicated with Bioruptor (Diagenode) to reach a fragment size of 200-500 bp. DNA concentration was quantified with the Qubit dsDNA BR Assay Kit (Thermo Fisher Scientific). Chromatin corresponding to 13 μg of DNA was diluted 10-fold in RIPA buffer without SDS complemented with protease inhibitors and incubated overnight at 4°C with 4 μg of H3K9me3 or H3K27me3 antibodies (Frapporti et al., 2019) or with 4 μL of H3K4me3 monoclonal antibody (Merck Millipore #04-745). As input control, 10% of sheared chromatin was taken aside. Antibody-bound chromatin was recovered after incubation for ^~^7 h at 4°C with Magna ChIP Protein A + G Magnetic Beads (16–663, Millipore). Beads were washed twice with Low salt buffer (0.1% SDS, 1% Triton, 2 mM EDTA, 20 mM Tris pH 8, 150 mM NaCl), twice with High salt buffer (0.1% SDS, 1% Triton, 2 mM EDTA, 20 mM Tris pH 8, 500 mM NaCl), once with LiCl wash buffer (10 mM Tris pH 8.0, 1% sodium deoxycholate, 1% Nonidet P-40, 250 mM LiCl, 1 mM EDTA) and once with TE 50 mM NaCl (50 mM Tris pH 8.0, 10 mM EDTA, 50 mM NaCl). Chromatin was eluted with TE 1% SDS (50 mM Tris pH 8.0, 10 mM EDTA, 1% SDS) at 65°C for 45 min. ChIP-enriched samples and inputs were then reverse crosslinked at 65°C overnight and treated with RNase A and proteinase K. DNA was extracted with phenol, precipitated with glycogen in sodium acetate and ethanol and resuspended in deionized distilled water. Enrichment compared to input was analyzed by qPCR. qPCR was performed using the ABsolute QPCR SYBR Green Capillary Mix (Thermo Fisher Scientific) on the LightCycler 1.5 thermal cycling system (Roche). qPCR primers were previously published (Frapporti et al., 2019). Approximately 5 ng of DNA were used for ChIP-seq library construction. For *EZL1* RNAi ChIP samples, DNA fragments were end repaired and dA-tailed (NEB #E7595), Illumina TruSeq adapters were ligated (NEB #E6040) and libraries were amplified with Kapa Hifi polymerase (Kapa Biosystem #KK2103). For the other samples, the Microplex Library Preparation kit V2 (Diagenode) was used for ChIP-Seq library construction following the manufacturer recommendations. Final libraries quality was assessed on an Agilent Bioanalyzer 2100, using an Agilent High Sensitivity DNA Kit. Libraries were pooled and sequenced on an Illumina NextSeq500 instrument, according to the manufacturer recommendations, in a Paired-End 2 × 75 pb run (NextSeq500/ 550 Kit v2 (150 cycles)). Demultiplexing was performed with bcl2fastq2 v2.18.12. Adapters were trimmed with Cutadapt v1.15, and only reads longer than 10pb were kept for further analysis.

### Reference genomes and datasets

The sequencing data were mapped on *Paramecium tetraurelia* strain 51 MAC v1 (ptetraurelia_mac_51.fa) and MAC+IES v1 (ptetraurelia_mac_51_with_ies.fa) reference genomes (Arnaiz et al., 2012). The gene annotation v2.0 (ptetraurelia_mac_51_annotation_v2.0.gff3) and IES annotation v1 (internal_eliminated_sequence_PGM_ParTIES.pt_51.gff3) were used in this study (Arnaiz et al., 2017). The MIC (ptetraurelia_mic2.fa) contigs was used to analyze MIC windows coverage and the TE coverage was done on the v1.0 annotation (ptetraurelia_mic2_TE_annotation_v1.0.gff3) (Guérin et al., 2017). All these files can be downloaded from ParameciumDB (Arnaiz et al., 2020).

### Bioinformatic analyses

Sequencing data were demultiplexed using CASAVA (v1.8.2) and bcl2fastq2 (v2.18.12). Illumina adapters were removed using cutadapt (1.12), keeping only reads with a minimal length of ten nucleotides. Paired-end reads were mapped on expected contaminants (mitochondrial genomes, ribosomal DNA, and bacterial genomes). DNA-seq and ChIP-seq data were mapped on reference genomes using Bowtie2 (v2.3 -X500 –local), RNA-seq data using TopHat2 (v2.1.1 –min-intron-length 15 –max-intron-length 100 –coverage-search –read-mismatches 1), sRNA-seq data using BWA (v0.7.15-n 0).

The IES retention score was calculated with ParTIES software (v1.02 MIRET -max_mismatch 1 - min_non_ambigous_distance 4 -score_method Boundaries) (Denby Wilkes et al., 2016) using the mean score of the two IES boundaries. The statistical significance test was done using a control MAC DNA sample (Lhuillier-Akakpo et al., 2014).

Using the MIC genome coverage analysis as in (Vanssay et al., 2020), the read counts were calculated with bedtools (v2.26.0 multicov -q 30) on MIC non overlapping 1 kb windows. Only the windows corresponding to MIC-limited or MAC-destined (22521 and 75106 respectively) regions were considered for this analysis (Guérin et al., 2017). The counts were normalized (RPKM) using the number mapped reads on the MIC genome. After removing the outliers, the distribution of the normalized coverage was displayed using violin plot function (ggplot2 R package).

Only TE copies of a length superior to 500 nucleotides on contigs bigger than 2kb were considered (N=1180; 770 LINEs 261, TIRs, 136 Solo-ORF and 13 SINEs). The coverage of the TE copies was evaluated on bowtie2 and Tophat alignments, for DNA-seq and RNA-seq respectively, using bedtools (v2.26.0 multicov -q 10). The RPKM for each copy was calculated using the number of mapped reads on the MIC genome. The heatmaps of RPKM (log2) were generated with ComplexHeatmap R package (10.1093/bioinformatics/btw313).

sRNA-seq data were analyzed as previously (Vanssay et al., 2020). Sequencing metrics are available in the Supplementary Table 2.

For ChIP-seq data, PCR duplicates (samtools rmdup) and low-quality reads (sickle v1.200 -q 20) were removed from the analysis. Mapped reads on the MIC genome (mapping quality > 10) were randomly sampled to 10M aligned reads corrected by a normalized factor related to the quantity of material of each sample (See Supplementary Table 3). The bedtools multicov (-q 10) software was used to calculate the read coverage of TE copies (only TE copies > 500nt on MIC contigs > 2kb and covered by at least 10 RPKM in all input samples were considered N=362). ChIP enrichments were calculated over the corresponding Input. The heatmaps of enrichments (log2) were generated with ComplexHeatmap R package. The TE copies were grouped by TE classes (251 LINEs, 54 TIRs, 54 Solo-ORFs and 3 SINEs).

## DATA AVAILABILITY

DNA-seq, RNA-seq, sRNA-seq and ChIP-seq datasets generated for this study have been deposited at the ENA with Study Accession number PRJEB46608. The proteomic data are provided in Supplementary Data. The statistical data and R scripts used to generate the bioinformatic images have been deposited at https://doi.org/10.5281/zenodo.5171885. All other relevant data supporting the key findings of this study are available within the article and its Supplementary Information files or from the corresponding author upon reasonable request.

