## Supplementary Data for "*Paramecium* Polycomb Repressive Complex 2 physically interacts with the small RNA binding PIWI protein to repress transposable elements"

```
sp|O75530|EED_HUMAN_EED                 MS-----EREVS------TAPAGT-----DMPAAKKQKLSSDENSNPDLSGDENDDAVSI      44
TTHERM_00442420_ESC1                    MSFIEKIEYEIEKYFHSSGVKEGFETLQNDFKQYTLEKNTQFQDIRKHLE-TLQQESLQT      59
PCAU.43c3d.1.P00820136                  ---------------------------------MEVEKQDFFQELKE-----LYTNRVKL      22
PTRED.209.2.P71800001293590075          ---------------------------------MDEEKLGFFEQLRQ-----LYQNREKI      22
PSEX.AZ8_4.1.P0350206                   ---------------------------------MDEEKGGFFEQLRQ-----LYQNREKI      22
PBIA.V1_4.1.P00490027                   ---------------------------------MDEEKQSFFEQLRQ-----LYQNREKI      22
PNOV.TE.1.P01200029                     ---------------------------------MDEEKPGFFEQLRQ-----LYQNREKI      22
PQUADEC.NiA.1.P00780073                 ---------------------------------MDEEKLGFFEQLRQ-----LYQNREKI      22
PPRIMP11287                             ---------------------------------MDEEKPGFFEQLRQ-----LYQNREKI      22
POCT.K8.1.P71800002767170233            ---------------------------------MDEEKPGFFEQLRQ-----LYQNREKI      22
PDODEC.274.1.P02380030                  ---------------------------------MDEEKPGFFEQLRQ-----LYQNREKI      22
PTET.51.1.P0240079                      ---------------------------------MDEEKPGFFEQLRQ-----LYQNREKI      22
PDEC.223.1.P00900098                    ---------------------------------MDEEKPGFFEQLRQ-----LYQNREKI      22
                                                                            :*    :: .         :  .

sp|O75530|EED_HUMAN_EED                 ESGTNTERPDTPTNTPNAPGRKSWGKGKWKSKKCKYSFKCVNSLKEDHNQPL--------      96
TTHERM_00442420_ESC1                    EIKIV--------------------------TENQIKDACFLYID---GISLIGVVSKNQ      90
PCAU.43c3d.1.P00820136                  ESKIV------P---PHVS--RQLKITNL--LQNEEAINCMDLIDLNPDESLIMVCSDSK      69
PTRED.209.2.P71800001293590075          ESKIV------G---FHMQ--KQFKISHL--YGNEEQINCMDLIDISENNLLLQYVQML-      68
PSEX.AZ8_4.1.P0350206                   ESKIV------G---FHMQ--KQFKISHL--YGNEEQINCMDLIDISENESFIIVCSDAK      69
PBIA.V1_4.1.P00490027                   ESKIV------G---FHMQ--KQFKISHL--YGNEEQINCMDLIDISENESFIIVCSDAK      69
PNOV.TE.1.P01200029                     ESKIV------G---FHMQ--KQFKISHL--YGNEEQINCMDLIDISDKESFIIVCSDAK      69
PQUADEC.NiA.1.P00780073                 ESKIV------G---FHMQ--KQFKISHL--YGNEEQINCMDLIDISENESFIIVCSDAK      69
PPRIMP11287                             ESKIV------G---FHMQ--KQFKISHL--YGNEEQINCMDLIDISENESFIIVCSDAK      69
POCT.K8.1.P71800002767170233            ESKIV------G---FHMQ--KQFKISHL--FGNEEQINCMDLIDISENESFIIVCSDAK      69
PDODEC.274.1.P02380030                  ESKIV------G---FHMQ--KQFKISHL--YGNEEQINCMDLIDISENESFIIVCSDAK      69
PTET.51.1.P0240079                      ESKIV------G---FHMQ--KQFKISHL--YGNEEQINCMDLIDISETESFIIVCSDAK      69
PDEC.223.1.P00900098                    ESKIV------G---FHMQ--KQFKISHL--YGNEEQINCMDLIDISENESFIIVCSDAK      69
                                        *                                 :    *.  :.      :

sp|O75530|EED_HUMAN_EED                 -------------------FGVQFNWHSKEGDPLVFATVGSNRVTLYECHSQGEIRLL--      135
TTHERM_00442420_ESC1                    IYVFYSSDSASSNINLMSQHTIQFNISD---NPYQYQSYP--------KSKQKIVNFVKG      139
PCAU.43c3d.1.P00820136                  IKL-YSH---KSELNKLFEFVIQYEVNE---H-LDYEVVK--------QGKQ--KRFLQL      111
PTRED.209.2.P71800001293590075          ------------KQNYIRILLNQINSNE---H-LDYDGLK--------NEKQ--KKFLRH      102
PSEX.AZ8_4.1.P0350206                   IKL-YSH---TTQLDKLFEFVIQFEVNE---H-LDYDGLK--------SEKQ--KKFLRH      111
PBIA.V1_4.1.P00490027                   IKL-YSH---TTKLDKLFEFVIQFEANE---H-LDYDGLK--------NEKQ--KKFLRH      111
PNOV.TE.1.P01200029                     IKL-YSH---TTKLDKLFEFVIQFEANE---H-LDYDGLK--------NEKQ--KKFLRH      111
PQUADEC.NiA.1.P00780073                 IKL-YSH---TTKLDKLFEFVIQFEANE---H-LDYDGLK--------NEKQ--KKFLRH      111
PPRIMP11287                             IKL-YSH---TTKLDKLFEFVIQFEANE---H-LDYDGLK--------NEKQ--KKFLRH      111
POCT.K8.1.P71800002767170233            IKL-YSH---TTKLDKLFEFVIQFEANE---H-LDYDGLK--------NEKQ--KKFLRH      111
PDODEC.274.1.P02380030                  IKL-YSH---TTKLDKLFEFVIQFEANE---H-LDYDGLK--------NEKQ--KKFLRH      111
PTET.51.1.P0240079                      IKL-YSH---TTKLDKLFEFVIQFEANE---H-LDYDGLK--------NEKQ--KKFLRH      111
PDEC.223.1.P00900098                    IKL-YSH---TTKLDKLFEFVIQFEANE---H-LDYDGLK--------NEKQ--KKFLRH      111
                                                              * :  .   .   :              .*   .::

sp|O75530|EED_HUMAN_EED                 -QSYVDADADENFYTCAWTYDSNTSHPLL--AVAGSRGIIRIINPITMQCIKHYVGHGNA      192
TTHERM_00442420_ESC1                    YRESVFPKTEEDLYSCDFFDLHLEKNIYTGVIAGGMTGFIHFVDIQSSTNQICFQVHGDT      199
PCAU.43c3d.1.P00820136                  SKELTLDRVKEVLLFCHFAYVNNQ----LFVLAGGEVGLVYQIEIKENPEYSILEGHHQI      167
PTRED.209.2.P71800001293590075          SKQLTTGPVKELIMFCHFAYIYDQ----LYVFAAGDLGYVYQIEIKENPEYFLLEGHHNI      158
PSEX.AZ8_4.1.P0350206                   SKQLTTGPVKELIMYCHFAYINEQ----LYVFAAGDLGYVYQIEIKENPEYFLLEGHHNV      167
PBIA.V1_4.1.P00490027                   SKQLTTGPVKELIMYCHFAYIHEQ----LYVFAAGDLGYVYQIEIKENPEYFLLEGHHNV      167
PNOV.TE.1.P01200029                     SKQLTTGPVKELIMFCHFAYIHDQ----LYVFAAGDLGYVYQIEIKENPEYFLLEGHHNV      167
PQUADEC.NiA.1.P00780073                 SKQLTTGPVKELIMFCHFAYIHDQ----LYVFAAGDLGYVYQIEIKENPEYFLLEGHHNV      167
PPRIMP11287                             SKQLTTGPVKELIMFCHFGYIHDQ----LYVFAGGDLGYVYQIEIKENPEYFLLEGHHNV      167
POCT.K8.1.P71800002767170233            SKQLTTGPVKELIMFCHFAYIHDQ----LYVFAGGDLGYVYQVEIKENPEYFLLEGHHNV      167
PDODEC.274.1.P02380030                  SKQLTTGPVKELIMFCHFAYIHDQ----LYVFAGGDLGYVYQIEIKENPEYFLLEGHHNI      167
PTET.51.1.P0240079                      SKQLTTGSVKELIMFCHFAYIHDQ----LYVFAGGDLGYVYQIEIKENPEYFLLEGHHNV      167
PDEC.223.1.P00900098                    SKQLTTGPVKELIMFCHFAYIHDQ----LYVFAGGDLGYVYQIEIKENPEYFLLEGHHNI      167
                                         :. .   ..* :  * :              ..*  * :  ::            * :

sp|O75530|EED_HUMAN_EED                 INELKFHPR------DPNLLLSVSKDHALRLWNIQTDTLVAIFGGV--EGHRDEVLSADY      244
TTHERM_00442420_ESC1                    IFDIKCLNRNQSCQDYKNIVVSASKDGSIFLSHLAKQEKLIQLKDFLSTSPMSQFTCIDI      259
PCAU.43c3d.1.P00820136                  ITGLS---S------NEKCVFSCSKDGSIIQWDTRNKQIVMIFRD--KDVPGAEVLSI--      214
PTRED.209.2.P71800001293590075          ITGLA---S------NNKGAFSSSKDGSIIQWDVTNKQIVQVLRD--GDTPEAEVLCV--      205
PSEX.AZ8_4.1.P0350206                   ITGLA---S------NSKGIFSSSKDGSIIQWDVTNRQIVQVFRD--GDTPEAEVLCV--      214
PBIA.V1_4.1.P00490027                   ITGLA---S------NNKGAFSSSKDGSIIQWDVTNRQIVQVFRD--GDTPEAEVLCV--      214
PNOV.TE.1.P01200029                     ITGLA---S------NNKAAFSSSKDGSIIQWDVTNRQIVQVFRD--GDTPEAEVLCV--      214
PQUADEC.NiA.1.P00780073                 ITGLA---S------NNKGAFSSSKDGSIIQWDVTNRQIVQVLRD--GDTPEAEVLCV--      214
PPRIMP11287                             ITGLA---S------NNKGVFSSSKDGSIIQWDVTNRQIVQVLRD--GDTPEAEVLCV--      214
POCT.K8.1.P71800002767170233            ITGLA---S------NNKGAFSSSKDGSIIQWDVTNRQIVQVLRD--GDTPEAEVLCV--      214
PDODEC.274.1.P02380030                  ITGLA---S------NNKGAFSASKDGSIIQWDVTNRQIVQVLRD--GETPEAEVLCV--      214
PTET.51.1.P0240079                      ITGLA---S------NNKGAFSSSKDGSIIQWDVTNRQIVQVLRD--GDTPEAEVLCV--      214
PDEC.223.1.P00900098                    ITGLT---S------NNKGAFSSSKDGSIIQWDVTNRQIVQVLRD--GDTPEAEVLCV--      214
                                        *  :             :  .* *** ::   .  .   :  : .        :. .

sp|O75530|EED_HUMAN_EED                 DLLGEKIMSCGMDHSLKLWRINSKRMMNAIKESYDYNPNKTNRPFISQKI---HFPDFST      301
TTHERM_00442420_ESC1                    DLACENIFSACGDGTIQVWQLSEQIKKIL-----LLKTPV--NNFVKIEKSQYNYQVQAQ      312
PCAU.43c3d.1.P00820136                  TTNEGYVLGSYTDGKLRFWKIEAEKIDNW-----THQTQL--KPYRV-------FQSPLQ      260
PTRED.209.2.P71800001293590075          SANQEHVIGAYSDGKLKLWTIQDYKNDSW-----TYKINK--QPTII-------FQSPLQ      251
PSEX.AZ8_4.1.P0350206                   SVNQEYLIGAYSDGKLKLWTIQDYKNDSW-----TYKINK--QPTII-------FQSPLQ      260
PBIA.V1_4.1.P00490027                   SANQEHVIGAYSDGKLKLWTIQDYKNESW-----TYKINK--QPTII-------FQGPLQ      260
PNOV.TE.1.P01200029                     SANQEHVIGAYSDGKLKLWTIQDYKNDSW-----TYKINK--QPTII-------FQGPLQ      260
PQUADEC.NiA.1.P00780073                 SANQEHVIGAYSDGKLKLWTIEHYKNDSW-----TYKINK--QPTII-------FQGPLQ      260
PPRIMP11287                             SANQEHIIGAYSDGKLKLWTIQDYKNDSW-----TYKINK--QPTII-------FQGPLQ      260
POCT.K8.1.P71800002767170233            SANQEHVIGAYSDGKVKLWTIQDYKNDSW-----TYKINK--SPTII-------FQGPLQ      260
PDODEC.274.1.P02380030                  SANQEHVIGAYSDGKLKLWTIQDYKNDSW-----TYKINK--SPTII-------FQGPLQ      260
PTET.51.1.P0240079                      SANQEHVIGAYSDGKLKLWTIQDYKNDSW-----TYKINK--SPTII-------FQGPLQ      260
PDEC.223.1.P00900098                    SANQEHVIGAYSDGKLKLWTIQDYKNDSW-----TYKINK--SPTII-------FQGPLQ      260
                                              ::..  * .::.* :.              :                 :

sp|O75530|EED_HUMAN_EED                 -RDIHRNYVDCVRWLGDLILSKSCENAIVCWKPGKMEDDIDKIKPSESNVTILGRFDYSQ      360
TTHERM_00442420_ESC1                    EDIMHTVQVTQCSFYGKYLLYLNSQGYITL--GTFIEPY---VYKAQYRMRK---FDKDN      364
PCAU.43c3d.1.P00820136                  DVYPHIGQV---VLLKHTIV------AKHI--NGTINYF---VFDMQGNVQSLHEWEMSS      306
```

```
PTRED.209.2.P71800001293590075      DIYPHITQI---FLLKQTIV------TKHL--NGILNYF---VFDLQGRTISLRDWEMSN      297
PSEX.AZ8_4.1.P0350206               DIYPHVTQI---FLLKQTIV------TKHL--NGIINYF---VFDLQGRTISLRDWEMSN      306
PBIA.V1_4.1.P00490027               DIYPHVTQI---FLLKQTIV------TKHL--NGILNYF---VFDLQGRTISLRDWEMSN      306
PNOV.TE.1.P01200029                 DIYPHITQI---FILKQTIV------TKHL--NGILNYF---VFDLQGRTISLRDWEMSN      306
PQUADEC.NiA.1.P00780073             DIYPHITQI---FLLKQTIV------TKHL--NGILNYF---VFDLQGRTISLRDWEMSN      306
PPRIMP11287                         DIYPHITQI---FLLKQTIV------TKHL--NGILNYF---VFDLQGRTISLRDWEMSN      306
POCT.K8.1.P71800002767170233        DIYPHITQI---FLLKQTIV------TKHL--NGILNYF---VFDLQGRTISLRDWEMSN      306
PDODEC.274.1.P02380030              DIYPHITQI---FLLKQTIV------TKHL--NGILNYF---VFDLQGRTISLRDWEMSN      306
PTET.51.1.P0240079                  DIYPHITQI---FLLKQTIV------TKHL--NGILNYF---VFDLQGRTISLRDWEMSN      306
PDEC.223.1.P00900098                DIYPHITQI---FLLKQTIV------TKHL--NGILNYF---VFDLQGRTISLRDWEMSN      306
                                        *   :       . ::               ::     :   : .      :: ..

sp|O75530|EED_HUMAN_EED             CDIWYM------RFSMDFWQKM--LALGNQVGKLYVWDLEVEDPHKAKCTTLTHHK----      408
TTHERM_00442420_ESC1                T------GKNLKFFQITNFD------LGSKV-ENY---------NKVMHVDQKNGLVWIG      402
PCAU.43c3d.1.P00820136              DNFWESIGVCFPYVIFTHLNRMQVVSLKSNK-LIYEAELDAFPPSKLTITKVSSRMLLIG      365
PTRED.209.2.P71800001293590075      ENYWESIDVKFPYLIYTCFNRMQVITLKKNK-LTYEAELDVLPSSKLTISKISNRLIVIG      356
PSEX.AZ8_4.1.P0350206               ENYWESIDVKFPYLIYSSFNRMQVITLKKNK-LTYEAELDVLPSSKLTISKISNRLIIIG      365
PBIA.V1_4.1.P00490027               ENYWESIDVKFPYLIYTSFNRMQVITLKKNK-LTYEAELDVLPSSKLTISKISNRLIVIG      365
PNOV.TE.1.P01200029                 ENYWESIDVKFPYLIYTCFNRMQVITLKKNK-LTYEAELDVLPSSKLTISKISNRLIVIG      365
PQUADEC.NiA.1.P00780073             ENYWESIDVKFPYLIYTCFNRMQVITLKKNK-LTYEAELDVLPSSKLTISKISNRLIVIG      365
PPRIMP11287                         ENYWESIDVKFPYLIYTCFNRMQVITLKKNK-LTYEAELDVLPSSKLTISKISNRLIVIG      365
POCT.K8.1.P71800002767170233        ENYWESIDVKFPYLIYTCFNRMQVITLKKNK-LTYEAELDVLPSSKLTISKISNRLIVIG      365
PDODEC.274.1.P02380030              ENYWESIDVKFPYLIYTCFNRMQVITLKKNK-LTYEAELDVLPSSKLTISKISNRLILIG      365
PTET.51.1.P0240079                  ENYWESIDVKFPYLIYTCFNRMQVITLKKNK-LTYEAELDVLPSSKLTISKISNRLIVIG      365
PDEC.223.1.P00900098                ENYWESIDVKFPYLIYTCFNRMQVITLKKNK-LTYEAELDVLPSSKLTISKISNRLIVIG      365
                                                 .     :      * .:    *          *      .

sp|O75530|EED_HUMAN_EED             --------------------------------CGAAIRQTSFSRDSSILIAVCDDASIWR      436
TTHERM_00442420_ESC1                KQQSIFCFDLKDRRNSEIDYNYSKPLIAYNTFIKANIQKIFFNENYSVLYILTDAKQLIK      462
PCAU.43c3d.1.P00820136              RMKSFAIF----------------------------------------------------      373
PTRED.209.2.P71800001293590075      RMNQYTILQQ--------------------------------------------------      366
PSEX.AZ8_4.1.P0350206               RMNQYTILQQ--------------------------------------------------      375
PBIA.V1_4.1.P00490027               RMNQYTILQQ--------------------------------------------------      375
PNOV.TE.1.P01200029                 RMNQYTILQQ--------------------------------------------------      375
PQUADEC.NiA.1.P00780073             RMNQYTILQQ--------------------------------------------------      375
PPRIMP11287                         RMNQYTILQQ--------------------------------------------------      375
POCT.K8.1.P71800002767170233        RMNQYTILQQ--------------------------------------------------      375
PDODEC.274.1.P02380030              RMNQYTILQQ--------------------------------------------------      375
PTET.51.1.P0240079                  RMNQYTILQQ--------------------------------------------------      375
PDEC.223.1.P00900098                RMNQYTILQQ--------------------------------------------------      375

sp|O75530|EED_HUMAN_EED             WDRLR--    441
TTHERM_00442420_ESC1                FSFIPNK    469
PCAU.43c3d.1.P00820136              -------    373
PTRED.209.2.P71800001293590075      -------    366
PSEX.AZ8_4.1.P0350206               -------    375
PBIA.V1_4.1.P00490027               -------    375
PNOV.TE.1.P01200029                 -------    375
PQUADEC.NiA.1.P00780073             -------    375
PPRIMP11287                         -------    375
POCT.K8.1.P71800002767170233        -------    375
PDODEC.274.1.P02380030              -------    375
PTET.51.1.P0240079                  -------    375
PDEC.223.1.P00900098                -------    375
```

**Supplemental Figure 1: Multiple alignment of EED proteins.**

CLUSTAL O (1.2.4) alignment of EED homologues from *P. tetraurelia (*PTET) and other sequenced *Paramecium species* (PCAU, PTRED, PSEX, PBIA, PNOV, PQUADEC, PPRIMP, POCT, PDODEC, PDEC), *Tetrahymena thermophila* (TTHERM) and *Homo sapiens* (HUMAN). The WD repeats in the human protein are highlighted in yellow. The residues forming the aromatic cage encircling the methyl-lysine in mouse EED (Phe-97, Tyr-148 and Tyr-365) are shown in red. Conserved amino acids I193 and L196 involved in Ezh2 interaction are underlined.

```
SUZ12_MOUSE                         MAPQKHGGGGGGGSGPSAGSGGGGFGGSAAAVAAAASGGKSGGGGCGGGGSYSASSSSAA     60
SUZ12_TTHERM_00149839               ------------------------------------------------------------     0
PJENN.M.1.P00090100                 ------------------------------------------------------------     0
PCAU.43c3d.1.P00310146              ------------------------------------------------------------     0
PSEX.AZ8_4.1.P0250090               ------------------------------------------------------------     0
PQUADEC.NiA.1.P00110192             ------------------------------------------------------------     0
PDEC.223.1.P00020098                ------------------------------------------------------------     0
PDODEC.274.1.P00030079              ------------------------------------------------------------     0
PTET.51.1.P0190277                  ------------------------------------------------------------     0
POCT.K8.1.P71800002770480261        ------------------------------------------------------------     0
PTRED.209.2.P71800001293730071      ------------------------------------------------------------     0
PBIA.V1_4.1.P00080251               ------------------------------------------------------------     0
PNOV.TE.1.P00020257                 ------------------------------------------------------------     0

SUZ12_MOUSE                         AAAAAAGAAVLPVKKPKMEHVQADHELFLQAFEKPTQIYRFLRTRNLIAPIFLHRTLTYM     120
SUZ12_TTHERM_00149839               ------------------------------------------------------------     0
PJENN.M.1.P00090100                 ------------------------------------------------------------     0
PCAU.43c3d.1.P00310146              ------------------------------------------------------------     0
PSEX.AZ8_4.1.P0250090               ------------------------------------------------------------     0
PQUADEC.NiA.1.P00110192             ------------------------------------------------------------     0
PDEC.223.1.P00020098                ------------------------------------------------------------     0
PDODEC.274.1.P00030079              ------------------------------------------------------------     0
PTET.51.1.P0190277                  ------------------------------------------------------------     0
POCT.K8.1.P71800002770480261        ------------------------------------------------------------     0
PTRED.209.2.P71800001293730071      ------------------------------------------------------------     0
PBIA.V1_4.1.P00080251               ------------------------------------------------------------     0
PNOV.TE.1.P00020257                 ------------------------------------------------------------     0

SUZ12_MOUSE                         SHRNSRTSIKRKTFKVDDMLSKVEKMKGEQESHSLSAHLQLTFTGFFHKNDKPSQNSENE     180
SUZ12_TTHERM_00149839               ------------------------------------------------------------     0
PJENN.M.1.P00090100                 ------------------------------------------------------------     0
PCAU.43c3d.1.P00310146              ------------------------------------------------------------     0
PSEX.AZ8_4.1.P0250090               ------------------------------------------------------------     0
PQUADEC.NiA.1.P00110192             ------------------------------------------------------------     0
PDEC.223.1.P00020098                ------------------------------------------------------------     0
PDODEC.274.1.P00030079              ------------------------------------------------------------     0
PTET.51.1.P0190277                  ------------------------------------------------------------     0
POCT.K8.1.P71800002770480261        ------------------------------------------------------------     0
PTRED.209.2.P71800001293730071      ------------------------------------------------------------     0
PBIA.V1_4.1.P00080251               ------------------------------------------------------------     0
PNOV.TE.1.P00020257                 ------------------------------------------------------------     0

SUZ12_MOUSE                         QNSVTLEVLLVKVCHKKRKDVSCPIRQVPTGKKQVPLNPDLNQTKPGNFPSLAVSSNEFE     240
SUZ12_TTHERM_00149839               ------------------------------------------------------------     0
PJENN.M.1.P00090100                 ------------------------------------------------------------     0
PCAU.43c3d.1.P00310146              ------------------------------------------------------------     0
PSEX.AZ8_4.1.P0250090               ------------------------------------------------------------     0
PQUADEC.NiA.1.P00110192             ------------------------------------------------------------     0
PDEC.223.1.P00020098                ------------------------------------------------------------     0
PDODEC.274.1.P00030079              ------------------------------------------------------------     0
PTET.51.1.P0190277                  ------------------------------------------------------------     0
POCT.K8.1.P71800002770480261        ------------------------------------------------------------     0
PTRED.209.2.P71800001293730071      ------------------------------------------------------------     0
PBIA.V1_4.1.P00080251               ------------------------------------------------------------     0
PNOV.TE.1.P00020257                 ------------------------------------------------------------     0

SUZ12_MOUSE                         PSNSHMVKSYSLLFRVTRPGRREFNGMINGETNENIDVSEELPARRKRNREDGEKTFVAQ     300
SUZ12_TTHERM_00149839               ------------------------------------------------------------     0
PJENN.M.1.P00090100                 ------------------------------------------------------------     0
PCAU.43c3d.1.P00310146              ------------------------------------------------------------     0
PSEX.AZ8_4.1.P0250090               ------------------------------------------------------------     0
PQUADEC.NiA.1.P00110192             ------------------------------------------------------------     0
PDEC.223.1.P00020098                ------------------------------------------------------------     0
PDODEC.274.1.P00030079              ------------------------------------------------------------     0
PTET.51.1.P0190277                  ------------------------------------------------------------     0
POCT.K8.1.P71800002770480261        ------------------------------------------------------------     0
PTRED.209.2.P71800001293730071      ------------------------------------------------------------     0
PBIA.V1_4.1.P00080251               ------------------------------------------------------------     0
PNOV.TE.1.P00020257                 ------------------------------------------------------------     0

SUZ12_MOUSE                         MTVFDKNRRLQLLDGEYEVAMQEMEECPISKKRATWETILDGKRLPPFETFSQGPTLQFT     360
SUZ12_TTHERM_00149839               ------------------------------------------------------------     0
PJENN.M.1.P00090100                 -----MNP------------------------------KLELKFIKTFLDSEIQIIFLFL     25
PCAU.43c3d.1.P00310146              -----MNR------------------------------KLELKLLSRFQSLEDHLSLLFL     25
PSEX.AZ8_4.1.P0250090               -----MNR------------------------------KLELKILSHFQSLEGKLSLLFL     25
PQUADEC.NiA.1.P00110192             -----MNR------------------------------KLELKLLSHFQSLEGKLSLLFL     25
PDEC.223.1.P00020098                -----MNR------------------------------KLELKLLSHFQSLEGNLSLLFL     25
PDODEC.274.1.P00030079              -----MNR------------------------------KLELKLLSHFQSLEGKLSLLFL     25
PTET.51.1.P0190277                  -----MNR------------------------------KLELKLLSHFQSLEGKLSLLFL     25
POCT.K8.1.P71800002770480261        -----MNR------------------------------KLELKLLSHFQSLEGKLSLLFL     25
PTRED.209.2.P71800001293730071      -----MNR------------------------------KLELKLLSHFQSLEGKLSLLFL     25
PBIA.V1_4.1.P00080251               -----MNR------------------------------KLELKLLSHFQSLEGKLSLLFL     25
PNOV.TE.1.P00020257                 -----MNR------------------------------KLELKLLSHFQSLEGKLSLLFL     25

SUZ12_MOUSE                         LRWTGETNDKSTAPVAKPLATRNSESLHQENKPGSVKPAQTIAVKETLTTELQTRKEKDN     420
SUZ12_TTHERM_00149839               ------------------------------------------------------------     0
PJENN.M.1.P00090100                 TYF-------------------------------------------TLFFNLQH------     36
PCAU.43c3d.1.P00310146              HIS-------------------------------------------DNQAGFQH------     36
PSEX.AZ8_4.1.P0250090               HIS-------------------------------------------DNSAGFQH------     36
```

```
PQUADEC.NiA.1.P00110192             HIS-------------------------------------------DNSAGFQH------   36
PDEC.223.1.P00020098                QIS-------------------------------------------DNSAGFQH------   36
PDODEC.274.1.P00030079              HIS-------------------------------------------DNSAGFQH------   36
PTET.51.1.P0190277                  HIS-------------------------------------------DNSAGFQH------   36
POCT.K8.1.P71800002770480261        HIS-------------------------------------------DNSAGFQH------   36
PTRED.209.2.P71800001293730071      HIS-------------------------------------------DNSAGFQH------   36
PBIA.V1_4.1.P00080251               HIS-------------------------------------------DNSAGFQH------   36
PNOV.TE.1.P00020257                 HIS-------------------------------------------DNSAGFQH------   36

SUZ12_MOUSE                         SNESRQKLRIFYQF---------LYNNNTRQQTEARDDLHCPWCTLN-CRKLYSLLKHLK   470
SUZ12_TTHERM_00149839               ------------------------------------------------------------   0
PJENN.M.1.P00090100                 VKAKKYSLQPFII-YLCKSSLIYFQKMLLNNNLLNN---NVYL----------------F   76
PCAU.43c3d.1.P00310146              IGSQKEQFTVFFNICMNQIGQLFFLKNGTNPQPFSR---NCIFCQNVKIQSSLQLIVHLL   93
PSEX.AZ8_4.1.P0250090               VGSQKEQFTAFYNICMQKINTIVFLKNGTQNQPFEQ---QCIFCTKLKTNSSQQLILHLL   93
PQUADEC.NiA.1.P00110192             VGSQKEQFTAFYNICIQKITKIIFLKNGTQNQPFEQ---QCIFCSKLKTNSSQQLILHLL   93
PDEC.223.1.P00020098                VGSQKEQFTAFYNICIQKITKIVFLKNGTQNQPFEQ---SCIFCSKLKTNSSQQLILHLL   93
PDODEC.274.1.P00030079              VGSQKEQFTAFYNICIQKITKIVFLKNGTQNQPFEQ---SCIFCAKLKTNSSQQLILHLL   93
PTET.51.1.P0190277                  VGSQKEQFTAFYNICIQKITKIVFLKNGTQNQPFEQ---SCIFCAKLKTHSSQQLILHLL   93
POCT.K8.1.P71800002770480261        VGSQKEQFTAFYNICIQKITKIVFLKNGTQNQPFEQ---SCIFCAKLKTNSSQQLILHLL   93
PTRED.209.2.P71800001293730071      VGSQKEQFTAFYNICIQKITKIIFLKNGTQNQPFEQ---QCIFCSKLKTNCSQQLILHLL   93
PBIA.V1_4.1.P00080251               VGSQKEQFTAFFNICIQKITKIIFLKNGTQNQPFEQ---QCIFCSKLKTNSSQQLILHLL   93
PNOV.TE.1.P00020257                 VGSQKEQFTAFYNICIQKITKIIFLKNGTQNQSFEQ---QCIFCSKLKTNSSQQLILHLL   93

SUZ12_MOUSE                         -LCHSRFIFNYVYHPKGA-------------------------RIDVSINECYDGSYAGN   504
SUZ12_TTHERM_00149839               ------------------------------------------------------------   0
PJENN.M.1.P00090100                 NLYGIKVKFKLIIF-----YLISQYKIQTLNLPIIKIIHGLFQIITENLTQCFETLQSQ-   130
PCAU.43c3d.1.P00310146              LFHRIKFSYKFTYYTNDKVLHISYFRQQPKQYP---NSFKYAKVIDERSLECYLFLNLFS   150
PSEX.AZ8_4.1.P0250090               LFHKIKYSYKFTYYQNEKVLHIAYFRQQPKILP---NVFRHYKALNQNQLDCYMFLTLFS   150
PQUADEC.NiA.1.P00110192             LFHKIKYSYKFTYYLNEKVLHIAYFRQQPKILP---NVFRHYKAINQNSLDCFMFLTLFS   150
PDEC.223.1.P00020098                LFHKIKYSYKFTYYQNEKVLHIAYFRQQPKILP---NVFRHYKAINQNQLDCFMFLTLFS   150
PDODEC.274.1.P00030079              LFHKIKYSYKFTYYQNEKVLHIAYFRQQPKILP---NVFRHSKAINQNQLDCFMFLTLFS   150
PTET.51.1.P0190277                  LFHKIKYSYKFTYYQNEKVLHIAYFRQQPKILP---NVFRHYKAINQNQLDCFMFLTLFS   150
POCT.K8.1.P71800002770480261        LFHKIKYSYKFTYYQNEKVLHIAYFRQQPKILP---NVFRHYKAINQNQLDCFMFLTLFS   150
PTRED.209.2.P71800001293730071      LFHKIKYSYKFTYYQNEKVLHIAYFRQQPKLLP---NVFRHYKAINQNQLDCFMFLTLFS   150
PBIA.V1_4.1.P00080251               LFHKIKYSYKFTYYQNEKVLHIAYFRQQPKILP---NVFRHYKAINQNQLDCFMFLTLFS   150
PNOV.TE.1.P00020257                 LFHKIKYSYKFTYYQNEKVLHIAYFRQQPKILP---NVFRHYKAINQNQLDCFMFLTLFS   150

SUZ12_MOUSE                         PQDIHRQPGFAFSRNGPVKRTPITHILVCRPKRTKASMSEFLESEDG--EVEQQRTYSSG   562
SUZ12_TTHERM_00149839               ---------------------------------------MANQKAKVIINPNINNSQAKG   21
PJENN.M.1.P00090100                 -------------------------------------LIRFQSLNKFNYT-------EIQ   146
PCAU.43c3d.1.P00310146              -----------------------------------QSNSEFHQKNKLPLGQDQHIANIMG   175
PSEX.AZ8_4.1.P0250090               -----------------------------------QSNSEFYQKTTLPLEQDQQISNLMS   175
PQUADEC.NiA.1.P00110192             -----------------------------------QSNSEFYQKNTLPLEQDQQISNLMS   175
PDEC.223.1.P00020098                -----------------------------------QSNSEFYQKNTLPLEQDQQISNLMS   175
PDODEC.274.1.P00030079              -----------------------------------QSNSEFYQKNTLPLEQDQQISNLMS   175
PTET.51.1.P0190277                  -----------------------------------QSNSEFYQKNTLPLEQDQQISNLMS   175
POCT.K8.1.P71800002770480261        -----------------------------------QSNSEFYQKNTLPLEQDQQISNLMS   175
PTRED.209.2.P71800001293730071      -----------------------------------QSNSEFYQKNTLPLEQDQQISNLMS   175
PBIA.V1_4.1.P00080251               -----------------------------------QSNSEFYQKNTLPLEQDQQISNLMS   175
PNOV.TE.1.P00020257                 -----------------------------------QSNSEFYQKNTLPLEQDQQISNLMS   175
                                                                              .

SUZ12_MOUSE                         HNRLYFHSDTCLPLRPQEMEVD-SEDEKDPEWLREKTITQIEEFSDVNEGEKEVMKLWNL   621
SUZ12_TTHERM_00149839               Q-RHYFESLDNFGKLIKEQDVSDLDSEYDIKDSVHQQFNQDILYTELDRDDQDFMSLWNQ   80
PJENN.M.1.P00090100                 QVKNNCKSTKNFRIVQRIEEFNEL------KTLYFEAFKQNKLQSITRIQKYFNMQRW--   198
PCAU.43c3d.1.P00310146              QKQFFCGPAKNFKPLTTMEEFKEK------EQIDFESFKQNELQCLLKSTPEFKDTPIQE   229
PSEX.AZ8_4.1.P0250090               QKQFYCGPAKNFRPVQRIEEFKEL------EALDFEAYKQNELLCLTKSSPEFKGSPVQQ   229
PQUADEC.NiA.1.P00110192             QKQFYCGPAKNFKPIQRIEEFKEY------EQLDFEAYKQNELLCLTKSSPEFKGSPIQQ   229
PDEC.223.1.P00020098                QKQFYCGPAKNFKPVQRIEEFKEL------EALDFEAYKQNELLCLTKSSPEFKGSPIQQ   229
PDODEC.274.1.P00030079              QKQFYCGPAKNFKPVQRIEEFKEL------EALDFEAYKQNELLCLTKSSPEFKGSPIQQ   229
PTET.51.1.P0190277                  QKQFYCGPAKNFKPVQRIEEFKEL------EALDFEAYKQNELLCLTKSSPEFKGSPIQQ   229
POCT.K8.1.P71800002770480261        QKQFYCGPAKNFKPVQRIEEFKEL------EALDFEAYKQNELLCLTKSSPEFKGSPIQQ   229
PTRED.209.2.P71800001293730071      QKQFYCGPAKNFKPIQRIEEFKEF------ESLDFEAFKQNELLCLTKSSPEFKGSPIQQ   229
PBIA.V1_4.1.P00080251               QKQFYCGPAKNFKPVQRIEEFKEL------EALDFEAFKQNELLCLTKSSPEFKGSPIQQ   229
PNOV.TE.1.P00020257                 QKQFYCGPAKNFKPVQRIEEFKEL------ESLDFEAFKQNELLCLTKSSPEFKGSPIQQ   229
                                    : :        :       :..        :    :  .*

SUZ12_MOUSE                         HVMKH----------------------GFIADNQMNHACMLFVENYGQKIIKKNLCRNFM   659
SUZ12_TTHERM_00149839               FMLKYNEKSNVKRSTSINSLNIHSLNGEFLKPSKFIFILDEFI-NENFDVLKNTLRNQFL   139
PJENN.M.1.P00090100                 --------SYERKAIIMK----------------------RF--SKVIQQIKRRVENLFL   226
PCAU.43c3d.1.P00310146              FMILYNTKVLEQRPLNMK----------------------EF--LKQFTQLQGNLAICFQ   265
PSEX.AZ8_4.1.P0250090               FMMLYNTRVMEERPQQLK----------------------DF--LKQFTKLQGELAICFQ   265
PQUADEC.NiA.1.P00110192             FMMLYNTRVLEERPQQLK----------------------EF--LKQFTKLQGELAICFQ   265
PDEC.223.1.P00020098                FMMLYNTRVLEERPQQLK----------------------DF--LKQFTKLQGELAICFQ   265
PDODEC.274.1.P00030079              FMMLYNTRVLEERPQQLK----------------------DF--LKQFTKLQGELAICFQ   265
PTET.51.1.P0190277                  FMMLYNTRVLEERPQQLK----------------------DF--LKQFTKLQGELAICFQ   265
POCT.K8.1.P71800002770480261        FMMLYNTRVLEERPQQLK----------------------DF--LKQFTKLQGELAICFQ   265
PTRED.209.2.P71800001293730071      FMMLYNTRVLEERPQQLK----------------------DF--LKQFTKLQGELAICFQ   265
PBIA.V1_4.1.P00080251               FMMLYNTRVLEERPQQLK----------------------DF--LKQFTKLQGELAICFQ   265
PNOV.TE.1.P00020257                 FMMLYNTRVLEERPQQLK----------------------DF--LKQFTKLQGELAICFQ   265
                                                                             *        ::  :   *

SUZ12_MOUSE                         LHLVSMHDFNLISIMSIDKAVTKLREMQQKLEKGESATPSNEEIAEEQNGTANGFSETNS   719
SUZ12_TTHERM_00149839               NHICTLIQYCIVTPNQFLQIALRLK-----------------------------------   164
PJENN.M.1.P00090100                 E-----------------------------------------------------------   227
PCAU.43c3d.1.P00310146              NHLFTMLCYNLIDQQTFVELSLMIQK----------------------------------   291
PSEX.AZ8_4.1.P0250090               NHLFTLLCYSMIDQQTFVELSLMIQK----------------------------------   291
PQUADEC.NiA.1.P00110192             NHLFTLLCYSMIDQQTFVELSLMIQK----------------------------------   291
PDEC.223.1.P00020098                NHLFTLLCYSMIDQQTFVELSLMIQK----------------------------------   291
PDODEC.274.1.P00030079              NHLFTLLCYSMIDQQTFVELSLMIQK----------------------------------   291
PTET.51.1.P0190277                  NHLFTLLCYSMIDQQTFVELSLMIQK----------------------------------   291
POCT.K8.1.P71800002770480261        NHLFTLLCYSMIDQQTFVELSLMIQK----------------------------------   291
```

```
PTRED.209.2.P71800001293730071       NHLFTLLCYSMIDQQTFVELSLMIQK---------------------------------          291
PBIA.V1_4.1.P00080251                NHLFTLLCYSMIDQQTFVELSLMIQK---------------------------------          291
PNOV.TE.1.P00020257                  NHLFTLLCYSMIDQQTFVELSLMIQK---------------------------------          291
```

**Supplemental Figure 2: Multiple alignment of Suz12.like proteins.**
CLUSTAL O (1.2.4) alignment of SUZ12 homologues from *P. tetraurelia (*PTET) and other sequenced *Paramecium* species (PCAU, PTRED, PSEX, PBIA, PNOV, PQUADEC, PPRIMP, POCT, PDODEC, PDEC), *Tetrahymena thermophila* (TTHERM) and *Mus musculus* (MOUSE). The *Paramecium* Suz12.like proteins lack the C2 RBBP4-binding domain (blue). A C2H2-type zinc-finger domain is highlighted in yellow. The VEFS-Box (green) is not conserved in *Paramecium* but an acidic patch (residues 189-205 in the *Paramecium tetraurelia* Suz12.like protein), shown in grey, is detected by EMBOSS/charge.

```
                                                        Zinc-finger CHHC
                                                    --------------------
RF2          1 MKQIEIDKESDQPTIQE-ETLLKTEQYLCPANRN--H--KLQSLDKLLSHLHTCKEVKLA----DKPR------------
TT_RNF1      1 MK-----KEFIEIKIKG----------------------KIQAVEHILYN------------------------------
RF1A         1 MQ-----NPENPDSIQQ-----PNFNQECPYNSVQLHTITFNNVWDYLVHLQKCHLLNQR----SEIRQSS---------
RF1B         1 MQ--------NPESIQQ-----PLFNQECPYNSVQLHTIVFNNVWDYLVHLQKCHLLNQR----SEIRQSS---------
RF3          1 MK------------------------FTCPQKE---HLIKFSSVWDYLNHLQKCHIINERITTYGETRKYQCSLNRIVNI
RF4          1 MK------------------------FACPQKA---HAIQFESAWDYLNHLQKCHLINDRIKDCGETRNEY---------
TT_RNF2      1 MEVE--VQEEEAKQIEYVRPLGQICRFECPANKEQ-HSKFLNSYQMYKYHIKYCRELRCL----QEV-------------

                       Zinc-finger CHHC/CCHC
                     ---------------------------
RF2         60 ----FVYYCQYYQ---GHCFLTTEDREKHERTCQWRIGLKKQPDPTANKYSIIQPLGK---PINPQQQQKPVAYTDQQQR
TT_RNF1     24 ------Y-----------------DRFYHEK----------------KKLKVPLPFFYDQ-FASDKVLDNILIYMDNDHT
RF1A        58 --QY--YMCHRC----LKVFEELQQLLDHKD--TEHIMNFEMVFD--KKPEQWINLSK---KGRWEIIKLIENYKD----
RF1B        55 --EY--YMCHNC----LKVFEEFQQLLDHTD--SEETKTFSKVFD--KMPGHSIHLSK---QGKWDIIKLIENYKE----
RF3         54 IFAPFAYKCSNCKRKGIITSQSVKRRIAPQF--LKIPMQFKSL-----SAKSSIPLGR---TGSSKIQELIKQYKS--RP
RF4         45 ------YFCASC----LQVFELITERDMHQH--LCKVDNFKQLF---KDVNKPIPLGK---SGTSKIQELIRFYKDIIHP
TT_RNF2     61 ------FVCKFC----VKLFLEIKKRDEHQINCSCRLVQENVIFEIPDHLKTRYPLNKITFENQEEFYQIIDKFPL-QEP

RF2        130 NYEYAI------------KKGLIKKKAYDEYIPEDGFEFNKD-FKKVDELMKLQIQKNISDQ---NQNNIQFTFKTHFFM
TT_RNF1     64 NFFNSII-----------SVGYLKND--LQYFQNTAFSNNVM------------------NK----------------FL
RF1A       119 -------------------HYLNKKD----HQPESIFPLH-----KTDP--RLNIPRDTVAK--IRNLQIEKKEKLQLYL
RF1B       116 -------------------NYLAGKD----SDTDNVFPSH-----KADP--RLNTPRDTITK--IKNLQIDKKEKLQLYL
RF3        122 NA----------------QKQMIMMK----EDAYNIFDQDVV--ITADP--RLSIPKIEFNQ--IKAANISIIDRISFNL
RF4        107 K-----------------QRNTIDSD----QTEKDDFQQEII--EQSDS--RAFLSKTVENS--FNPKTMIKNSRLTFSL
TT_RNF2    130 LSINLIDSSCLKSDFDEEGEELIKTDKSCCENSNNVFSEDLMCSDEDDS--QLSELEQFSPKKYLQNYIGSNNSQQNLSL

RF2        194 VQPQNKQLIEFLIKRIPYNLYF----------------RKHKFYRE----------------------------------
TT_RNF1     97 VSSENRQFTEFFQIY---NES-----------------NKEELSGF----------------------------------
RF1A       167 LQSKV-LESAYHKLF---NHYKDLSMINKEF-------------------------------------------------
RF1B       164 LQSKV-LESAYYKLF---NHYKDLSMINKEYPYKL---TKDSFHQL----------------------------------
RF3        176 LGGHAKMEAAYFKLH---NNYK-LEQVNRDYSFKI---KQDKLKQI----------------------------------
RF4        160 LGGHTKMEAAYYKLF---NKQV-IEQINKEYPFKI---TPESFKLF----------------------------------
TT_RNF2    208 ISQQIDILSVQEQLIQPSNTIM-VEEEEQDYDDKKPSRKKHSLRSFKNIIKKKIEQPKAIQESAVLQQQIVSQNNQKLTH

RF2        224 ----EEKNQI----VQKFLMKEKKREQRRQK-----------------------------GKPDIKYN-MVNQIPNDILL
TT_RNF1    123 --------NINCAMIEYNYIQQKAAFEKKLK-----------------------------G----------NSQPT----
RF1A       194 ------LSSIN----GQTQYQIEQVCEFMET--------------------------------------AL-TNP-----
RF1B       203 ----MAKHNIN---LSQKLYEQTKVCEFMET--------------------------------------AL-TNP-----
RF3        215 ----QITHNFQ---ITKKLLEDSRVLEFCES--------------------------------------KLDKNP-----
RF4        199 ----FQKNNIT---LSKELLNEAKVCEFCES--------------------------------------KIDQNP-----
TT_RNF2    287 QVEQFFNNNIEL-PFEQQILQQKKILEIQASLVNEKITDLQNLKLIIDQKNEKESSKLECGQEEAQNQKSISSNQ-----

RF2        266 SLLNFNTYEDYLDKQMDEFV----------------------------------------NCKKMEEENVLTSTYLM---
TT_RNF1    152 ----TSSYPNQLNKNIKE------------------------------------------QCDSLREE------------
RF1A       220 -----EDHIMYMNRRQINKFFY--------------------------------------LWILIHEDVNPFPGVQF---
RF1B       232 -----EEHIMYMNRRAINKFFY--------------------------------------LWILIHEDVNPFPGVQF---
RF3        245 -----EIYLQFIQNKGQKQF-----------------------------------------WLLIPESQNLVKDFCF---
RF4        229 -----EMYLQIIQTNNSKSI-----------------------------------------WLMIPEIHNPVKEFGF---
TT_RNF2    361 -----KSQKNYIDKDDDDSFCFLEAQECILNGIHNNQKQYDRFFKVKQHIQIKVMDVTSENQVILQQEENFIDDYNYVSY

RF2        303 -----------------------------------------------TTGKKAIFILQIGRD-----------FKDIFGR
TT_RNF1    174 --------------------------------------------------------LSVICD------------------
RF1A       254 --------------------------------------------------QEPWVILAIPTS------------------
RF1B       266 --------------------------------------------------QEPWVIIAIPTS------------------
RF3        276 --------------------------------------------------DEPWCIFNIVVT------------QSTNQN
RF4        260 --------------------------------------------------DEPWCVFKIMTK------------QVICEK
TT_RNF2    436 LSMKNFQKFNSYLVKNKSKLQPQDHLIASYTVNHYDSKKYLEEHLMKTPNDNQFCLMQIEITKLQDLFNTKQPHNCLFED

RF2        325 NINLLDFKAIFADFLAFSFKRDENM-----------NKQIEPYLTQTS--------EHVESGKNQKQNQNIFEEEIDIQQ
TT_RNF1    180 KINIMNMRMQ-SD---LQQRKQESN--------------------------------QYKQRKKRLQAQLKQEDELNKKL
RF1A       266 SEQLLNLP----------TKKPTDN---------------------------------IQLLELQKLLNEQLEI--AKSI
RF1B       278 SEQLLNLS----------VHKPSDN---------------------------------TDLLELQKQLNEQQEI--AKQI
RF3        294 TQDSIQMS----------TQKQDPI---------------------------------IQQLQMQLQKQEELSVLLKEKL
RF4        278 SIDLIKI------------KKQDPL---------------------------------ISQLEYQIQQEELRGIRLKQQL
TT_RNF2    516 SSNITDSI----------YKSQYSIFILVLENSKIFDKDFVPQPPQNALDCTDVYMKPMQQYSFQQKQLTAQQAKIQNKS
```

```
                                                                             coiled-coil
RF2        386 QR---------------------------------------------LIGELKAKL-EIQKQKNQEDKSQL--------E
TT_RNF1    224 VR---------------------------------------------MIHELTAQK-KVLEQQKQNLL-----------K
RF1A       301 EQ---------------------------------------------EKSKFDQKKIEESKVLEQK-------------I
RF1B       313 EQ---------------------------------------------EKSKFDQKKIEESKVLEQK-------------I
RF3        331 QN---------------------------------------------SFKEIEQAK-DINQTMIQA-------------Q
RF4        313 QN---------------------------------------------SVDDLDSVL-KYLTKKNQK-------------L
TT_RNF2    586 ENKSQLSNDCNETIRLNLYAFEGFQVFNNDLNLHEVYLFDKLSIELLEATAIDLKILEHLKSQTQKQLFQYSNDIEIYSK

                                   coiled-coil
RF2        412 RNQRELQELIEKKKCLIQKESDIEQQIKRVNAQI---NSYHSKIKSYKQE--LELKNQEKIKILYDQKEKELNEF---ID
TT_RNF1    247 NFSEELQKEVQKKL-----------------------KDYQDMLRAKHKDEILKLEQT----------------------
RF1A       323 QKLENLETQLRIQLT----TQNKQL------------NDFESVKQQLKEQSIIDLQDRSTKLDHYYQVQR-TK-------
RF1B       335 LKLENLETQLRIQLT----AQNKQQ------------NDFES--------SILDLQDRSSKLDHYYQVQR-VK-------
RF3        352 DKNETLLVQLQIKLR----EMD---------------LGFESVKEQYNAINIKKLEQAESNLAGFYNNQRQLE-------
RF4        334 DQQENLLVKLQIKLR----END---------------TNNKNLVSQYLALNSKKLQQAEENLIGFYQKMKSME-------
TT_RNF2    666 QKEEKIQQLLEIKSRI---EQDKRATLTKKKARIGKEKNLEELIQ-------KKMSEKENNLLKFNQMQKEIKEMHSQVN

                                coiled-coil
RF2        484 ALNQQVNQYESSQQEIKDEIQSLKEEKQKLKSTINR----------------LQPFKLQLERQLEDLQKQESSVPQQMT-
TT_RNF1    282 -----YNFDLAKRKKITQDISELKNENQSMKQSVES----------------SKKIINDILIKQRDKLTEIQEVKKKIN-
RF1A       379 -----IQSDMTNQSELFKLADSFAKEKLKLLQRVEI----------------YTEKRNKVHYETQDLKAQITNLGEKLR-
RF1B       383 -----IQNDMTNQSEFFKLADSFAKEKQKLLQRVEI----------------YADKRNKIHYETQDLKAQITELGEKLR-
RF3        406 -----TYKSSMRREEQSIDYLQKIRKLTKTKEQ-------------------LHNKKDQIVDQCESLKNQIAVQQNQTN-
RF4        388 -----SEKQITKLDELKVDLNEKELKQKKLKEKVRS----------------LFEEKEQLIQTCNDKKNKIQQTQKQLE-
TT_RNF2    736 QLRQNLQIEIDKEYQLAKEKSDQIYYQQMDKHRAEVISLASENILDLYSVIEFYKQENQLLFKEQYELMDIQRKQGQALL

                                                          RING-HC (C3HC4 type)
RF2        547 -----QSEYYTGNEFVFDISD--------KIKCKACKIRKINVVYIPCGDAALCKIC--HEKFEQLLPHV------CPIC
TT_RNF1    340 DLDQQYQLLCSQSDQQNDIIN-----------CKLCQERPAQIIFRPCGHLIYCESCWIDYLTEERDPYVNQAKYMCLAC
RF1A       437 --QDEQDKLKSKTRRRQRIIQ---------ELCIKCQKNTREVIHFPCQHFLFCIQCIIQGMVEKRQQ--------CPLV
RF1B       441 --QDEQERMKSKTRRRQRIIQ---------DLCIKCQKNTREIIHFPCQHFLFCTQCIIQGMVEKRQQ--------CPLV
RF3        461 ------EKLAKASATKQRIQMIK------QLLCIQCKQAERNVIQFPCQHFLFCETCFLTKISTNHFS--------CPLY
RF4        446 ------DKIAKASVTRKKIQKIK------QLLCFQCQTRERNVIQFPCQHFLFCEQCYLSNIQQKNLK--------CPLK
TT_RNF2    816 KYDAKYENLQNAKRNKEKLISLQKNKQQTAYLCLKCKKYPQNCLQKPCGHVVFCYSCLNEENEEKEIIF-------CKIC

RF2        606 EQF--VKRVYHFMWENEG--------
TT_RNF1    409 REFRNVCRIEGFSELKEQ--------
RF1A       498 AIG--QCRDQRYLDKKFKTVRFSKND
RF1B       502 AIG--QCRDQRNLDKKFKTVRFSKND
RF3        521 EVQ--ECQVEQ-LSSKYKTANI-ITL
RF4        506 EQN--KCQTEN-LNNKYKLVNI-QNV
TT_RNF2    889 -----KIEIIKFSEIKYKIIH-----
```

**Supplemental Figure 3: Multiple alignment of RING finger proteins**.
CLUSTAL O (1.2.4) multiple protein sequence alignment by MAFTT (v7.471) of *Paramecium tetraurelia* Rf1A, Rf1B, Rf2, Rf3 and Rf4, and *Tetrahymena thermophila* RNF1 (TTHERM_00637350A), and RNF2 (TTHERM_00522660) shaded with BoxShade (version 3.21). All proteins contain a coiled coil domain and a RING-HC domain at the C terminus. Rf2, Rf4 and RNF2 harbor two Zinc fingers in tandem at the N terminus. Rf4 displays 54% identity with Rf4b (PTET.51.1.P0990162), which is encoded by a duplicated gene from the most recent whole genome duplication, and appears to be a pseudo-protein.

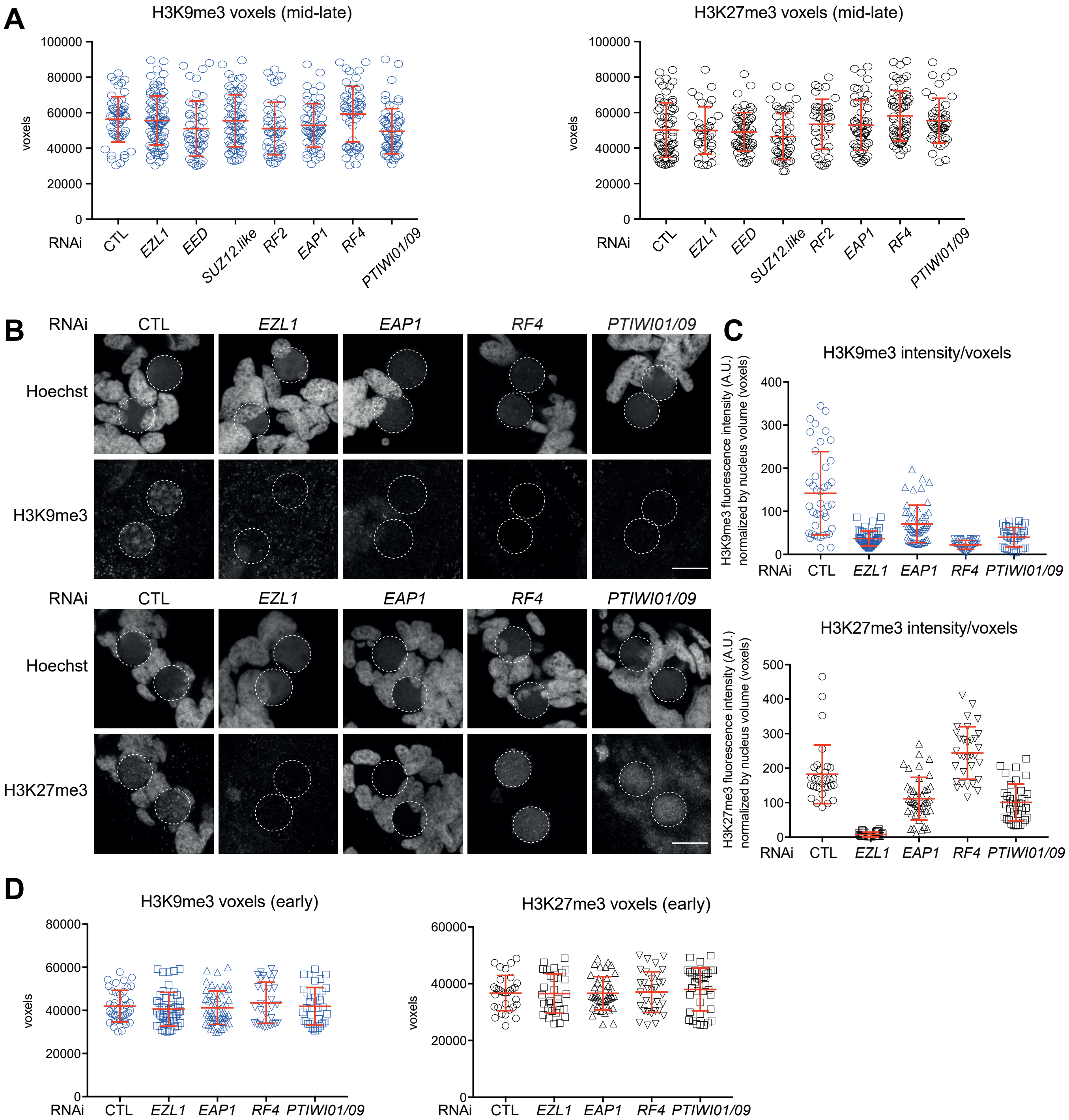

**Supplementary Figure 4. Depletion of Ezl1 cofactors alters H3K9me3 and H3K27me3 accumulation.**

**A.** Boxplot of the estimated nucleus (developing MAC) volume in voxels from the same data as in Figure 2. Estimation of nuclear volume indicated KD cell populations were at comparable developmental stages.

**B.** Immunostaining with H3K9me3 (top) or H3K27me3 (bottom) antibodies at early developmental stages (T=8-13 after the onset of sexual events). Representative images are displayed. Overlay of Z-projections of magnified views of Hoechst staining and H3K27me3- or H3K9me3-specific antibodies are presented. Dashed white circles indicate the new developing MACs. The other Hoechst-stained nuclei are fragments from the old vegetative MAC. In control cells, H3K27me3 is detected in the fragments from the old vegetative MAC, while both H3K9me3 and H3K27me3 are detected in the new developing MACs. Scale bar is 10 µm.

**C.** Quantification of H3K9me3 and H3K27me3 fluorescence signal in the same populations as in B. Bar plots represent total H3K9me3 or H3K27me3 fluorescence intensity in the developing new MAC divided by the number of voxels in the three conditions. Mann-Whitney statistical tests: significant differences with CTL in all conditions (H3K9me3: [P-value<0,0001] for all conditions except *EAP1* [P-value=0,0003]; H3K27me3: [P-value<0,0001] in all conditions except *RF4* [P-value=0,0004])

**D.** Boxplot of the estimated nucleus (developing MAC) volume in voxels from the same data as in panel C. Estimation of nuclear volume indicated KD cell populations were at comparable developmental stages.

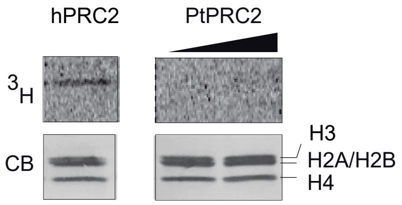

**Supplementary Figure 5. Histone methyltransferase assay.**
Recombinant PRC2 from either human (h) or *Paramecium tetraurelia* (Pt) purified from Sf9 cells by anti-FLAG IP were used for in vitro histone methyltransferase assay with recombinant oligonucleosomes as substrates and S-adenosyl-[methyl 3H]-methionine as methyl donor. The reaction products were separated by SDS-PAGE and transferred onto a PVDF membrane. Coomassie stain (CB, bottom panel) shows histones and the autoradiograph (3H, top panel) indicates H3 histone methyltransferase activity.

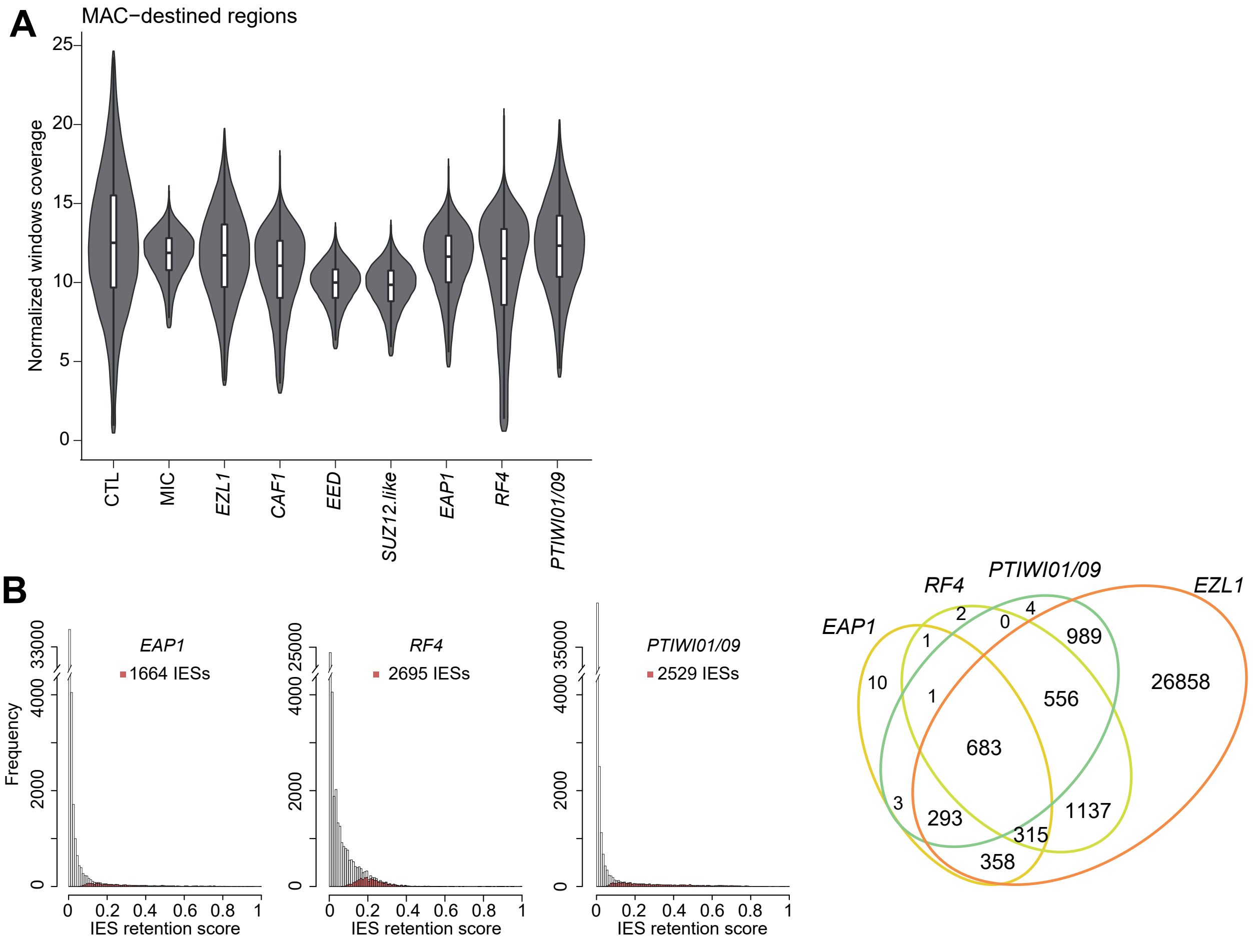

**Supplementary Figure 6. Normalized MAC coverage and IES retention.**

**A.** Violin plot superimposed with a boxplot of normalized coverage of MAC regions using RPKM of 1 kb windows of the MIC assembly for the following samples: control (CTL), MIC and KD of *EZL1, CAF1*, *EED, SUZ12.like, EAP1, RF4* or *PTIWI01/09.* The coverage of MAC-destined regions is similarly covered in all datasets.

**B.** Left: Histograms of IES retention scores (RS) for *EAP1, RF4*, or *PTIWI01/09* KD datasets. The distribution of IES RS for significantly retained IESs is shown in red. RS = ($IES^+$) / ($IES^+$ + $IES^-$), where $IES^+$ are the reads that contain an IES end sequence and $IES^-$are the reads that contain the macronuclear IES junction. Right: Venn diagram of significantly retained IESs upon *EZL1, EAP1*, *RF4 and PTIWI01/09* silencing.

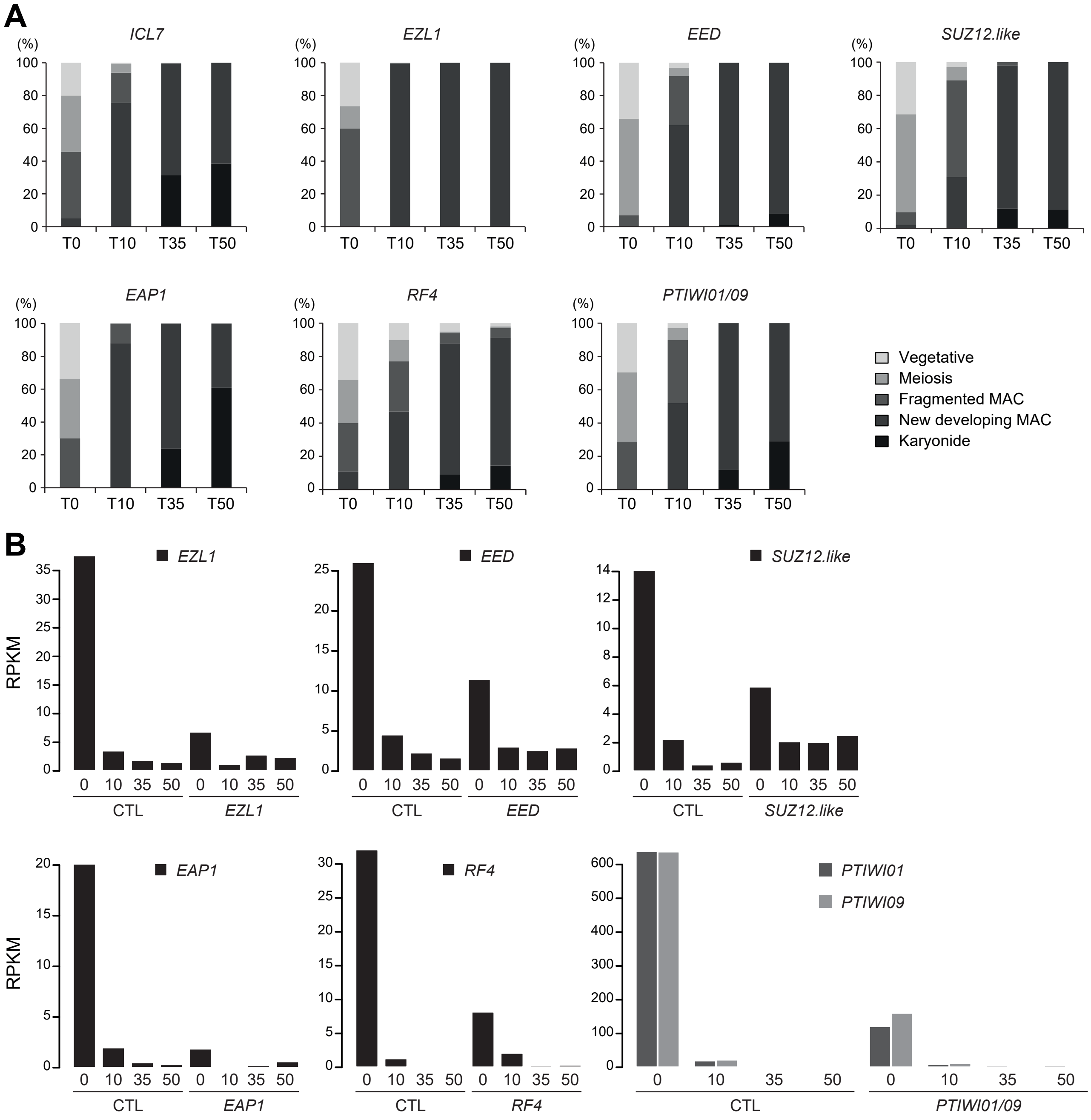

**Supplementary Figure 7. Autogamy progression and knock-down efficiency in RNAi experiments.**

**A.** Progression of autogamy was followed by cytology with Hoechst staining in the time course experiments upon RNAi mediated KD. At least 100 cells were scored for each time point by fluorescence microscopy. The time-points refer to hours after T=0 hours that is defined as the time when cells begin fragmentation of the maternal MAC, as evaluated by cytological observation. Because cells enter autogamy from a fixed point of the cell cycle, a minimum asynchrony of 5–6 h is observed between the first and the last cells to undergo meiosis.

**B.** Steady state mRNA expression levels are measured using normalized read counts (RPKM, Reads per kilobase per million mapped reads) at four time points (T=0; T=10; T=35 and T=50 hours) during autogamy upon CTL KD or *EZL1, EED, SUZ12.like, EAP1, RF4* or *PTIWI01/09* KD. Only the nucleotides outside of the targeted RNAi region are considered.

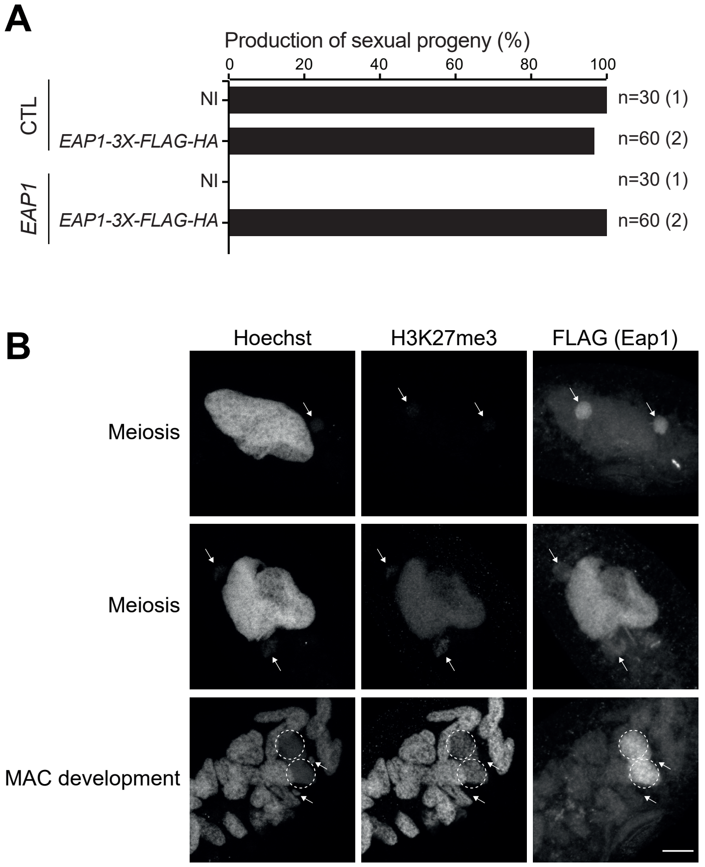

**Supplementary Figure 8. Localization of the functional Eap1-3XFLAG-HA fusion protein.**

**A.** Genetic complementation assays show that the *Eap1-3XFLAG-HA* fusion protein is functional. Production of viable sexual progeny following control or *EAP1* gene silencing in non-injected (NI) cells and *EAP1-3X-FLAG-HA* (RNAi resistant) transformed cells. Cells were starved in each medium to induce autogamy and, following 3-4 days of starvation, post-autogamous cells were transferred individually to standard growth medium to assess the ability of sexual progeny to resume vegetative growth. The total number of cells analyzed for each RNAi and the number of independent experiments (in parenthesis) are indicated. The tagged RNAi-resistant *EAP1* transgene fully complemented the RNAi-mediated depletion of the endogenous Eap1 protein.

**B.** Localization of the Eap1-3XFLAG-HA protein. H3K27me3 and anti-FLAG immunostaining of cells expressing the *EAP1-3X-FLAG-HA* transgene upon *EAP1* RNAi during meiosis and MAC development. Its localization is very similar to that of the Ezl1 and Ptiwi09 proteins, being present in the MIC and in the maternal MAC during meiosis, then accumulating in the new developing MACs. Representative images are displayed. Overlay of Z-projections of magnified views are presented. The arrows show the MICs (one MIC is not visible in the Hoechst staining of the first image) and dashed white circles indicate the new developing MACs. The maternal MAC is fragmented when the new MACs develop. The scale bar is 10 µm.

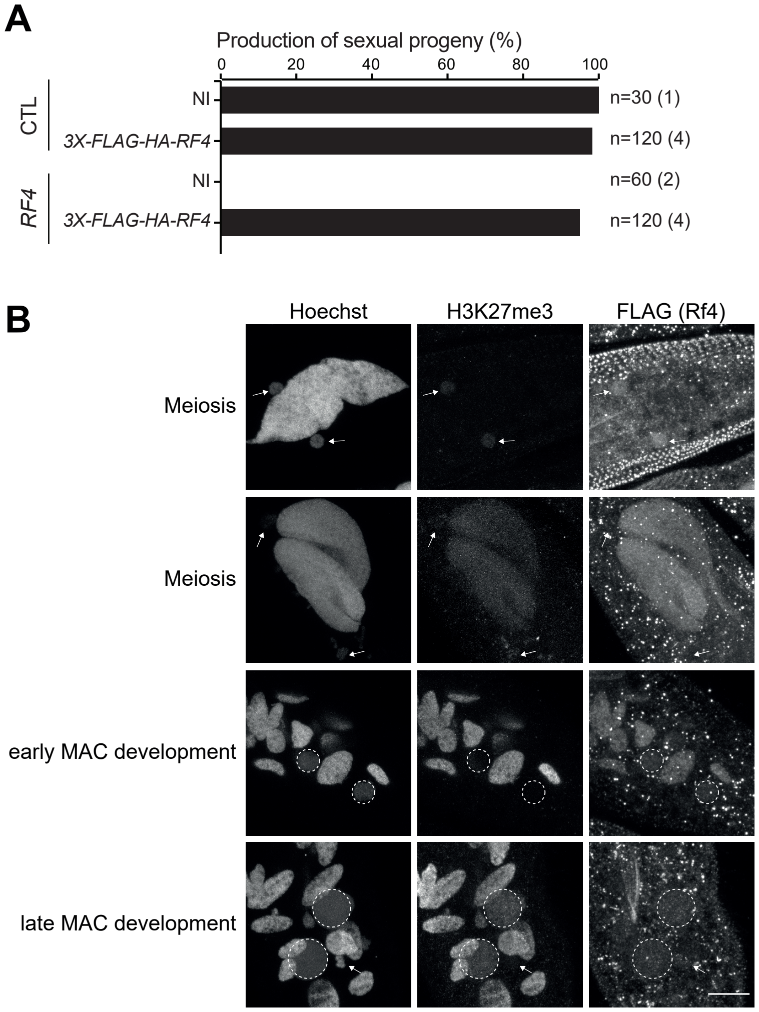

**Supplementary Figure 9. Localization of the functional 3XFLAG-HA-Rf4 fusion protein.**

**A.** Genetic complementation assays show that the 3XFLAG-Rf4 fusion protein is functional. Production of sexual progeny following control or *RF4* gene silencing in non-injected (NI) cells and *3X-FLAG-HA-RF4* (RNAi resistant) transformed cells. Cells were starved in each medium to induce autogamy and, following 3-4 days of starvation, post-autogamous cells were transferred individually to standard growth medium to assess the ability of sexual progeny to resume vegetative growth. The total number of cells analyzed for each RNAi and the number of independent experiments (in parenthesis) are indicated.

**B.** Localization of the 3XFLAG-HA-Rf4 fusion protein. H3K27me3 and anti-FLAG immunostaining of cells expressing the *3X-FLAG-HA-RF4* transgene upon *RF4* RNAi during meiosis and MAC development. Its localization is comparable to that of the Ezl1 and Ptiwi09 proteins (MIC, maternal MAC then new MAC). Representative images are displayed. Overlay of Z-projections of magnified views are presented. The arrows show the MICs, and the dashed white circles indicate the new developing MACs. The other Hoechst-stained nuclei are the maternal MACs (fragmented at the MAC development stage). The scale bar is 10 µm.

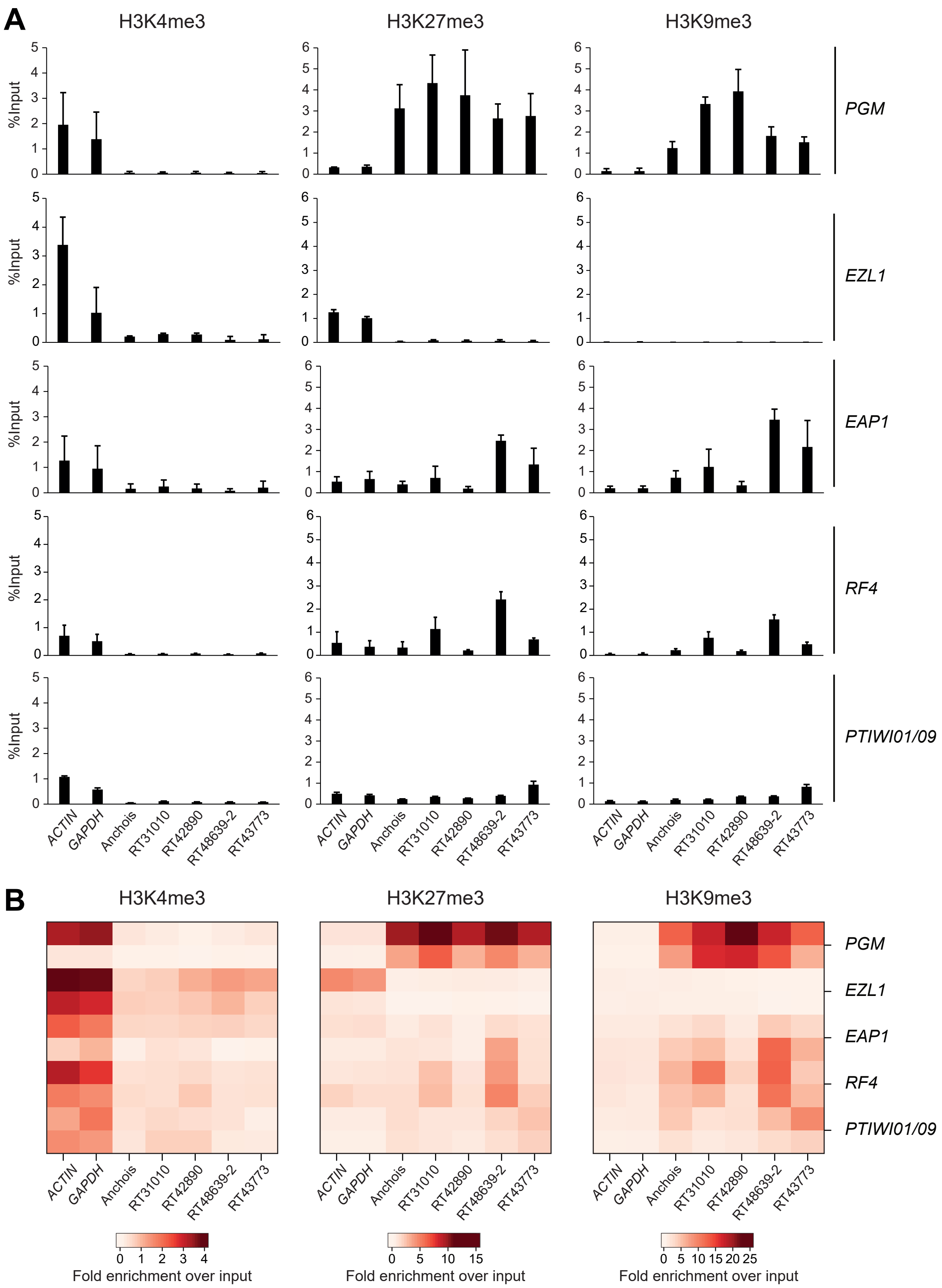

**Supplementary Figure 10. H3K4me3, H3K27me3 and H3K9me3 enrichment on TE copies as determined by ChIP-qPCR.**
**A.** H3K4me3, H3K9me3, and H3K27me3 enrichment at individual copies from different TE families (at T=50 hours) in cells silenced for *PGM, EZL1, EAP1, RF4* or *PTIWI01/09*, as measured by ChIP-qPCR. Quantitative data are expressed as the percentage of ChIP over input. *ACTIN* and *GAPDH* are control genes (Frapporti et al., 2019).
**B.** Heatmap of H3K4me3, H3K27me3 and H3K9me3 fold enrichment over input for the same TE copies and genes as in A. determined by ChIP-seq in two biological replicates.

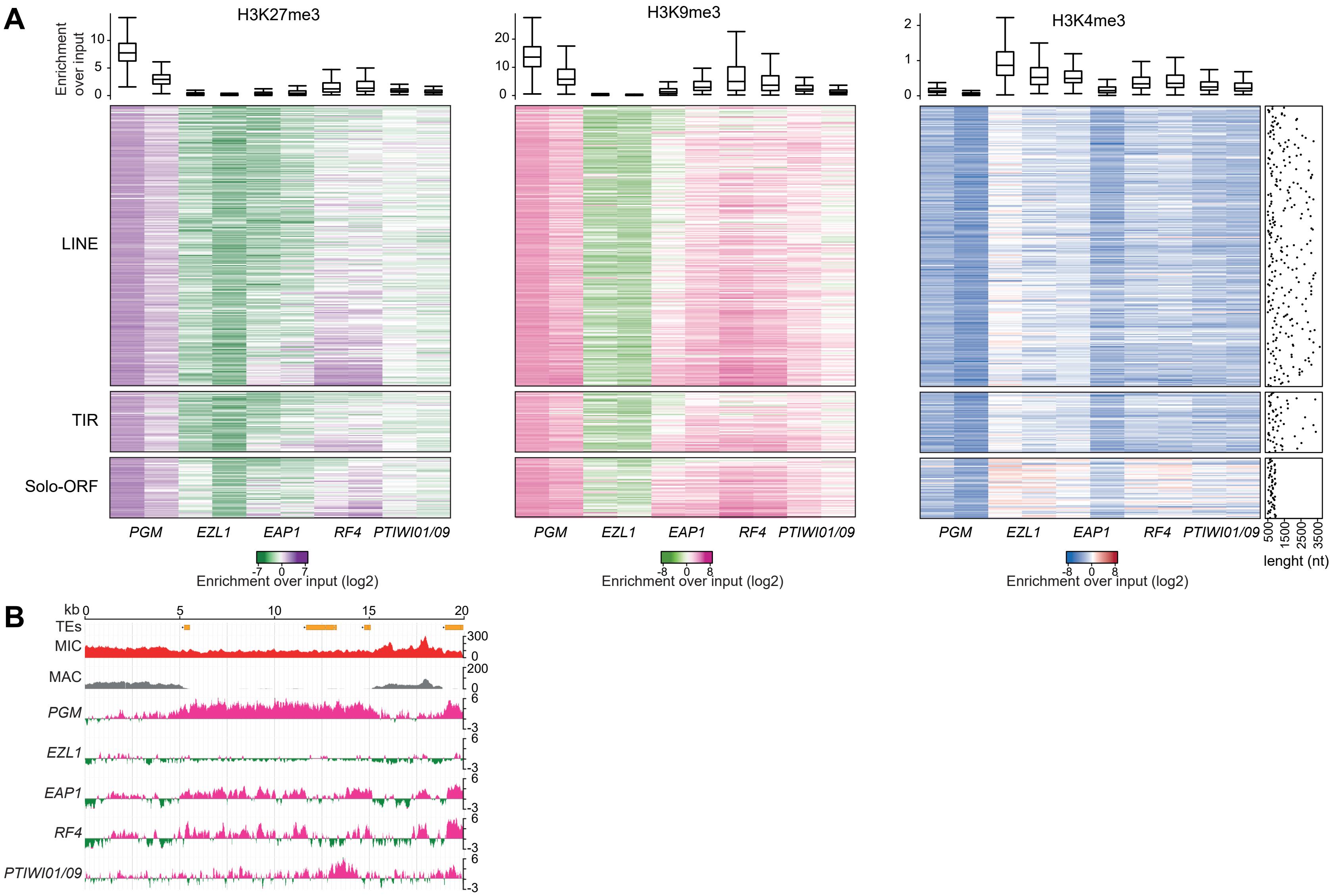

**Supplementary Figure 11. H3K27me3, H3K9me3 and H3K4me3 enrichment for TE copies as determined by ChIP-seq.**

**A.** The heatmap shows H3K27me3, H3K9me3 and H3K4me3 enrichment for TE copies as determined by ChIP-seq for 2 biological replicates in each RNAi condition. The TE copies are grouped by TE families (LINE, TIR, Solo-ORF, SINE) and then ordered by *EAP1* DNA coverage. Colors represent the fold-change (log2) in a given TE family for each histone mark relative to input. The boxplots show the fold enrichment over the input for the considered TE copies (See Materials and Methods). The dotplots show the nucleotide length (nt) for each TE copies.

**B.** Representative genomic region depicting H3K9me3 at TEs (NODE_1273_length_39901_cov_28.471216 between 20kb et 40kb). Lanes show TE annotation, MIC and MAC genome coverages and H3K9me3 normalized enrichment over input (log2) in the different KD conditions.

| Experiment n° | KD gene | Viable progeny | | Regenerants |
|---|---|---|---|---|
| 1 | ND7 | Alive | 30 | |
| | | Dead | 0 | |
| | EZL1 | Alive | 0 | |
| | | Dead | 30 | 2 |
| 2 | ND7 | Alive | 30 | |
| | | Dead | 0 | |
| | EZL1 | Alive | 0 | |
| | | Dead | 30 | |
| 3 | ND7 | Alive | 30 | |
| | | Dead | 0 | |
| | EZL1 | Alive | 0 | |
| | | Dead | 30 | |
| 4 | EZL1 | Alive | 0 | |
| | | Dead | 30 | |
| 5 | SUZ12.like | Alive | 0 | |
| | | Dead | 30 | |
| | EED | Alive | 0 | |
| | | Dead | 30 | |
| | RF1a#1 | Alive | 29 | |
| | | Dead | 1 | |
| | RF1b#1 | Alive | 29 | |
| | | Dead | 1 | |
| | RF2 | Alive | 0 | |
| | | Dead | 30 | |
| | ICL7 | Alive | 30 | |
| | | Dead | 0 | |
| 6 | RF1b#1 | Alive | 29 | |
| | | Dead | 1 | |
| | RF1a+1b#1 | Alive | 28 | |
| | | Dead | 2 | |
| | ND7 | Alive | 28 | |
| | | Dead | 2 | |
| 7 | EAP1 | Alive | 0 | |
| | | Dead | 30 | |
| | ND7 | Alive | 30 | |
| | | Dead | 0 | |
| 8 | RF2 | Alive | 0 | |
| | | Dead | 30 | |
| | ND7 | Alive | 30 | |
| | | Dead | 0 | |
| 9 | RF2 | Alive | 0 | |
| | | Dead | 30 | |
| | ICL7 | Alive | 30 | |
| | | Dead | 0 | |

| | | | | |
|---|---|---|---|---|
| 10 | EAP1 | Alive | 0 | |
| | | Dead | 30 | 17 |
| 11 | RF2 | Alive | 0 | |
| | | Dead | 60 | 1 |
| 12 | EED | Alive | 0 | |
| | | Dead | 30 | |
| | ICL7 | Alive | 29 | |
| | | Dead | 1 | |
| 13 | ICL7 | Alive | 30 | |
| | | Dead | 0 | |
| | EZL1 | Alive | 0 | |
| | | Dead | 30 | 1 |
| 14 | EAP1 | Alive | 0 | |
| | | Dead | 30 | 5 |
| | SUZ12.like | Alive | 0 | |
| | | Dead | 30 | 2 |
| 15 | EAP1 | Alive | 0 | |
| | | Dead | 30 | |
| 16 | EAP1 | Alive | 2 | |
| | | Dead | 28 | 1 |
| 17 | ICL7 | Alive | 60 | |
| | | Dead | 0 | |
| | EAP1 | Alive | 0 | |
| | | Dead | 60 | 22 |
| | PTIWI01/09 | Alive | 1 | |
| | | Dead | 59 | 29 |
| 18 | PTIWI01/09 | Alive | 0 | |
| | | Dead | 30 | 12 |
| | PTIWI01/09 | Alive | 1 | |
| | | Dead | 29 | 18 |
| 19 | ICL7 | Alive | 30 | |
| | | Dead | 0 | |
| | EAP1 | Alive | 0 | |
| | | Dead | 30 | 5 |
| | RF2 | Alive | 0 | |
| | | Dead | 30 | |
| | PTIWI01/09 | Alive | 0 | |
| | | Dead | 30 | 11 |
| 20 | ICL7 | Alive | 30 | |
| | | Dead | 0 | |
| | RF2 | Alive | 0 | |
| | | Dead | 30 | |
| | RF4#2 | Alive | 0 | |
| | | Dead | 30 | |
| | RF1a#1 | Alive | 27 | |
| | | Dead | 3 | |

| | | | | |
|---|---|---|---|---|
| | RF1a#2 | Alive | 30 | |
| | | Dead | 0 | |
| | RF1b#2 | Alive | 30 | |
| | | Dead | 0 | |
| | RF1a#2+RF1b#2 | Alive | 30 | |
| | | Dead | 0 | |
| 21 | ICL7 | Alive | 29 | |
| | | Dead | 1 | |
| | RF2 | Alive | 5 | |
| | | Dead | 25 | |
| | RF4#2 | Alive | 1 | |
| | | Dead | 29 | |
| | RF1a#1 | Alive | 29 | |
| | | Dead | 1 | |
| | RF1a#2 | Alive | 30 | |
| | | Dead | 0 | |
| | RF1b#2 | Alive | 27 | |
| | | Dead | 0 | |
| | RF1a#2+RF1b#2 | Alive | 29 | |
| | | Dead | 1 | |
| 22 | RF4#2 | Alive | 0 | |
| | | Dead | 30 | 7 |
| 23 | SUZ12.like | Alive | 0 | |
| | | Dead | 30 | 2 |
| 24 | RF4#2 | Alive | 1 | |
| | | Dead | 29 | 11 |
| | RF4#2 | Alive | 0 | |
| | | Dead | 30 | 4 |
| 25 | ICL7 | Alive | 29 | |
| | | Dead | 1 | |
| | EZL1 | Alive | 0 | |
| | | Dead | 30 | |
| | EAP1 | Alive | 3 | |
| | | Dead | 27 | 10 |
| | RF4#2 | Alive | 0 | |
| | | Dead | 30 | 12 |
| | PTIWI01/09 | Alive | 0 | |
| | | Dead | 30 | 16 |
| 26 | ICL7 | Alive | 30 | |
| | | Dead | 0 | |
| | EED | Alive | 0 | |
| | | Dead | 30 | |
| | SUZ12.like | Alive | 0 | |
| | | Dead | 30 | |
| | RF2 | Alive | 0 | |
| | | Dead | 30 | 1 |

**Supplementary Table 1: Production of sexual progeny following RNAi-mediated gene silencing.**

In each RNAi experiment (1-26), the number of cells that survived or died is indicated. Cells with a non-functional new MAC (regenerants) were identified as growing survivors able to undergo a novel round of autogamy if starved after ~8 divisions and were not counted as sexual progeny.

| Sequencing | Label | ENA Accession | Number of reads | Aligned reads on the MAC | | Aligned reads on the MIC | |
|---|---|---|---|---|---|---|---|
| DNAseq | EED RNAi | ERS6677533 | 115677916 | 97940715 | 85% | 114229827 | 99% |
| DNAseq | EAP1 RNAi | ERS6677534 | 121174536 | 114582801 | 95% | 119952087 | 99% |
| DNAseq | SUZ12 like RNAi | ERS6677535 | 115874122 | 99177026 | 86% | 114583514 | 99% |
| DNAseq | RF4 RNAi | ERS6677536 | 64341466 | 60847916 | 95% | 63639478 | 99% |
| RNAseq | EAP1 RNAi at T0 | ERS6678311 | 120544116 | 115855252 | 96% | 108492145 | 90% |
| RNAseq | EAP1 RNAi at T10 | ERS6678312 | 98843514 | 94297298 | 95% | 88349814 | 89% |
| RNAseq | EAP1 RNAi at T35 | ERS6678313 | 79026428 | 75035697 | 95% | 70828743 | 90% |
| RNAseq | EAP1 RNAi at T50 | ERS6678314 | 93704426 | 88735917 | 95% | 84144265 | 90% |
| RNAseq | PTIWI01/09 RNAi at T0 | ERS6678315 | 105639316 | 101094185 | 96% | 94913158 | 90% |
| RNAseq | PTIWI01/09 RNAi at T10 | ERS6678316 | 85631710 | 81641834 | 95% | 77099405 | 90% |
| RNAseq | PTIWI01/09 RNAi at T35 | ERS6678317 | 97923744 | 92657101 | 95% | 87811034 | 90% |
| RNAseq | PTIWI01/09 RNAi at T50 | ERS6678318 | 97927868 | 90279890 | 92% | 88072432 | 90% |
| RNAseq | EED RNAi at T0 | ERS6678319 | 128734366 | 122476524 | 95% | 113943899 | 89% |
| RNAseq | EED RNAi at T10 | ERS6678320 | 131947106 | 125197387 | 95% | 117556289 | 89% |
| RNAseq | EED RNAi at T35 | ERS6678321 | 118001468 | 106021195 | 90% | 103522026 | 88% |
| RNAseq | EED RNAi at T50 | ERS6678322 | 124225020 | 110752133 | 89% | 109713934 | 88% |
| RNAseq | RF4 RNAi at T0 | ERS6678323 | 128532682 | 122703613 | 95% | 114187265 | 89% |
| RNAseq | RF4 RNAi at T10 | ERS6678324 | 132427922 | 125880208 | 95% | 118071743 | 89% |
| RNAseq | RF4 RNAi at T35 | ERS6678325 | 122247878 | 115687616 | 95% | 109036516 | 89% |
| RNAseq | RF4 RNAi at T50 | ERS6678326 | 137882198 | 130895035 | 95% | 123318709 | 89% |
| RNAseq | SUZ12 RNAi at T0 | ERS6678327 | 149891700 | 143314807 | 96% | 133434602 | 89% |
| RNAseq | SUZ12 RNAi at T10 | ERS6678328 | 116230092 | 110087863 | 95% | 102817381 | 88% |
| RNAseq | SUZ12 RNAi at T35 | ERS6678329 | 133124084 | 117185608 | 88% | 117932277 | 89% |
| RNAseq | SUZ12 RNAi at T50 | ERS6678330 | 132299518 | 115428311 | 87% | 116822810 | 88% |
| RNAseq | EZL1 RNAi at T0 | ERS6679026 | 72271092 | 68823528 | 95% | 64496008 | 89% |
| RNAseq | EZL1 RNAi at T10 | ERS6679027 | 69918912 | 66269160 | 95% | 62327577 | 89% |
| RNAseq | EZL1 RNAi at T35 | ERS6679028 | 107514064 | 95599260 | 89% | 95843293 | 89% |
| RNAseq | EZL1 RNAi at T50 | ERS6679029 | 95433848 | 83497532 | 87% | 84987545 | 89% |
| RNAseq | ICL7 RNAi at T0 | ERS6679030 | 79765916 | 76013189 | 95% | 71116120 | 89% |
| RNAseq | ICL7 RNAi at T10 | ERS6679031 | 86517736 | 82495013 | 95% | 76931108 | 89% |
| RNAseq | ICL7 RNAi at T35 | ERS6679032 | 109355820 | 104159230 | 95% | 96948071 | 89% |
| RNAseq | ICL7 RNAi at T50 | ERS6679033 | 100914984 | 96275495 | 95% | 89594828 | 89% |
| ChIPseq | ChIP EAP1 RNAi Input r1 | ERS6678581 | 60818036 | 41351387 | 68% | 43243148 | 71% |
| ChIPseq | ChIP EAP1 RNAi H3K27me3 r1 | ERS6678582 | 41084402 | 37545787 | 91% | 38463595 | 94% |
| ChIPseq | ChIP EAP1 RNAi H3K4me3 r1 | ERS6678583 | 39261878 | 31764447 | 81% | 32398850 | 83% |
| ChIPseq | ChIP EAP1 RNAi H3K9me3 r1 | ERS6678584 | 34009760 | 26810174 | 79% | 29833617 | 88% |
| ChIPseq | ChIP EAP1 RNAi Input r2 | ERS6678585 | 36796120 | 34368329 | 93% | 35686888 | 97% |
| ChIPseq | ChIP EAP1 RNAi H3K27me3 r2 | ERS6678586 | 23890210 | 21486837 | 90% | 21888560 | 92% |
| ChIPseq | ChIP EAP1 RNAi H3K4me3 r2 | ERS6678587 | 28440186 | 25183721 | 89% | 25440423 | 89% |
| ChIPseq | ChIP EAP1 RNAi H3K9me3 r2 | ERS6678588 | 35067168 | 30981225 | 88% | 33385493 | 95% |
| ChIPseq | ChIP EZL1 RNAi Input r1 | ERS6678589 | 19451890 | 15994317 | 82% | 17436527 | 90% |
| ChIPseq | ChIP EZL1 RNAi H3K27me3 r1 | ERS6678590 | 51942492 | 23672757 | 46% | 23467765 | 45% |
| ChIPseq | ChIP EZL1 RNAi H3K4me3 r1 | ERS6678591 | 20445628 | 16446181 | 80% | 16991552 | 83% |
| ChIPseq | ChIP EZL1 RNAi H3K9me3 r1 | ERS6678592 | 45218752 | 9615263 | 21% | 13111252 | 29% |
| ChIPseq | ChIP EZL1 RNAi Input r2 | ERS6678593 | 23799754 | 20374560 | 86% | 21254811 | 89% |
| ChIPseq | ChIP EZL1 RNAi H3K27me3 r2 | ERS6678594 | 16848144 | 9450292 | 56% | 9836893 | 58% |
| ChIPseq | ChIP EZL1 RNAi H3K4me3 r2 | ERS6678595 | 20332448 | 17911200 | 88% | 18244748 | 90% |
| ChIPseq | ChIP EZL1 RNAi H3K9me3 r2 | ERS6678596 | 78925068 | 6843165 | 9% | 6927395 | 9% |

| | | | | | | | |
|---|---|---|---|---|---|---|---|
| ChIPseq | ChIP PGM RNAi Input r3 | ERS6678597 | 42264448 | 28698336 | 68% | 30240207 | 72% |
| ChIPseq | ChIP PGM RNAi H3K27me3 r3 | ERS6678598 | 23379466 | 11884667 | 51% | 15662806 | 67% |
| ChIPseq | ChIP PGM RNAi H3K4me3 r3 | ERS6678599 | 46885356 | 40811643 | 87% | 41048201 | 88% |
| ChIPseq | ChIP PGM RNAi H3K9me3 r3 | ERS6678600 | 20305568 | 5232653 | 26% | 10017108 | 49% |
| ChIPseq | ChIP PGM RNAi Input r4 | ERS6678601 | 34262094 | 31732022 | 93% | 33430744 | 98% |
| ChIPseq | ChIP PGM RNAi H3K27me3 r4 | ERS6678602 | 23257042 | 17748412 | 76% | 20049600 | 86% |
| ChIPseq | ChIP PGM RNAi H3K4me3 r4 | ERS6678603 | 29640870 | 26943306 | 91% | 27245102 | 92% |
| ChIPseq | ChIP PGM RNAi H3K9me3 r4 | ERS6678604 | 20237056 | 9562921 | 47% | 14841716 | 73% |
| ChIPseq | ChIP PTIWI01 09 RNAi Input r1 | ERS6678605 | 51011914 | 43541890 | 85% | 45556806 | 89% |
| ChIPseq | ChIP PTIWI01 09 RNAi H3K27me3 r1 | ERS6678606 | 29079750 | 25285027 | 87% | 27126862 | 93% |
| ChIPseq | ChIP PTIWI01 09 RNAi H3K4me3 r1 | ERS6678607 | 27837818 | 25613611 | 92% | 26211507 | 94% |
| ChIPseq | ChIP PTIWI01 09 RNAi H3K9me3 r1 | ERS6678608 | 25294946 | 21087579 | 83% | 23504560 | 93% |
| ChIPseq | ChIP PTIWI01 09 RNAi Input r2 | ERS6678609 | 31497902 | 29370872 | 93% | 30609824 | 97% |
| ChIPseq | ChIP PTIWI01 09 RNAi H3K27me3 r2 | ERS6678610 | 28354956 | 24596262 | 87% | 26064649 | 92% |
| ChIPseq | ChIP PTIWI01 09 RNAi H3K4me3 r2 | ERS6678611 | 29999212 | 27626085 | 92% | 28014732 | 93% |
| ChIPseq | ChIP PTIWI01 09 RNAi H3K9me3 r2 | ERS6678612 | 22448082 | 17524330 | 78% | 19357647 | 86% |
| ChIPseq | ChIP RF4 RNAi Input r1 | ERS6678613 | 37756334 | 34530702 | 91% | 36379162 | 96% |
| ChIPseq | ChIP RF4 RNAi H3K27me3 r1 | ERS6678614 | 38755718 | 29051060 | 75% | 32368786 | 84% |
| ChIPseq | ChIP RF4 RNAi H3K4me3 r1 | ERS6678615 | 54348542 | 51099917 | 94% | 51501728 | 95% |
| ChIPseq | ChIP RF4 RNAi H3K9me3 r1 | ERS6678616 | 53641812 | 32512990 | 61% | 45230585 | 84% |
| ChIPseq | ChIP RF4 RNAi Input r2 | ERS6678617 | 48298142 | 45048359 | 93% | 46320835 | 96% |
| ChIPseq | ChIP RF4 RNAi H3K27me3 r2 | ERS6678618 | 39603396 | 28927273 | 73% | 35013111 | 88% |
| ChIPseq | ChIP RF4 RNAi H3K4me3 r2 | ERS6678619 | 57327350 | 53221021 | 93% | 54508396 | 95% |
| ChIPseq | ChIP RF4 RNAi H3K9me3 r2 | ERS6678620 | 37471814 | 21487183 | 57% | 30003513 | 80% |
| sRNAseq | EZL1 RNAi | ERS6677526 | 12403991 | 6977983 | 56% | 10654026 | 86% |
| sRNAseq | EAP1 RNAi | ERS6677527 | 22038891 | 11385359 | 52% | 15456460 | 70% |
| sRNAseq | RF4 RNAi | ERS6677528 | 14441033 | 6187255 | 43% | 8964237 | 62% |
| sRNAseq | SUZ12 like RNAi | ERS6677529 | 19494897 | 9077678 | 47% | 13261892 | 68% |

**Supplementary Table 2: Sequencing data and mapping statistics**

DNA-seq, mRNA-seq, ChIP-seq and sRNA-seq data used in this study. All the data were deposited to ENA under the accession number PRJE46608. For each sequencing sample the ENA accession is specified followed by the number of reads sequenced and the number of mapped reads on the MAC and MIC reference genomes (See Materials and methods).
